# Using ddRAD-seq phylogeography to test for genetic effects of headwater river capture in suckermouth armored catfish (Loricariidae: *Hypostomus*) from the central Brazilian Shield

**DOI:** 10.1101/2021.04.18.440224

**Authors:** Justin C. Bagley, Pedro De Podestà Uchôa de Aquino, Tomas Hrbek, Sandra Hernandez-Rangel, Francisco Langeani, Guarino R. Colli

## Abstract

River capture is a geological process of potentially great importance in shaping the genetic diversity, distributions, and community composition of freshwater taxa. Using phylogeographic analyses of ddRAD-seq data from suckermouth armored catfish (*Hypostomus* sp. 2) populations, we tested for predicted genetic effects of headwater river capture events in central Brazil, previously supported by geological and community ecological data. We analyzed 227 ddRAD tags (3829 SNP loci) across 42 samples. Molecular results strongly supported six *Hypostomus* genetic clusters/lineages, with the deepest divergence ∼1.25 million years ago in the early Pleistocene between a clade from the Upper Paraná and Upper São Francisco river basins versus all other lineages. Consistent with the ‘Paraná Capture Hypothesis’, several lines of evidence supported mid-Pleistocene colonization and vicariant isolation of *Hypostomus* populations from an ancestral Upper Paraná population, including: (1) significant phylogeographic structure, with predicted phylogenetic patterns, (2) higher Paraná lineage diversity, (3) ancestral geographic locations reconstructed in the Paraná basin, and (4) non-random interdrainage dispersal and vicariance events, indicating river captures primarily into the Tocantins and Upper São Francisco basins *c.* ∼220,000–145,500 years ago. Phylogeographic inference was complicated by lack of lineage monophyly across loci and lineages distributed in multiple basins, the latter of which lent support to the non-mutually exclusive ‘Frequent Interdrainage Dispersal Hypothesis’. However, species tree and demographic modeling results suggested these were artefacts of incomplete sorting of alleles in large ancestral populations over a geologically recent timeframe of divergence. Qualitative and quantitative sensitivity analyses demonstrated that our downstream genetic results were robust to effects of varying ddRAD-seq assembly parameters, which heavily influenced the number of output loci. We predict that codistributed freshwater taxa in Central Brazil may not exhibit phylogeographic patterns similar to *Hypostomus* sp. 2 due to complex patterns of superimposed river capture events, or if smaller ancestral population sizes have allowed more complete lineage sorting in other taxa.

## 1. Introduction

The distribution of freshwater organisms across multiple drainage basins has long intrigued biogeographers and provided natural systems for testing and developing hypotheses of dispersal and vicariance, extinction, and taxon cycles (Myers, 1938; Darlington, 1957; Rosen, 1978; Swift et al., 1985; Mayden, 1988; Banarescu, 1992; Avise, 2000; Hoagstrom et al., 2014). Obligate freshwater organisms are isolated in ‘islands’ of hydrological networks separated by drainage divides (elevational barriers) and inhospitable terrestrial habitats, and by marine environments between river mouths (Unmack, 2001). Thus, geographical distributions of freshwater species spanning multiple drainage basins may arise via fragmentation (vicariance) of a wider-ranging ancestral lineage, or interdrainage dispersal (e.g. Rosen, 1978; Mayden, 1988; Craw et al., 2007). One mechanism of vicariance and dispersal is the displacement of river sections between adjacent drainages, or ‘river capture’ (Swift et al., 1985; Bishop, 1995; Unmack, 2001; Smith and Bermingham, 2005). However, other geomorphological mechanisms could drive interdrainage dispersal, including ‘wet connections’ created by episodic flooding at lakes, swamps, or valleys, or following the migration of a drainage divide (Craw et al., 2007, 2008).

The past 20 years have witnessed a surge of interest in inferring the coevolution of geological and tectonic settings using freshwater species in historical biogeography, particularly through testing hypotheses of vicariance and dispersal with molecular data (e.g. Waters et al., 2001, 2006, 2020; Craw et al., 2007, 2008; Burridge et al., 2006, 2008; Unmack et al., 2012). According to Bishop (1995) and Waters and Craw (2006), the best approach in making such inferences is to assess geological and biological evidence for river capture or reversal independently. Perhaps the strongest biogeographic evidence for these processes thus far comes from using independent geological evidence on the date and scale of river capture or reversal events (ancient versus modern drainage boundaries) to calibrate molecular divergences of freshwater species in New Zealand (Waters et al., 2001, 2007, 2020; Craw et al., 2007, 2008; Burridge et al., 2006, 2008). These studies suggest that river capture has played a major role in the diversification of New Zealand freshwater fishes (Waters et al., 2001; Burridge et al., 2006, 2008). Moreover, for areas in which geological or ecological evidence supports a hypothesis of river capture events, these New Zealand studies also suggest several testable phylogeographic predictions. We expect that river capture (1) isolates populations previously occupying the same drainage by vicariance, causing phylogenetic divergence of ‘captured’ and ‘source’ populations; (2) yields captured-population alleles sister to, or nested within, clades of source population alleles in gene trees or species trees; and (3) causes simultaneous dispersal/vicariance, leaving signatures of gene flow or spatial–demographic expansion (e.g. lower allele frequencies, tip-positioned haplotypes) in captured populations. Still, genetic signatures of population expansion events could be overwritten by multiple events or large changes in effective population size, *N*_e_.

Phylogeography permits testing the above predictions and inferring spatiotemporal patterns of past connections between drainage basins, whether geological calibrations are available or not (e.g. Bermingham and Martin, 1998; Smith and Bermingham, 2005), and with or without established molecular clocks (e.g. Page, 1989). Yet when relevant geological data are lacking, the challenge becomes distinguishing between processes that could yield the same genetic or distributional patterns as river capture, including migration across flooded drainage divides or marine dispersal (Bishop, 1995; Waters and Craw, 2006). Fortunately, freshwater fishes from headwater streams in high-relief areas provide ideal systems for testing hypotheses of river capture events, because they are not subject to confounding effects of marine dispersal. Indeed, ‘primary’ freshwater fishes lack physiological adaptations for past or on-going dispersal through marine environments (Myers, 1938). Marine dispersal is also physically impossible for headwater fish populations isolated in upland habitats.

Another challenge in testing or inferring interdrainage dispersal and gene flow due to river capture is that drawing on a limited number of loci, or only mitochondrial DNA (mtDNA), can be problematic due to low phylogenetic signal, introgression, large population sizes, or the recent timing of river capture events (e.g. Waters et al., 2001; Burridge et al., 2006, 2008). Advances in next-generation sequencing (NGS) offer the hope of overcoming these issues by simultaneously genotyping non-model organisms at hundreds to thousands of loci from throughout the genome (e.g. Andrews et al., 2016). Modeling analyses of genome-scale data allow controlling for coalescent and mutational stochasticity during phylogeographic inference and deriving estimates of population size, gene flow, and divergence times with narrower confidence intervals than single-locus estimates (e.g. Edwards and Beerli, 2000; Knowles, 2009; Gronau et al., 2011). Using many variable nuclear markers can also overcome the inability of small numbers of nuclear genes to infer recent events (e.g. Zink and Barrowclough, 2008).

Neotropical cis-Andean South America provides a geologically and ecologically complex natural laboratory for studying the influences of drainage evolution and river capture on genetic diversity and community assembly, particularly within its crystalline shield highlands (Lima and Ribeiro, 2011; Aquino and Colli, 2017). Drainage rearrangements via headwater river capture are thought to have played an important role shaping primary freshwater fish species diversity and distributions in this region (Ribeiro, 2006; Albert and Carvalho, 2011; Lima and Ribeiro, 2011). Characteristics of headwater streams above 1200 m elevation in the central Brazilian Shield create a particularly ideal system for testing hypotheses of river capture at the triple drainage divide where the Upper Paraná, Upper Araguaia–Tocantins, and Upper São Francisco rivers meet (Aquino, 2013; Aquino and Colli, 2017). In this area, the Trans Brazilian Lineament passes through the Araguaia–Tocantins basin (Saadi, 1993), and tectonic uplift along this fault causes adjacent basins to be captured into the Tocantins (reviewed by Aquino and Colli, 2017).

In this study, we use ddRAD-seq (Peterson et al., 2012) to infer the phylogeographic history of a suckermouth armored catfish (Loricariidae: Hypostominae) lineage, ‘*Hypostomus* sp. 2’ (Aquino et al., 2009; Aquino, 2013), and test for predicted genetic effects of river capture events in the Paraná–Tocantins–São Francisco headwater river system in central Brazil. A recent DNA barcoding study showed that *Hypostomus* sp. 2 is a genetically diverse lineage or species complex containing multiple Barcode Index Numbers, or operational taxonomic units (OTUs), that is sister to a distinct candidate species ‘*Hypostomus* sp. 1’ forming a single OTU (BOLD:ADR4025; Bagley et al., 2019). Aquino and Colli (2017) conducted a phylogenetic community ecology study testing whether species richness and phylogenetic alpha and beta diversity in the Paraná–Tocantins–São Francisco headwater river system were consistent with the direction and frequency of river capture predicted from the geological setting, including higher relative richness in the capturing Tocantins basin. We set out to test specific genetic predictions (summarized in Table 1) derived from two main hypotheses in Aquino and Colli (2017): (1) the ‘Paraná Capture Hypothesis’ stating that Upper Tocantins and Upper São Francisco headwater fish communities were recently colonized from the Upper Paraná basin via headwater captures, based on their lower species richness and phylogenetic diversity values; and (2) the ‘Frequent Interdrainage Dispersal Hypothesis’, which postulates based on random phylogenetic beta diversity among basins that geomorphological processes have caused frequent, recent dispersal events across drainages, which would result in shared alleles or lineages among basins, likely with detectable signatures of gene flow. We test these hypotheses using multiple methods, including inference of population structure, phylogenomics and lineage sorting, coalescent-based estimates of demographic parameters, and phylogeographic spatial diffusion models and randomization tests (Table 1).

**Table 1.**
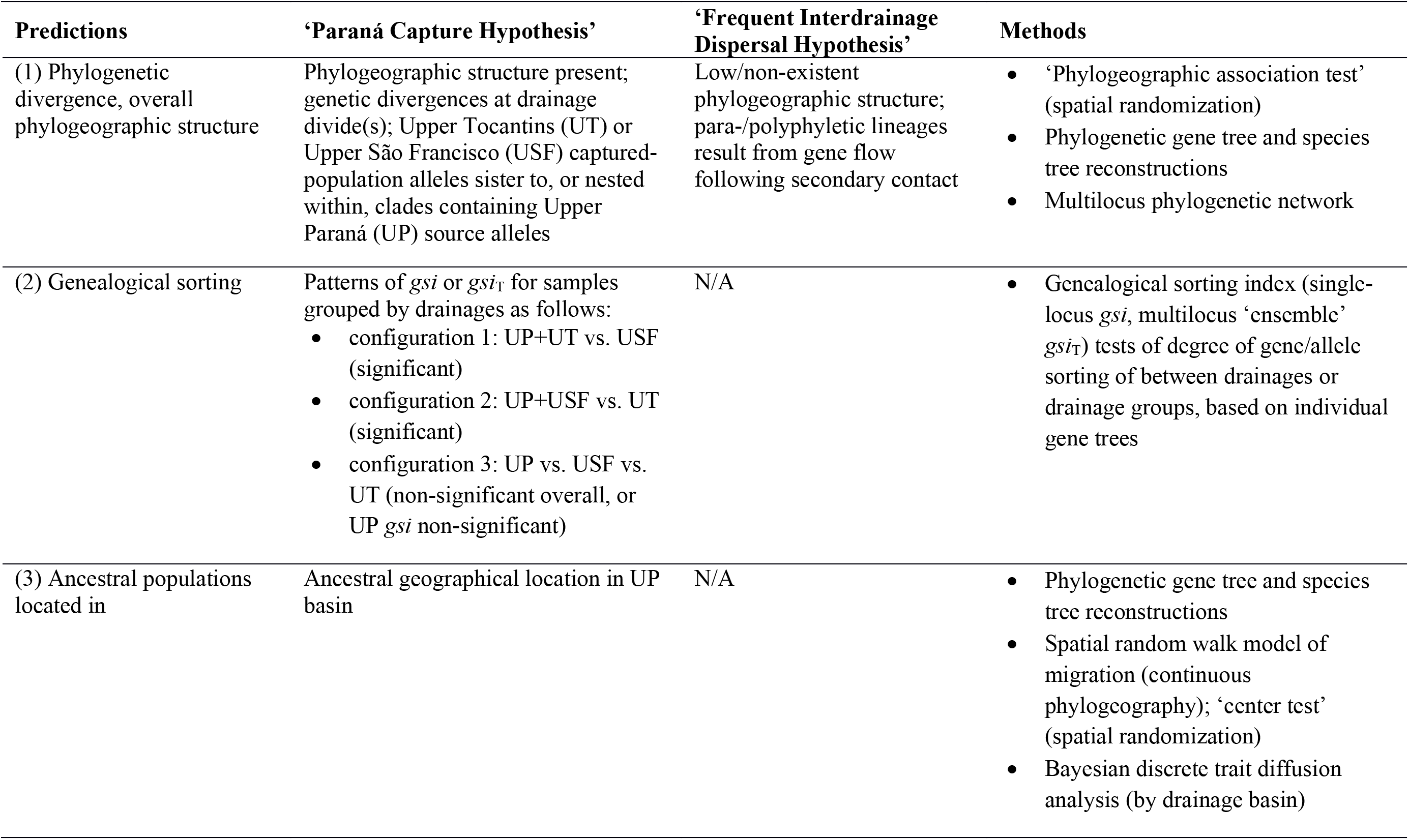
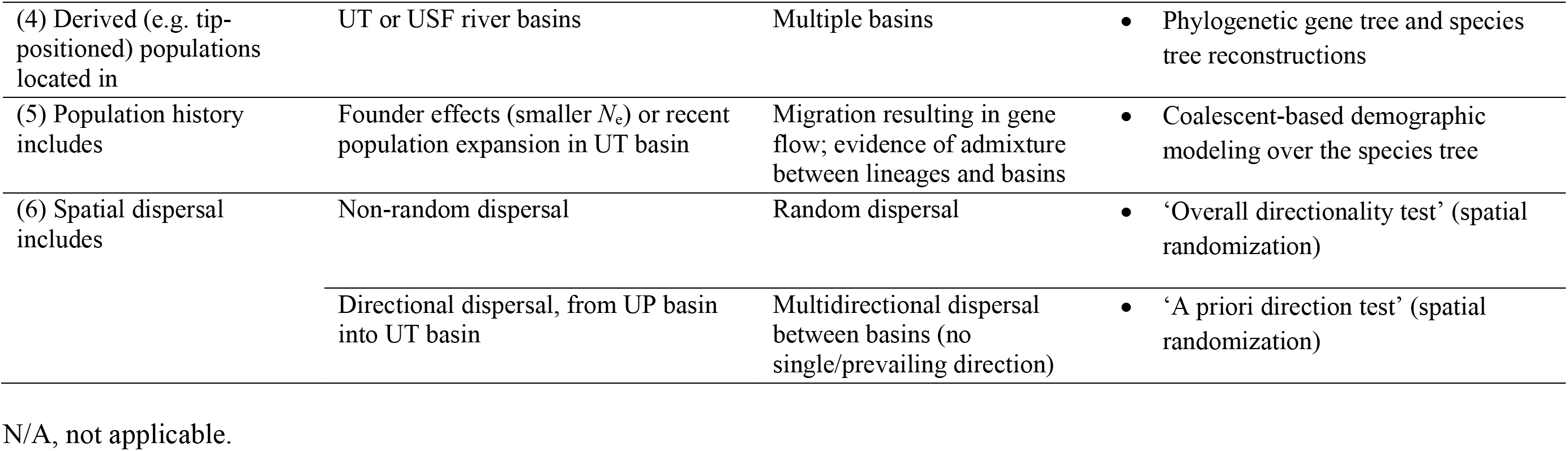
Summary of predicted effects of river capture on fish species phylogeographic history in the study area, under two hypotheses advanced by Aquino and Colli (2017), and overview of methods used to test them.

## 2. Material and methods

### 2.1. Study area and sampling

Permission to undertake field collections and transport material within Brazil was obtained through SISBIO (Sistema de Autorização e Informação em Biodiversidade) permits #48145-1 to JCB and #48111-4 to FLN. Fish were collected from rivers and streams in the study area between March and September of 2015, using 3 × 1 m seines with 2 mm mesh diameter. Specimens or tissue samples (fin clips or muscle plugs) were preserved in 99% ethanol in the field and stored in the laboratory at room temperature. Voucher specimens were preserved in 10% formalin and deposited in the Coleção Ictiológica da Universidade de Brasília (CIUnB). Additional tissue samples of ingroup individuals were obtained from CIUnB collections made under SISBIO permit #42573-1. We sequenced two individuals of the hypostomine candidate species ‘*Hypostomus* sp. 1’, which is the sister species to *Hypostomus* sp. 2 (Aquino et al., 2009; Aquino, 2013; Bagley et al., 2019), as outgroups. Our final dataset encompassed 40 *Hypostomus* sp. 2 samples from 13 collection localities plus two outgroup samples (Figure 1a; Table S1).

**Fig. 1.**
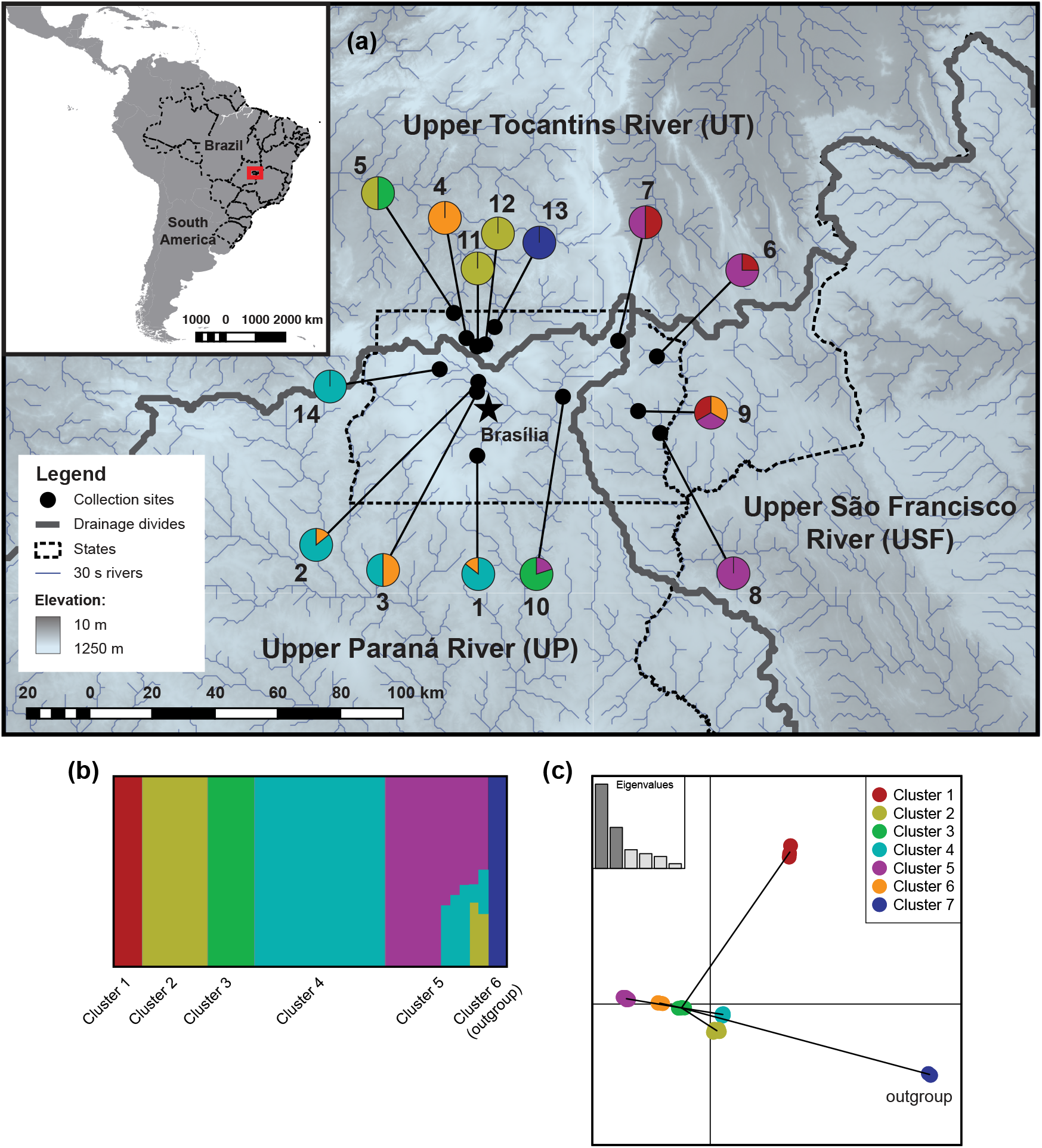
Map of sampling localities and patterns of genetic structure of *Hypostomus* sp. 2 and outgroup individuals inferred using SNPs from 227 ddRAD tags. (a) Sampling localities (dots) are shown plotted over the regional hydrological network, river basin boundaries (drainage divides), and a 250 m digital elevation map (DEM) of the study area (red box, inset). Locality numbers are plotted beside pie charts for each site showing the proportion of individuals belonging to each genetic cluster in panel (c). (b) Barplot of *Q* values from the best model (*K* = 6) obtained during inference of population structure and admixture in fastSTRUCTURE v1.0 (Raj et al., 2014). (c) Scatterplot of relationships between seven genetic clusters (6 ingroup clusters, 1 outgroup cluster) identified during discriminant analysis of principal components (DAPC) in adegenet v2.0.1 (Jombart, 2008; Jombart et al., 2010), as captured by the first two discriminant axes; clusters are connected by a minimum spanning tree and colored according to clusters shown in (b), except for novel DAPC cluster 5.

### 2.2. DNA extraction, library preparation, and sequencing

We extracted whole genomic DNA from tissue samples using Invitrogen PureLink Genomic Purification kits. In the final extraction step, DNA was eluted from spin columns using two consecutive steps, each consisting of a 2 min incubation in 75–100 µL of elution buffer followed by a 1 min spin at 14,000×*g*. We confirmed the presence of high molecular weight genomic DNA on 1% agarose gels and then quantified sample DNA concentration and purity using a NanoDrop 2000c Spectrophotometer (Thermo Scientific). Samples with < 10 ng/µL of DNA were dried down for 30 min on a vacuum centrifuge and resuspended in ∼50 µL of deionized, ultra-purified water. Preparation of multiplexed ddRAD-seq libraries for each barcoded individual sample (Table S2); equimolar pooling of sample libraries; and final library concentration, size selection, and purification followed steps outlined in Appendix S1A. The final pooled library was diluted to an appropriate concentration and then subjected to Ion Sphere Particle (ISP) emulsion PCR amplification using an Ion One Touch 2 and an Ion PI Template OT2 200 Kit (Life Technologies). We sequenced the final library in two runs on an Ion Torrent Personal Genome Machine (Life Technologies) using an Ion 318 Chip Kit.

### 2.3. De novo assembly and SNP processing

We converted raw sequence reads from BAM format to FASTQ format using SAMtools v1.3.1 (Li et al., 2009). We then used PyRAD v3.0.66 (Eaton, 2014) to de-multiplex the data and assemble ddRAD tags de novo, because it has been shown to yield more output loci than similar assembly pipelines (e.g. Jungwirth, 2017). We assigned sequences to individuals, removed reads with errors in the individual-specific barcodes, and required reads to have Phred quality scores of at least 20. We kept trimmed reads longer than 330 bp. Given minimum read depth of coverage of ≥ 5–10× is desirable for genotyping-by-sequencing (Kenny et al., 2011; Peterson et al., 2012), we clustered reads into stacks at 88% sequence similarity and then called SNPs in PyRAD while enforcing a minimum stack depth of five and aiming for ∼10× coverage. We capped the amount of missing data in each ddRAD-seq locus in our final assembly at 25% by requiring minimum taxon-coverage of 30 individuals. Sequences were clustered within and across individuals by similarity using VSEARCH (Rognes et al., 2016) and aligned using MUSCLE (Edgar, 2004).

### 2.4. Population structure and admixture

We estimated the number of distinct genetic clusters (*K*) among our samples, and admixture between clusters, in a variational Bayesian framework in fastSTRUCTURE v1.0 (Raj et al., 2014). We automated independent runs of the program with the simple admixture model on *K* = 1–15 clusters using the ‘fastSTRUCTURE’ function in PIrANHA v0.4-alpha-4 (Bagley, 2020). We estimated the best *K* value according to the model with the fewest components accounting for 99.99% of the ancestry in the sample. We interpreted the best*-K* model as the most likely number of genetic clusters, assumed to be at Hardy–Weinberg equilibrium. We visualized best-*K* run results in DISTRUCT v1.1 (Rosenberg, 2004) by averaging the variational posterior distribution over admixture proportions (*Q*). We also estimated the number of distinct genetic clusters among samples using discriminant analysis of principal components (DAPC; Jombart et al., 2010) in the adegenet v2.0.1 package (Jombart, 2008; Jombart and Ahmed, 2011) for the R environment (R Core Team, 2018). During DAPC, we retained the minimum number of PCs (*N* = 5) capturing ≥ 75% of data variance and we conducted *k*-means clustering on the PCs to determine the number of distinct clusters, as identified by a leveling off of Bayesian information criterion (BIC) scores plotted against *K*. The best *K* was specified in the DAPC prior, and we retained six discriminant axes and eigenvalues from discriminant analysis before plotting results.

### 2.5. Phylogenomic gene tree and species tree analyses

We used PartitionFinder v1.1.1 (Lanfear et al., 2012) to select appropriate partitioning schemes and models of sequence evolution for each DNA subset, or ddRAD tag locus, in our final alignment. We ran PartitionFinder on all subsets using relaxed hierarchical clustering (Lanfear et al., 2014). Across 56 substitution models, PartitionFinder estimated subset-specific phylogenetic branch lengths (branchlengths = linked), and derived maximum-likelihood (ML) estimates for the base frequencies, proportion of invariant sites (I), and the gamma shape distribution (Γ). The ‘best-fit’ partitioning scheme was determined based on BIC values.

We empirically tested for patterns of phylogenetic gene tree and species tree divergences predicted under our hypotheses (Table 1) using three approaches. First, we conducted ML phylogenetic analyses on our final *Hypostomus* alignment in RAXML v8.2.8 (Stamatakis, 2014). In each of three replicate runs, we partitioned the data supermatrix according to the scheme and models selected in PartitionFinder, and we conducted a simultaneous ML tree and bootstrap (500 pseudoreplicates) analysis. We also inferred gene trees through a partitioned Bayesian analysis in MrBayes v3.2 (Ronquist et al., 2012) run for 10 million generations (sampling every 1000) using the same partitions and models used in RAXML. We used the ‘MrBayesPostProc’ function in PIrANHA (Bagley 2020) to (1) discard the first 25% of trees as ‘burn-in’, (2) summarize posterior distributions, and ensure convergence and adequate effective sample size (ESS), and (3) compute a 50% majority-rule consensus tree annotated with Bayesian posterior probabilities (BPPs) along nodes in MrBayes. We considered nodes with ≥ 0.95 BPP to be strongly supported.

Second, we estimated species trees using two approaches while specifying the genetic clusters identified in our population structure and admixture analyses as ‘species’. We estimated a species tree using the multispecies coalescent SNP model implemented in the SNAPP (Bryant et al., 2012) module of BEAST v2.4.5 (Bouckaert et al., 2014). SNAPP runs inferred the species tree from biallelic SNPs while integrating over all possible gene trees and estimating lineage population sizes. We experimented with different rate priors (uniform, gamma) and ran each SNAPP model for 10 replicate runs each 1 million generations in length (sampling every 1000), while fixing the forward and reverse mutation rates to their empirical values. Convergence was assessed in Tracer v1.6 (Rambaut et al., 2013) based on parameter stationarity and ESS scores greater than 200. We also inferred species tree relationships using SVDquartets (Chifman and Kubatko, 2014) as implemented in PAUP* v4.0a147 (Swofford, 2003). SVDquartets samples quartets of sequences from each individual and infers an unrooted phylogeny, and then uses all sampled quartets to conduct species tree or gene tree inference using a multispecies coalescent model (Chifman and Kubatko, 2014). We exhaustively sampled quartets and used them to infer a species tree, then a gene tree, and we estimated nodal support using 500 bootstrap pseudoreplicates. Our sequence alignments and phylogenetic results are archived in Zenodo (doi:10.5281/zenodo.16XXXX). Third, the above phylogenetic results were compared with a multilocus phylogenetic network (Table 1; additional details in Appendix S1A).

We empirically tested for predicted patterns (Table 1) in the degree and distribution of exclusive ancestry of tip alleles (individuals) among drainage basins, as quantified by calculating the genealogical sorting index (*gsi*; Cummings et al., 2008). We calculated *gsi* for tips when assigned to groups while assuming different drainage basin or captured/capturing basin configurations, using RAxML gene trees for each ddRAD tag locus. We also calculated the multilocus, ‘ensemble’ *gsi* statistic (*gsi*_T_) as the weighted sum of *gsi* across gene trees. Values of *gsi* range from 0 to 1, indicating increasing lineage sorting from non-exclusive ancestry to complete monophyly. Analyses were run on three tip configurations in the genealogicalSorting v0.92 R package (Cummings et al., 2008) while using the ‘multiTreeAnalysis’ function and testing significance with 10,000 permutations. Two configurations that we tested represented captures of Upper Paraná tributaries into adjacent basins: (1) Upper Paraná + Upper Tocantins tips versus Upper São Francisco tips, and (2) Upper Paraná + Upper São Francisco tips versus Upper Tocantins tips (Table 1). A third configuration consisted of (3) tips from each of the three drainage basins (Table 1). Under a Paraná Capture Hypothesis scenario, we expected significant *gsi*_T_ across groups for configurations 1 and 2; however, we expected non-significant *gsi*_T_ for the Upper Paraná group in configuration 3 analyses. We also predicted lower *gsi* values for groups including Upper Paraná tips, reflecting admixture of specific parts of the genome during historical to recent drainage connections with the Paraná basin. We expected the last two predictions above to be particularly likely to hold if river capture and divergence have been recent in time, constraining the timeframe for allele sorting (e.g. Avise 2000). Given incomplete lineage sorting and gene flow can yield similar genetic patterns (Rieseberg et al., 1999; Funk and Omland, 2003; Sousa and Hey, 2013), and demographic histories involving multiple migration events could be complex, we refrained from making any a priori predictions about genealogical sorting under a Frequent Interdrainage Dispersal Hypothesis scenario (Table 1).

### 2.6. Demographic history and migration

We estimated divergence times, population sizes, and migration rates among lineages in the species tree using SNAPP (discussed above) and G-PhoCS v1.2.3 (Generalized Phylogenetic Coalescent Sampler; Gronau et al., 2011), while assigning individuals to lineages formed by genetic clusters identified by DAPC. Our SNAPP runs inferred population sizes but did not incorporate migration or yield interpretable divergence time estimates. In contrast, G-PhoCS is a Bayesian coalescent genealogy sampler that estimates divergence times, allows gene flow between populations, and integrates over possible phases of unphased diploid genotypes (Gronau et al., 2011). By default, G-PhoCS assumes constant mutation and no intralocus recombination. We used results to infer demographic history and also test the Frequent Interdrainage Hypothesis prediction of gene flow between lineages or basins (Table 1).

We supplied G-PhoCS with the SVDquartets species tree and ran replicate runs for 1 million generations (sampling every 100) on the full SNP dataset for a no-migration model (*m_1_*) and a low-migration model (*m_2_*). As estimating many migration parameters is computationally difficult and could potentially bias divergence times (Gronau et al., 2011), we conducted separate *m_2_* runs estimating migration rates between each pair of genetic clusters treated as ‘species’. We set the population mutation rate parameter (*θ* = 4*N*_e_*µ*, where *µ* = mutation rate per generation) and divergence time parameter (*τ*; coalescent units, in generations; note *τ* = *Tµ*, where *T* = time in millions of years ago) to gamma priors of ∼G(2, 1000) determined by pilot runs using a range of prior values. In *m_2_* runs, the gamma prior for the migration rate parameter (*m_sx_*) was set to ∼G(1, 10). We converted divergence time estimates to absolute time using a per-lineage mean rate calibration of 0.29% Myr^−1^ derived from BEAST divergence time analyses of a multilocus dataset in a recent phylogenetic study of *Hypostomus* (Silva et al., 2016). This mean rate closely matches other teleost fish genome evolution rate estimates, e.g. the ‘fish’ SNP evolution rate of 0.19% Myr^−1^ estimated from the medaka genome (Setiamarga et al., 2009). We converted G-PhoCS *τ* values to absolute time (*T*) using the relationship *τ = Tµ*/*g* (where *g* = generation time in years, set to 1.0 due to uncertainty for our taxa). Raw *θ_x_* values from SNAPP and G-PhoCS were converted to effective population sizes (*N*_e_). Migration parameters were converted to migration per generation using the value of *θ_x_* for the recipient population (*M_sx_* = *m_sx_* × *θ_x_*/4; per generation proportion of individuals in population *x* that arrived by migration from population *s*).

### 2.7. Phylogeographic analyses

We inferred the geographical origin of *Hypostomus* sp. 2 using the ML random walk method implemented in PhyloMapper v1b1 (Lemmon and Lemmon, 2008). We supplied PhyloMapper with the multispecies coalescent gene tree from SVDquartets and fixed the ingroup node age to its mean age estimate from G-PhoCS. We used nonparametric rate smoothing (Sanderson, 1997) to ultrametricize the topology, smoothing over the clade whose ancestral area was being estimated. To produce a distribution of ancestor locations and ensure that a global optimum was reached, we conducted 100 independent runs in PhyloMapper, each of which inferred the ancestral location of the ingroup using 100 ML search replicates and the default optimization criteria. We also used PhyloMapper to conduct four randomization tests of phylogeographic hypotheses. First, we tested for a significant correlation between ingroup geographic distances and phylogenetic distances (‘phylogeographic association test’) by randomizing geographical coordinates of localities across the tree tips 10,000 times and using the scaled dispersal parameter (*ψ*) as a test statistic (Lemmon and Lemmon, 2008). Second, we tested the null hypothesis that the ingroup ancestor was located at the center of the sampled localities (‘center test’), as expected under equilibrium conditions with no recent population expansions (Lemmon and Lemmon, 2008). This test compared two separate optimizations of the model using a likelihood ratio test, one using the estimated center of the locality coordinates and the other using the ML ancestral location estimate. Third, we tested the null hypothesis that the average direction of dispersal was random based on 10,000 randomizations of the mean displacement parameter (‘overall directionality test’). Using similar randomizations, we tested whether individuals tended to historically migrate north/northwest into the Tocantins basin, or east into the São Francisco basin (‘a priori direction test’). Statistical tests were considered significant at the α = 0.05 level.

To complement our PhyloMapper analyses, we estimated the ancestral area location and inferred spatial patterns of diffusion migration of *Hypostomus* sp. 2 lineages using the discrete phylogeographic model (Lemey et al., 2009) in BEAST. Whereas PhyloMapper uses a fixed gene tree estimate, discrete trait analysis in BEAST estimates phylogeographic parameters in a Bayesian model without assuming a known tree topology, and also simultaneously estimates divergence times. In each of three replicate runs using drainage basin as the discrete ‘location’ trait value for each individual, we ran BEAST for 100 million MCMC generations (sampling every 10,000). Again, we partitioned the data according to the data subsets identified in PartitionFinder. BEAST runs linked tree models and DNA partitions, applied a relaxed uncorrelated lognormal clock to the DNA partitions and a strict clock to the locations, and used a Coalescent Bayesian Skyline tree prior. For simplicity, and to increase chances of convergence, we applied a GTR+Γ+I site model (Tavaré, 1986) to each DNA partition and a gamma site model to the locations. We ensured adequate run length and convergence using Tracer and then visualized post-burn-in results in DensiTree (Bouckaert, 2010).

### 2.8. Effects of varying ddRAD-seq assembly parameters on downstream genetic analyses

We explored the effects of varying PyRAD assembly parameters on our downstream genetic analyses by (1) producing a large test set of PyRAD assemblies reflecting a range of parameter settings, (2) *qualitatively* comparing assembly characteristics (numbers of ddRAD tag loci and SNPs), and gene tree and admixture group inferences derived from the assemblies, and (3) *quantitatively* evaluating sensitivity of outcomes using variance-based sensitivity analyses. A test set of 280 different PyRAD assemblies was generated by varying five parameters (mC, minimum number of samples in a final locus; SH, max. individuals with a shared heterozygous site; pO, Phred score offset; mD, minimum depth of coverage; cP, clustering percentage threshold) across two- or three-value ranges within ±14–30% of the assembly parameters used in our empirical analyses above. Our sampling of parameter values across test assembly runs approximated a near-random sample from a multidimensional distribution. For each assembly, we estimated a gene tree in RAXML using a concatenated, partitioned analysis under the GTR+Γ model, and we estimated and plotted results from fastSTRUCTURE models over *K* = 1–15, to match our regular analyses. We used fastSTRUCTURE instead of DAPC because it could more readily be incorporated into our custom scripts than DAPC, which requires visual inspection of output. Results were evaluated and graphically plotted using custom R scripts. Gene trees from each assembly were plotted and compared in multivariate treespace using principal coordinates analysis (PCoA) on Robinson–Foulds distances (*d*_RF_) reflecting topological differences, and then geodesic distances (*d_G_*) reflecting branch length variation. Assuming our choice of PyRAD assembly settings did not unduly biased our empirical results, we expected treespace to be rather uniform, with topological variance approximately normally distributed around *d*_RF_ and *d_G_* values for the gene tree from our original empirical results, and that best *K* estimates across assemblies would be similar to our empirical fastSTRUCTURE results (*K* = 6).

We conducted quantitative variance-based sensitivity analysis using Sobol’ sensitivity indices (Sobol’, 1990; Saltelli et al., 2008) estimated using the Latin hypercube space method of Tissot and Prieur (2015) as implemented in the R package ‘sensitivity’ (Pujol et al., 2017). We estimated Sobol’ first-order and second-order indices of sensitivity of PyRAD parameters based on a vector of four assembly outcome variables: (1) mean best-tree *d*_RF_ from all other best trees; (2) mean best-tree *d*_G_; (3) best *K*; and (4) number of ddRAD tag loci. For each model input variable *X_i_* that was varied, Sobol’ first-order indices (*S_Xi_*) reflect the contribution to total output variance of the decomposed variance attributable to *X_i_* (Saltelli et al., 2008). Likewise, Sobol’ second-order indices (*S_Xi_*_,*Xj*_) reflect the effect of varying two parameters, *X_i_* and *X_j_*, simultaneously, beyond the effect of the variation of each individual parameter, akin to interaction effects between variables (Saltelli et al., 2008). Discrete variables, and continuous variables identified as non-normally distributed using Shapiro–Wilk tests (α *=* 0.05), were log-transformed prior to sensitivity analyses.

## 3. Results

### 3.1. De novo assembly and SNP processing

Our Ion Torrent sequencing experiment yielded 9.6 million raw, single-end reads, for a total of 2.42 billion bp. After de-multiplexing and data filtering, clustering at 88% similarity yielded a total of 156,088 clusters, with on average 3716.4 clusters per individual. Filtering further to a minimum depth of coverage of five in PyRAD yielded a final set of 27,242 clusters with a mean coverage depth of 20.5× (Table 2). On average, 637 ddRAD tag sequences from each individual (range = 513–1094) passed the paralogy-filtering step (Table 2). On average, only 10.8 tag sequences (range = 4–38; mean = 1.63%, range = 0.7–3.8%) were flagged as paralogs within individuals and excluded, indicating that paralogy was not a prevalent issue in our dataset (additional details in Appendix S1). Among consensus sequences, mean heterozygosity per individual was approximately 4-fold greater than the mean error rate, providing an excellent basis for SNP calling (Table S3). The raw number of SNPs was 4197 loci. After filtering the assembly to have no more than ∼25% missing data for any individual or locus, we arrived at a final alignment consisting of 227 ddRAD sequences, each 330–374 bp in length, for a total alignment length of 78,265 bp. The final sequence alignment contained 3969 variable sites, of which 3047 (77%) were parsimony informative. The final SNP dataset contained 3829 loci.

**Table 2.**
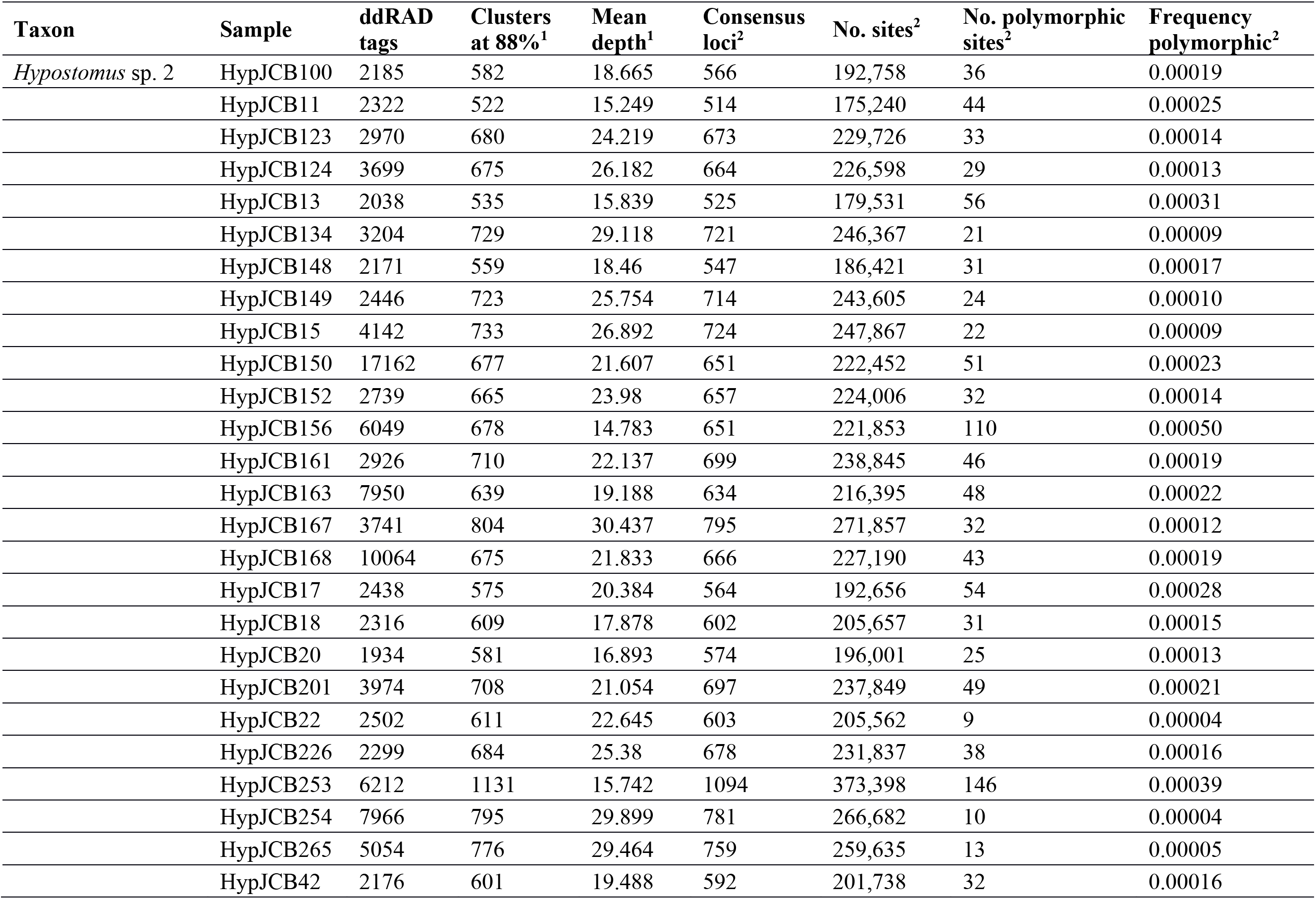
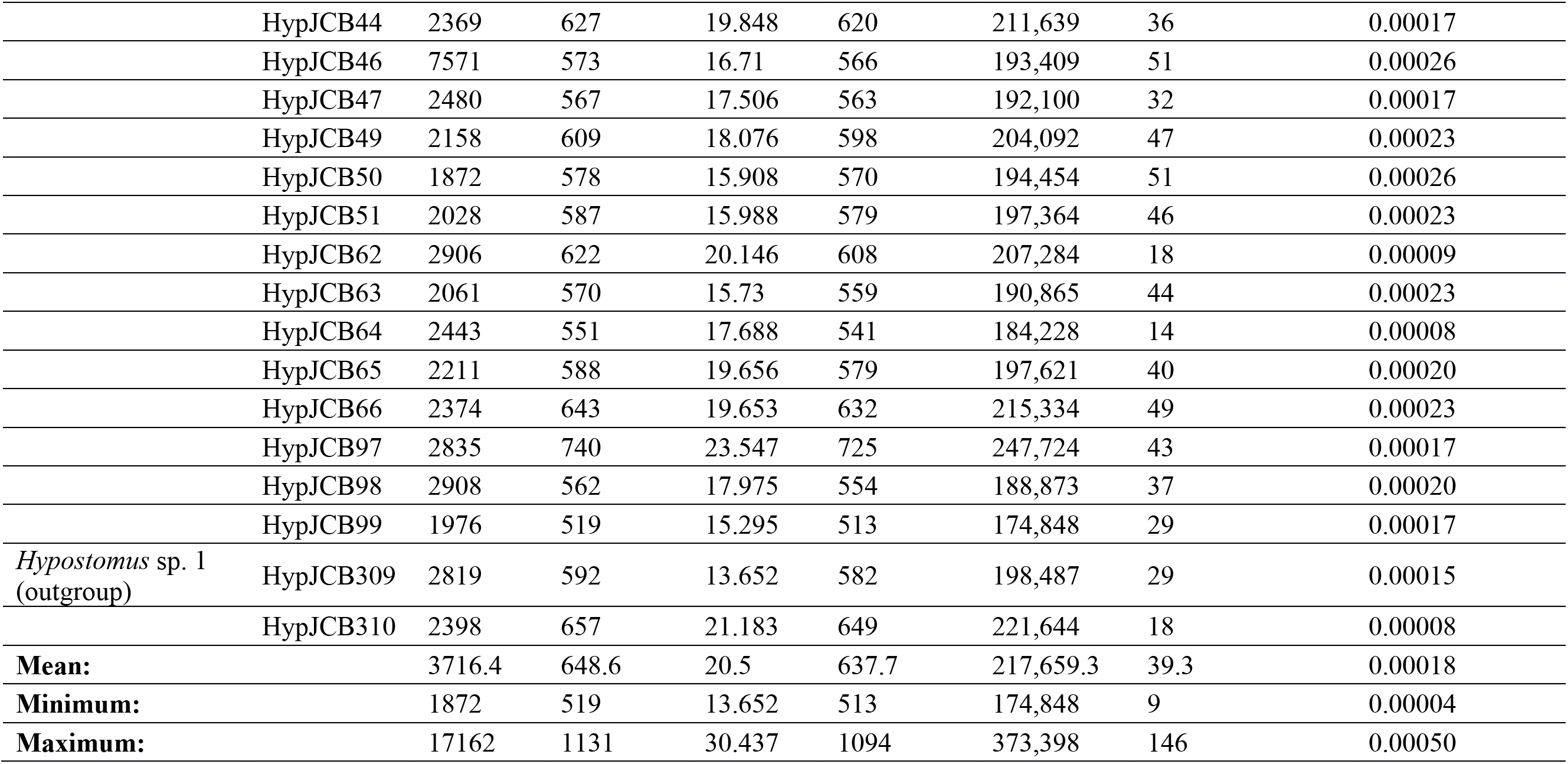
Results of filtering and clustering ddRAD tags from each sample in this study in PyRAD v3.0.66 (Eaton, 2014). Additional sampling information is given in Table S1 of the Supporting Information. ^1^Clusters with more than or equal to the minimum depth of coverage (‘depth’) of five reads. ^2^Summary statistics for number and characteristics of consensus loci that passed additional filtering for a maximum of four sites with Phred quality scores less than 20, as well as paralog filtering. We arrived at the final set of 227 ddRAD-seq loci by subsequently filtering these consensus loci to obtain only loci with no more than ∼25% missing taxa, and additional filtering was conducted on the SNP loci contained in these sequences.

| Taxon | Sample | ddRAD tags | Clusters at 88%<sup>1</sup> | Mean depth<sup>1</sup> | Consensus loci<sup>2</sup> | No. sites<sup>2</sup> | No. polymorphic sites<sup>2</sup> | Frequency polymorphic<sup>2</sup> |
| --- | --- | --- | --- | --- | --- | --- | --- | --- |
| *Hypostomus* sp. 2 | HypJCB100 | 2185 | 582 | 18.665 | 566 | 192,758 | 36 | 0.00019 |
|  | HypJCB11 | 2322 | 522 | 15.249 | 514 | 175,240 | 44 | 0.00025 |
|  | HypJCB123 | 2970 | 680 | 24.219 | 673 | 229,726 | 33 | 0.00014 |
|  | HypJCB124 | 3699 | 675 | 26.182 | 664 | 226,598 | 29 | 0.00013 |
|  | HypJCB13 | 2038 | 535 | 15.839 | 525 | 179,531 | 56 | 0.00031 |
|  | HypJCB134 | 3204 | 729 | 29.118 | 721 | 246,367 | 21 | 0.00009 |
|  | HypJCB148 | 2171 | 559 | 18.46 | 547 | 186,421 | 31 | 0.00017 |
|  | HypJCB149 | 2446 | 723 | 25.754 | 714 | 243,605 | 24 | 0.00010 |
|  | HypJCB15 | 4142 | 733 | 26.892 | 724 | 247,867 | 22 | 0.00009 |
|  | HypJCB150 | 17162 | 677 | 21.607 | 651 | 222,452 | 51 | 0.00023 |
|  | HypJCB152 | 2739 | 665 | 23.98 | 657 | 224,006 | 32 | 0.00014 |
|  | HypJCB156 | 6049 | 678 | 14.783 | 651 | 221,853 | 110 | 0.00050 |
|  | HypJCB161 | 2926 | 710 | 22.137 | 699 | 238,845 | 46 | 0.00019 |
|  | HypJCB163 | 7950 | 639 | 19.188 | 634 | 216,395 | 48 | 0.00022 |
|  | HypJCB167 | 3741 | 804 | 30.437 | 795 | 271,857 | 32 | 0.00012 |
|  | HypJCB168 | 10064 | 675 | 21.833 | 666 | 227,190 | 43 | 0.00019 |
|  | HypJCB17 | 2438 | 575 | 20.384 | 564 | 192,656 | 54 | 0.00028 |
|  | HypJCB18 | 2316 | 609 | 17.878 | 602 | 205,657 | 31 | 0.00015 |
|  | HypJCB20 | 1934 | 581 | 16.893 | 574 | 196,001 | 25 | 0.00013 |
|  | HypJCB201 | 3974 | 708 | 21.054 | 697 | 237,849 | 49 | 0.00021 |
|  | HypJCB22 | 2502 | 611 | 22.645 | 603 | 205,562 | 9 | 0.00004 |
|  | HypJCB226 | 2299 | 684 | 25.38 | 678 | 231,837 | 38 | 0.00016 |
|  | HypJCB253 | 6212 | 1131 | 15.742 | 1094 | 373,398 | 146 | 0.00039 |
|  | HypJCB254 | 7966 | 795 | 29.899 | 781 | 266,682 | 10 | 0.00004 |
|  | HypJCB265 | 5054 | 776 | 29.464 | 759 | 259,635 | 13 | 0.00005 |
|  | HypJCB42 | 2176 | 601 | 19.488 | 592 | 201,738 | 32 | 0.00016 |
|  | HypJCB44 | 2369 | 627 | 19.848 | 620 | 211,639 | 36 | 0.00017 |
|  | HypJCB46 | 7571 | 573 | 16.71 | 566 | 193,409 | 51 | 0.00026 |
|  | HypJCB47 | 2480 | 567 | 17.506 | 563 | 192,100 | 32 | 0.00017 |
|  | HypJCB49 | 2158 | 609 | 18.076 | 598 | 204,092 | 47 | 0.00023 |
|  | HypJCB50 | 1872 | 578 | 15.908 | 570 | 194,454 | 51 | 0.00026 |
|  | HypJCB51 | 2028 | 587 | 15.988 | 579 | 197,364 | 46 | 0.00023 |
|  | HypJCB62 | 2906 | 622 | 20.146 | 608 | 207,284 | 18 | 0.00009 |
|  | HypJCB63 | 2061 | 570 | 15.73 | 559 | 190,865 | 44 | 0.00023 |
|  | HypJCB64 | 2443 | 551 | 17.688 | 541 | 184,228 | 14 | 0.00008 |
|  | HypJCB65 | 2211 | 588 | 19.656 | 579 | 197,621 | 40 | 0.00020 |
|  | HypJCB66 | 2374 | 643 | 19.653 | 632 | 215,334 | 49 | 0.00023 |
|  | HypJCB97 | 2835 | 740 | 23.547 | 725 | 247,724 | 43 | 0.00017 |
|  | HypJCB98 | 2908 | 562 | 17.975 | 554 | 188,873 | 37 | 0.00020 |
|  | HypJCB99 | 1976 | 519 | 15.295 | 513 | 174,848 | 29 | 0.00017 |
| *Hypostomus* sp. 1<br>(outgroup) | HypJCB309 | 2819 | 592 | 13.652 | 582 | 198,487 | 29 | 0.00015 |
|  | HypJCB310 | 2398 | 657 | 21.183 | 649 | 221,644 | 18 | 0.00008 |
| **Mean:** |  | 3716.4 | 648.6 | 20.5 | 637.7 | 217,659.3 | 39.3 | 0.00018 |
| **Minimum:** |  | 1872 | 519 | 13.652 | 513 | 174,848 | 9 | 0.00004 |
| **Maximum:** |  | 17162 | 1131 | 30.437 | 1094 | 373,398 | 146 | 0.00050 |

### 3.2. Population structure and admixture

By comparing runs with different *K* values in fastSTRUCTURE, we identified five distinct genetic clusters among ingroup samples, plus one cluster of outgroup samples (*K* = 6; marginal likelihood = −0.442) (Figure 1b). The genotypes of five individuals within cluster 5 indicated admixture with cluster two (*N* = 3 individuals), or clusters two and four (*N* = 2 individuals). The six discriminant functions used in DAPC captured 82.4% of cumulative data variance. DAPC results supported the same six clusters as fastSTRUCTURE, with the only exception being that fastSTRUCTURE cluster 5 (*N* = 11) was split into two clusters (5 and 6), yielding seven total clusters (*K* = 7) (Figure 1c), a value that matched the inflection point in BIC values (Figure S1). DAPC cluster 3 contained the five putatively admixed individuals from fastSTRUCTURE cluster 5, while DAPC cluster 2 contained the remaining six non-admixed individuals from fastSTRUCTURE cluster 5. We took our DAPC results as the best estimate of genetically distinct clusters among our population structure results because DAPC clusters provided a better match to our phylogenetic results, putatively admixed individuals might represent a distinct population or lineage, and a recent study showed that *k*-means clustering used in DAPC is more accurate than fastSTRUCTURE over a range of conditions (Stift et al., 2019). Hereafter, we refer only to the genetic clusters inferred from DAPC, unless stated otherwise.

### 3.3. Phylogenomic gene tree and species tree analyses

PartitionFinder identified a best-fit scheme partitioning the DNA alignment into 11 partition subsets ranging from 334 bp to 44,131 bp in length and containing up to 128 partitions (groups of ddRAD tag sequences; Table 3). Based on BIC values, the GTR model with an estimated proportion of invariable sites (*I*) and a gamma model of rate heterogeneity (Γ) was selected as the optimal model of DNA sequence evolution for most subsets (Table 3); thus, we specified the GTR+Γ+I model in all partitioned phylogenetic analyses.

**Table 3.** Summary of data partitions and substitution models selected by PartitionFinder v1.1.1 for the ddRAD-seq dataset analyzed in this study. Data subsets correspond to groups of data partitions read into PartitionFinder, i.e. groups of ddRAD tag sequences. bp, nucleotide base pairs.

| Subset | No. subset partitions | Total length (bp) | Best-fit model |
| --- | --- | --- | --- |
| 1 | 23 | 7919 | GTR+$\Gamma$+I |
| 2 | 1 | 336 | GTR+$\Gamma$+I |
| 3 | 1 | 348 | GTR+$\Gamma$+I |
| 4 | 128 | 44,131 | GTR+$\Gamma$+I |
| 5 | 53 | 18,296 | GTR+$\Gamma$+I |
| 6 | 12 | 4119 | GTR+$\Gamma$+I |
| 7 | 3 | 1050 | GTR+$\Gamma$+I |
| 8 | 2 | 695 | GTR+$\Gamma$+I |
| 9 | 2 | 685 | GTR+$\Gamma$+I |
| 10 | 1 | 352 | GTR+$\Gamma$+I |
| 11 | 1 | 334 | GTR+$\Gamma$+I |
| **Total:** | 227 | 78,265 | — |

Partitioned phylogenetic analyses using ML and Bayesian inference analyses strongly supported six distinct monophyletic groups with two or more ingroup samples (clades 1–6 below; BP=100/BPP=1), plus a clade of outgroup samples. The best topology from RAXML had a log likelihood (ln *L*) score of −152055.9024 (Figure 2a). Here, ‘clade 1’ was comprised of three individuals from cluster 1 from localities 6, 7, and 9 in the very upstream-most regions of the Upper Paraná and Upper São Francisco basins. ‘Clade 2’ contained five putatively admixed individuals from fastSTRUCTURE cluster 3 and DAPC cluster 6 from all three drainage basins (Figure 1). ‘Clade 3’ included six putatively non-admixed individuals in cluster 2 from the Upper Tocantins River basin, plus one cluster 3 individual (HypJCB156) from the Upper Paraná. ‘Clade 4’ was distributed in the five eastern-most collection localities in the study area (6–10) and included seven cluster 5 individuals from plus one cluster 3 individual (HypJCB66). ‘Clade 5’ contained the three remaining cluster 3 individuals, each collected from locality 10 in the Upper Paraná basin. Last, ‘clade 6’ included all 14 individuals from cluster 4. Only clades 3 and 6 were endemic to one river drainage—the Upper Tocantins and Upper Paraná rivers, respectively. The partitioned analysis in MrBayes exhibited excellent chain mixing and convergence characteristics (potential scale reduction factor converged on a value of 1.0) and yielded a consensus tree that was identical to the best ML topology, except for placing HypJCB167 sister to HypJCB168 rather than HypJCB201. While BPP support from MrBayes is given along nodes in the ML tree (Figure 2a), we provide the Bayesian consensus topology as supplementary material (Figure S2).

**Fig. 2.**
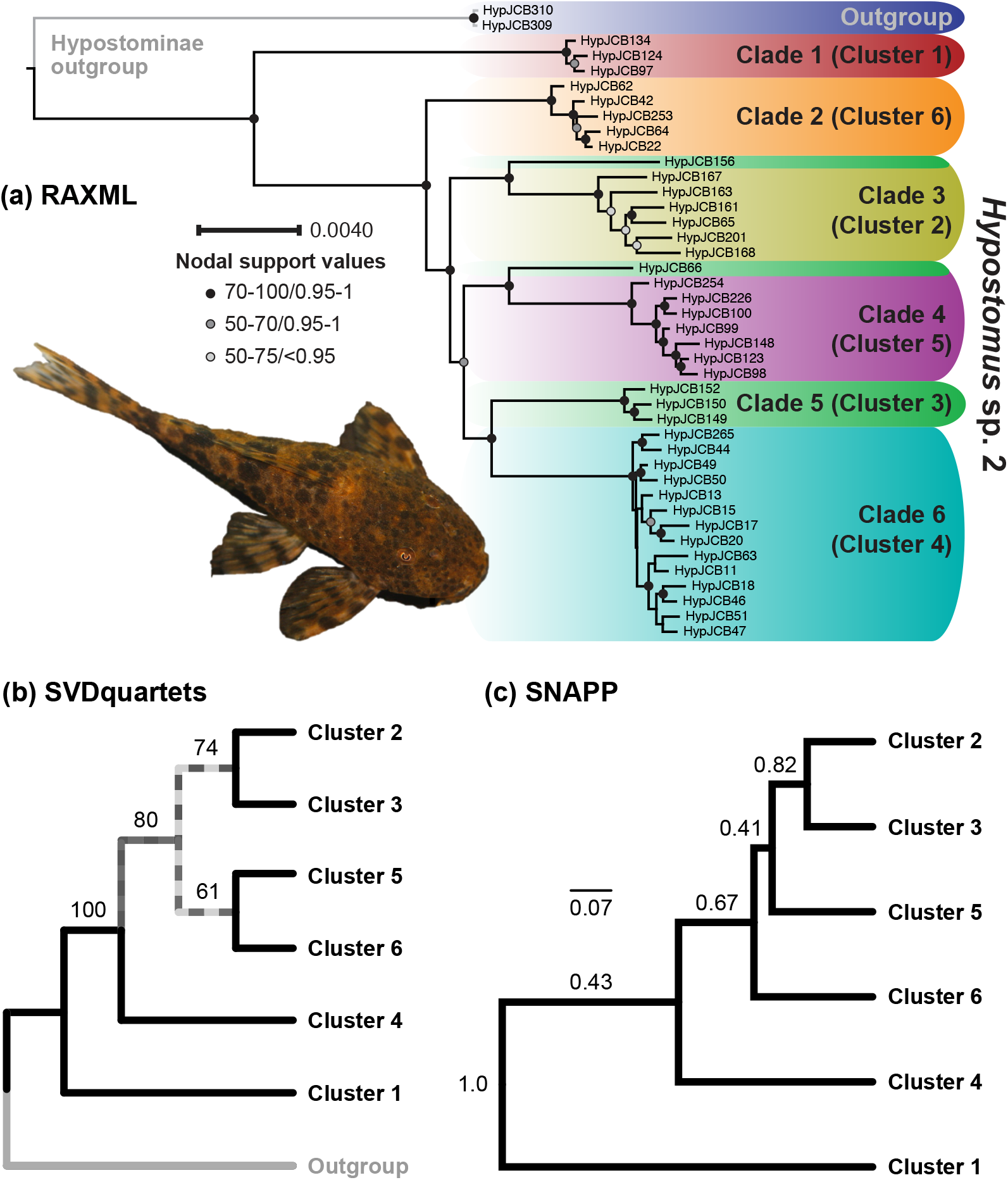
Phylogenomic gene tree and species tree relationships for *Hypostomus* sp. 2 lineages inferred from 227 ddRAD tags. (a) ‘Best’ maximum likelihood (ML) gene tree inferred in RAXML v8.2.8 (Stamatakis, 2014) while partitioning the dataset according to the optimum partitioning scheme and models of sequence evolution supported by PartitionFinder v1.1.1 (Table 1). Node circles are colored to reflect nodal support given by ML bootstrap proportions (≥ 50%) from RAxML/Bayesian posterior probabilities from MrBayes v3.2 (Ronquist et al., 2012), and clade colors correspond to genetic clusters in Figure 1c. A nodal support key is given next to an inset photograph of the study species. (b) Species tree inferred in SVDquartets (Chifman and Kubatko, 2014) under the multispecies coalescent. (c) Species tree from SNP-based analysis under the multispecies coalescent model in SNAPP (Bryant et al., 2012). Measures of nodal support shown in panels (a) and (b) include nonparametric bootstrap and Bayesian posterior probability values, respectively.

Species tree topologies from SVDquartets (Figure 2b) and SNAPP (Figure 2c) agreed with our gene tree results in placing cluster 1 sister to all other lineages with strong bootstrap and Bayesian posterior support, respectively. Otherwise, the species trees differed from the gene tree in several respects, including placing cluster 4 sister to a clade containing clusters 2, 3, 5 and 6, with strong support from SVDquartets bootstrap proportions but limited support in the SNAPP tree. Another key difference was that the SVDquartets species tree resolved cluster 5, which was polyphyletic in the gene tree, as sister to cluster 6 with good nodal support after accounting for incomplete lineage sorting under the multispecies coalescent model. A DensiTree plot revealed a variable and often high degree of uncertainty of relationships between clusters 2–6 across 5000 posterior trees from SNAPP, with many shallowly coalescing genes (Figure S3). Given higher nodal support from SVDquartets and the fact that SVDquartets has been shown to be consistent in the face of low levels of gene flow (Long and Kubatko, 2018), we took the SVDquartets result as our best estimate of the species tree. The multispecies coalescent gene tree from SVDquartets contained tip relationships similar to the ML and Bayesian gene trees but yielded slightly different placements of clusters 5 and 6 (Figure S4). The multilocus phylogenetic network (Figure S5) was treelike, with clades identical to our ML and Bayesian phylogenetic hypotheses, with boxes indicating uncertainty in relationships possibly due to incomplete lineage sorting, admixture, or other sources of homoplasy (see Appendix S1B for additional details).

Our *gsi* analyses yielded mixed results. On one hand, *gsi* findings rejected two predictions of the Paraná Capture Hypothesis, with *gsi*_T_ being comparably low to moderate and non-significant (*p* > 0.05) across drainage groups under tip configuration 1 (UP+UT *gsi*_T_ = 0.162; USF *gsi*_T_ = 0.291; Figure 3a) and configuration 2 (UP+USF *gsi*_T_ = 0.238; UT *gsi*_T_ = 0.231; Figure 3b). Overall, *gsi* values were randomly distributed, with values from different drainage groups overlapping, when plotted from calculations assuming the different configurations. On the other hand, the non-significant result under configuration 3 (UP *gsi*_T_ = 0.288; USF *gsi*_T_ = 0.291; UT *gsi*_T_ = 0.360; Figure 3c) was instead consistent with Paraná Capture Hypothesis predictions and a possible history of multiple captures involving Paraná source populations. While incomplete lineage sorting remained substantial, a greater degree of lineage sorting was observed within our two deepest-diverged empirical clades than the predefined drainage configurations above. Ensemble *gsi* was larger and significant when calculated for two of the main lineages in our species tree (Figure 2b), including cluster 1 (*gsi*_T_ = 0.662, *p* < 0.001; Figure 3d) and the clade formed by clusters 2 + 3 + 5 + 6 (*gsi*_T_ = 0.630, *p* < 0.001; Figure 3e).

**Fig. 3.**
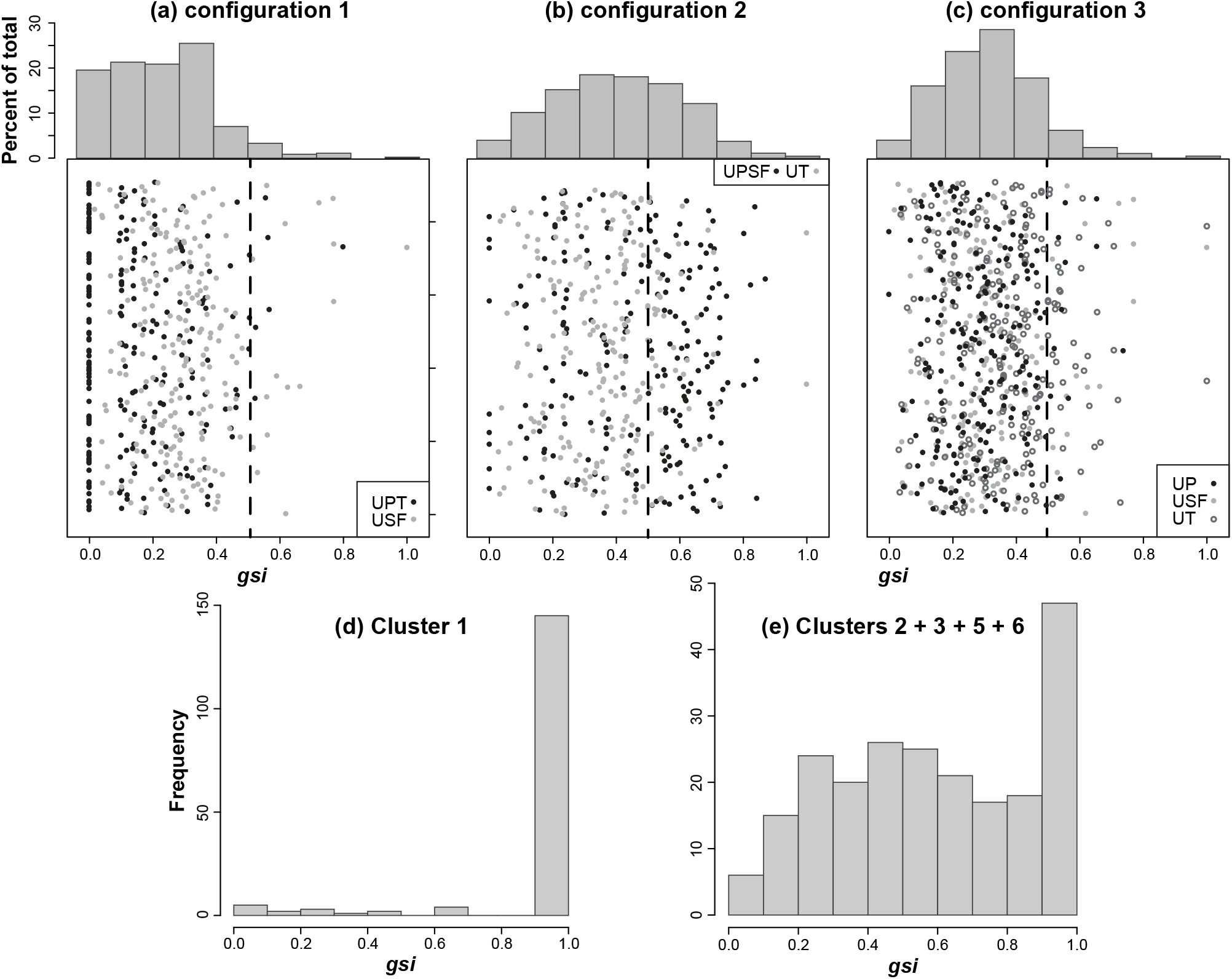
Genealogical sorting index (*gsi*; Cummings et al., 2008) histograms from three drainage configuration tests (a–c), and also calculated from two main ingroup clades (d, e) inferred in our best species tree shown in Figure 2b. Corresponding ensemble *gsi* (*gsi*_T_) calculations and *p*-values are given in the text. For panels (a) through (c), raw *gsi* values for each locus (circles) are plotted for each drainage grouping, with circles colored by drainage group, and vertical dashed lines indicate *gsi* = 0.5 (precisely half-way between unsorted and completely sorted allelic variation).

### 3.4. Demographic history and migration

Based on converged G-PhoCS runs (Appendix S1B), the deepest divergence in the species tree, separating cluster 1 from all other lineages, was dated to ∼1.25 million years ago (Ma) (highest posterior density, HPD = [0.274, 1.43 Ma]), whereas the remaining five lineages diversified since ∼220,000 years before present (ybp) in the mid–late Pleistocene (mean age range = ∼145,500 to 220,000 ybp) (Figure 4a; Tables S4 and S5). Estimated *N*_e_ values for clusters 1–6 indicated moderate to large contemporary population sizes ranging from ∼24,000 to ∼118,000 breeding individuals, but were largest in cluster 4 and much smaller in clusters 2 and 5, with non-overlapping HPDs. Plotting the G-PhoCS *N*_e_ estimates across the species tree revealed a pattern of population fragmentation following initial divergence of a very large ancestral population, followed by establishment of smaller populations corresponding to clusters 2–6 during the mid–late Pleistocene (Figure 4a). Likewise, SNAPP results indicated establishment of smaller populations, particularly in cluster 2 in the Tocantins basin (Figure S6; Table S6). These findings are consistent with Paraná Capture Hypothesis prediction 5 (Table 1). By contrast, SNAPP estimated *N*_e_ for cluster 3 to be much larger than that for other taxa, suggesting a massive ancestral population may have persisted in the Upper Paraná basin (area of majority membership for cluster 3; Tables S1) over mid-Pleistocene to recent.

**Fig. 4.**
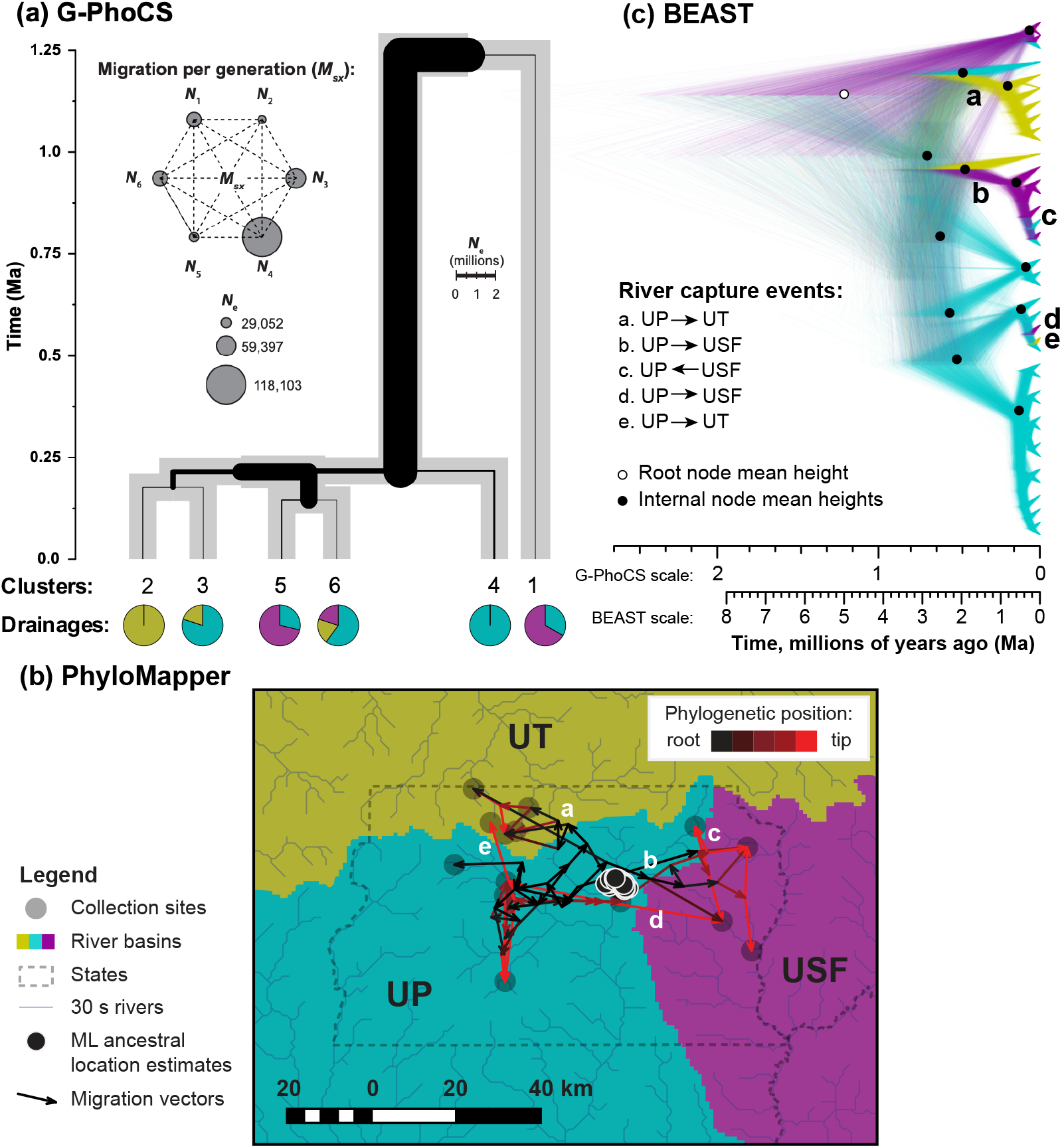
Results of demographic modeling analyses and phylogeographic analyses of *Hypostomus* genetic variation. (a) Results from Bayesian demographic analyses estimating parameters over the species tree in G-PhoCS v1.2.3 (Gronau et al., 2011). Tree node heights correspond to *T* in millions of years ago (Ma), while inner (black) branch widths are proportional to *N*_e_ estimates (inset scale). Drainage proportions for each cluster are represented by pie charts. In left inset, per-generation migration is visualized between clades, which are represented by gray circles with diameters proportional to their *N*_e_ estimates; migration proportions were weak and reflected scaled or total migration rates that were effectively zero (Tables 4, S4, and S5; Appendix S1B), hence they are shown as dashed lines. (b) Results from a spatial random walk model of the ancestral origin and dispersal routes of the ingroup lineage in PhyloMapper v1b1 (Lemmon and Lemmon, 2008). Arrows in (b) are migration vectors inferred from optimizing the model over an ultrametricized gene tree topology, with darker arrows showing inferred phylogeographic dynamics near the root of the tree and lighter (red) arrows indicating dispersal near the tips of the tree. (c) Result of discrete Bayesian spatial diffusion analysis in BEAST mapping sampling locations to drainage basins. A DensiTree plot of 5000 post-burn-in gene trees from the posterior of the analysis is shown with lineages colored to match their drainage areas. In panels (b) and (c), the same five non-random river capture (dispersal/vicariance) events were inferred, and these are labelled ‘a’ through ‘e’ along the internal or tip branches corresponding to the event.

**Table 4.**
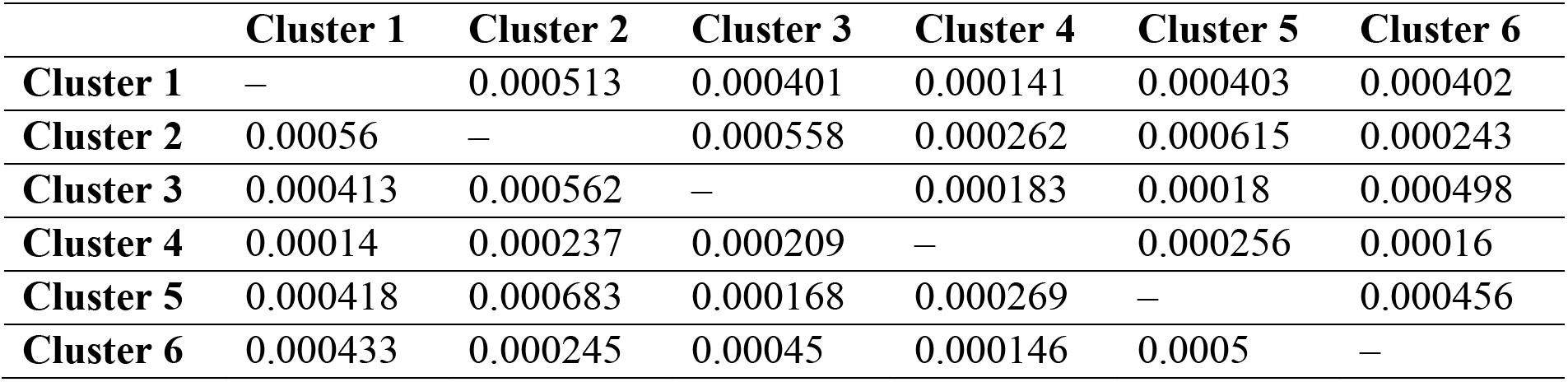
Pairwise migration rates per generation (*m_sx_*) between clades in the species tree, as estimated by G-PhoCS. Results are presented as means from each of 15 runs with the highest post-burn-in effective sample sizes (ESS) for *M_sx_*. See Table S4 and Table S5 in the Supporting Information for raw and converted migration parameter estimates and their credible intervals, respectively. After factoring in the timespan of each migration band (*τ_sx…_*; Gronau et al., 2011), these values are equivalent to effectively zero total migration rates (JCB, unpublished results).

|  | **Cluster 1** | **Cluster 2** | **Cluster 3** | **Cluster 4** | **Cluster 5** | **Cluster 6** |
| --- | --- | --- | --- | --- | --- | --- |
| **Cluster 1** | – | 0.000513 | 0.000401 | 0.000141 | 0.000403 | 0.000402 |
| **Cluster 2** | 0.00056 | – | 0.000558 | 0.000262 | 0.000615 | 0.000243 |
| **Cluster 3** | 0.000413 | 0.000562 | – | 0.000183 | 0.00018 | 0.000498 |
| **Cluster 4** | 0.00014 | 0.000237 | 0.000209 | – | 0.000256 | 0.00016 |
| **Cluster 5** | 0.000418 | 0.000683 | 0.000168 | 0.000269 | – | 0.000456 |
| **Cluster 6** | 0.000433 | 0.000245 | 0.00045 | 0.000146 | 0.0005 | – |

Results are presented as means from each of 15 runs with the highest post-burn-in effective sample sizes (ESS) for $M_{sx}$ . See Table S4 and Table S5 in the Supporting Information for raw and converted migration parameter estimates and their credible intervals, respectively. After factoring in the timespan of each migration band ( $\tau_{sx...}$ ; Gronau et al., 2011), these values are equivalent to effectively zero total migration rates (JCB, unpublished results).

Migration rate estimates from G-PhoCS strongly rejected prediction 5 of the Frequent Interdrainage Dispersal Hypothesis of gene flow between drainage basins or lineages (Table 1). Instead of detecting significant gene flow, we found that migration signals in our data were weak and yielded converted rates, *M_sx_*, that were effectively zero in all pairwise comparisons (≤ 1.23 × 10^−7^; Tables S4 and S5), which is indicated by dotted lines in Figure 4a (inset diagram).

### 3.5. Ancestor geographic location analyses

Several of our PhyloMapper results were consistent with Paraná Capture Hypothesis predictions (Table 1), with dispersal of *Hypostomus* sp. 2 from the Upper Paraná River into the Upper Tocantins basin. First, we rejected the null hypothesis of no phylogeographic association of genotypes (mean dispersal distance, *ψ* = 20.35; *p* < 0.001). Second, center test results failed to reject the null hypothesis (ln *L* = −225.430) that the ancestral geographic location was located at the center of the sampling localities (−15.714223 °S −47.757160 °W) in the Upper Paraná basin, as compared to the alternative ML-estimated ancestral location (ln *L* = −224.717; −15.694609 °S −47.665523 °W; likelihood ratio test statistic, χ^2^ _(2, 2)_ = 1.426; *p* = 0.49), which was consistently obtained across 100 different optimizations (Figure 4b). Results from optimizing a spatial random walk model over the gene tree inferred that migration vectors originate in the Upper Paraná (near the gene tree root) and indicated progressive and overlapping pathways of dispersal into adjacent basins, with recent colonization of the Tocantins basin, and no migration routes from the Tocantins into other basins (Figure 4b). Third, we rejected the null hypothesis of a random direction of migration (overall directionality test; mean displacement = 0.0597; *p* < 0.001). Contrasting the above, our a priori direction test results did not support significant directional dispersal from the Upper Paraná into the upper Tocantins (‘north’ mean angle difference = 1.61 radians; *p* > 0.05; ‘northwest’ mean angle difference = 1.54 radians, *p* > 0.05) or São Francisco basins (‘east’ mean angle difference = 1.56 radians, *p* < 0.001).

Our converged BEAST discrete trait diffusion analyses (Appendix S1B) yielded a single best tree with relationships similar to that of our RAxML gene tree (Figure 2a), but with clade 2 (cluster 6) sister to clade 6 (cluster 4) rather than sister to clades 3–6 (clusters 2–5; Figure S7). The inferred patterns of movements between drainage basins, presumably due to river capture events, were essentially identical to those inferred from our PhyloMapper analyses. Results were sufficiently similar that we could identify five river capture events shared between PhyloMapper results (Figure 4b [a–e]) and a DensiTree representation of 5000 post-burn-in trees from the BEAST discrete phylogeography analysis (Figure 4c [a–e]). Overall, these results highlighted two river captures from the Upper Paraná into the Upper São Francisco basin, one river capture in the opposite direction, and two river captures from the Upper Paraná into the Upper Tocantins basin (Figure 4b,c); thus, they strongly supported the Paraná Capture Hypothesis.

### 3.6. Effects of varying RAD-seq assembly parameters on downstream genetic analyses

*Qualitatively* comparing assembly output and gene tree and admixture group inferences derived from 280 PyRAD assemblies run using a range of parameter values, including our empirical assembly, yielded wide variation in practically important outcomes including the total number of ddRAD-seq loci generated, which ranged from 16 to 456 loci (i.e. tags), and total number of SNPs, which ranged from 255 to 8432 loci (Figure 5a–e). At 227 ddRAD tag sequences and 3829 SNP loci, our empirical datasest fell within the intermediate values of these ranges, suggesting our results were broadly representative. The pO parameter had the greatest effect on assembly outcomes, and increasing the Phred offset was negatively correlated with numbers of loci and SNPs (Figures 5c,f), which tended to be lowest across assemblies when pO was set to more stringent values of 26 or 33, and highest when pO was set to the PyRAD default setting of 20 (used in our empirical assembly) or a lower value of 14. Also concerning *qualitative* sensitivity analyses, our concatenated ML gene tree (Figure 2a) clustered with other trees or in the middle of ordinal treespace, irrespective of whether the PCoAs were based on *d*_G_ (Figure 6a) or *d*_RF_ (Figure 6b) distances, which reflect different kinds of topological variation. The mean *d*_RF_ for the best tree (relative to the best trees from all 279 other assemblies) also fell well within the distribution of the Robinson–Foulds distances, close to the mean value (Figure 6c), indicating that the result was typical. Our admixture group results from fastSTRUCTURE were even more streamlined: the best *K* of 6 from our empirical analyses (Figure 1b) overlapped with the levelling off of fastSTRUCTURE marginal likelihood values over *K* in results from most assemblies, and a histogram of the best component *K* values across assemblies showed *K* of 6 or 7 were common outcomes (mean *K* = 6.45), with *K =* 6 being the mode of the distribution.

**Fig. 5.**
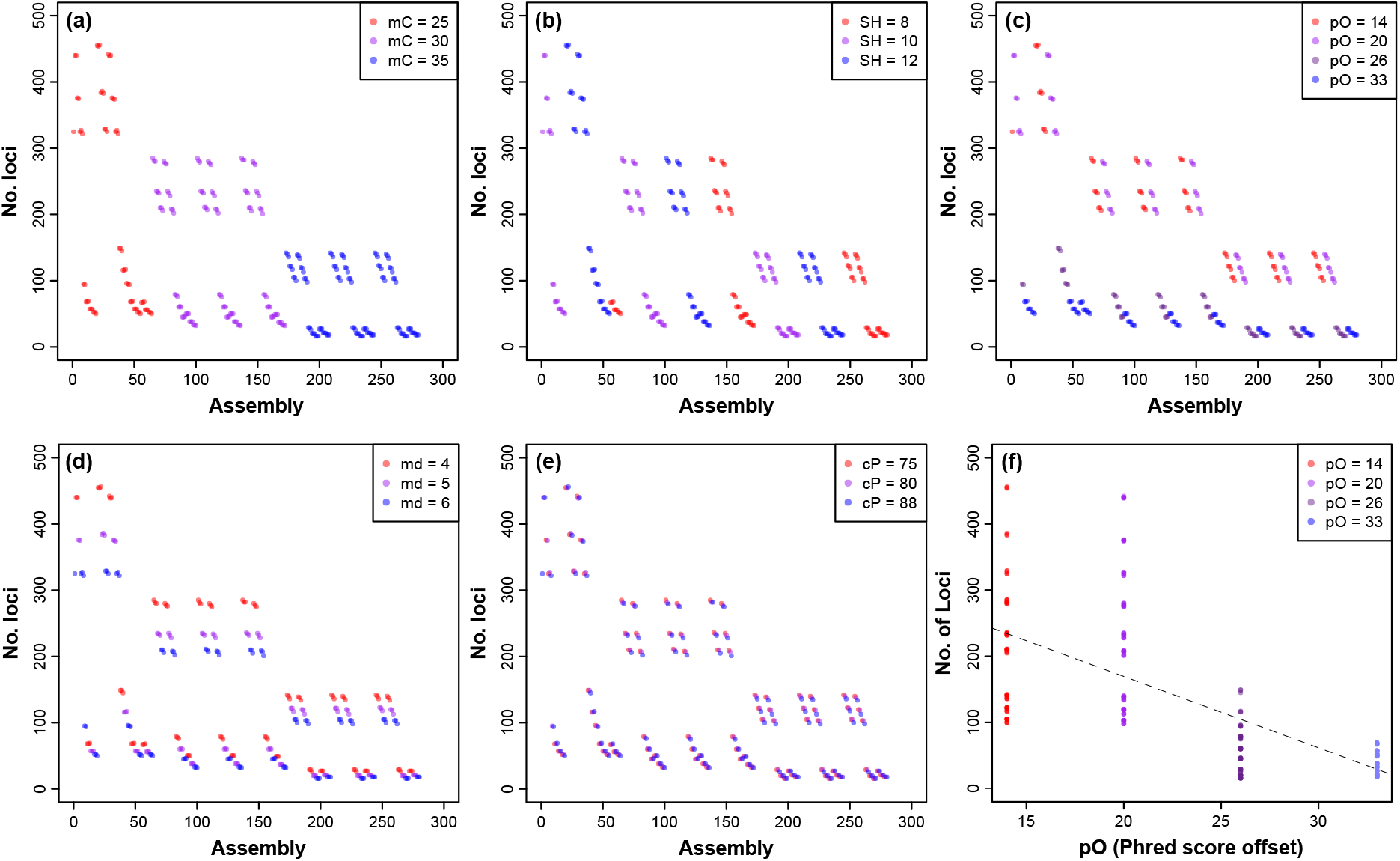
Assembly output under varying ddRAD-seq assembly parameters in PyRAD (Eaton, 2014). The first five panels show correlations between the number of ddRAD tag sequences output by PyRAD and settings for five test parameters that we varied: (a) mC, minimum number of samples in a final locus; (b) SH, max. individuals with a shared heterozygous site; (c) pO, Phred score offset; (d) mD, minimum depth of coverage; and (e) cP, clustering percentage threshold. Each of 280 PyRAD assemblies of our *Hypostomus* dataset is represented by a single result (circle) plotted against its index, ordered by increasing mC, SH, and pO, etc. (f) Biplot showing significant negative relationship between the number of output ddRAD tag sequences and pO (*p* < 0.05 in generalized linear modeling or nonlinear ordinal regression; Appendix S1B).

**Fig. 6.**
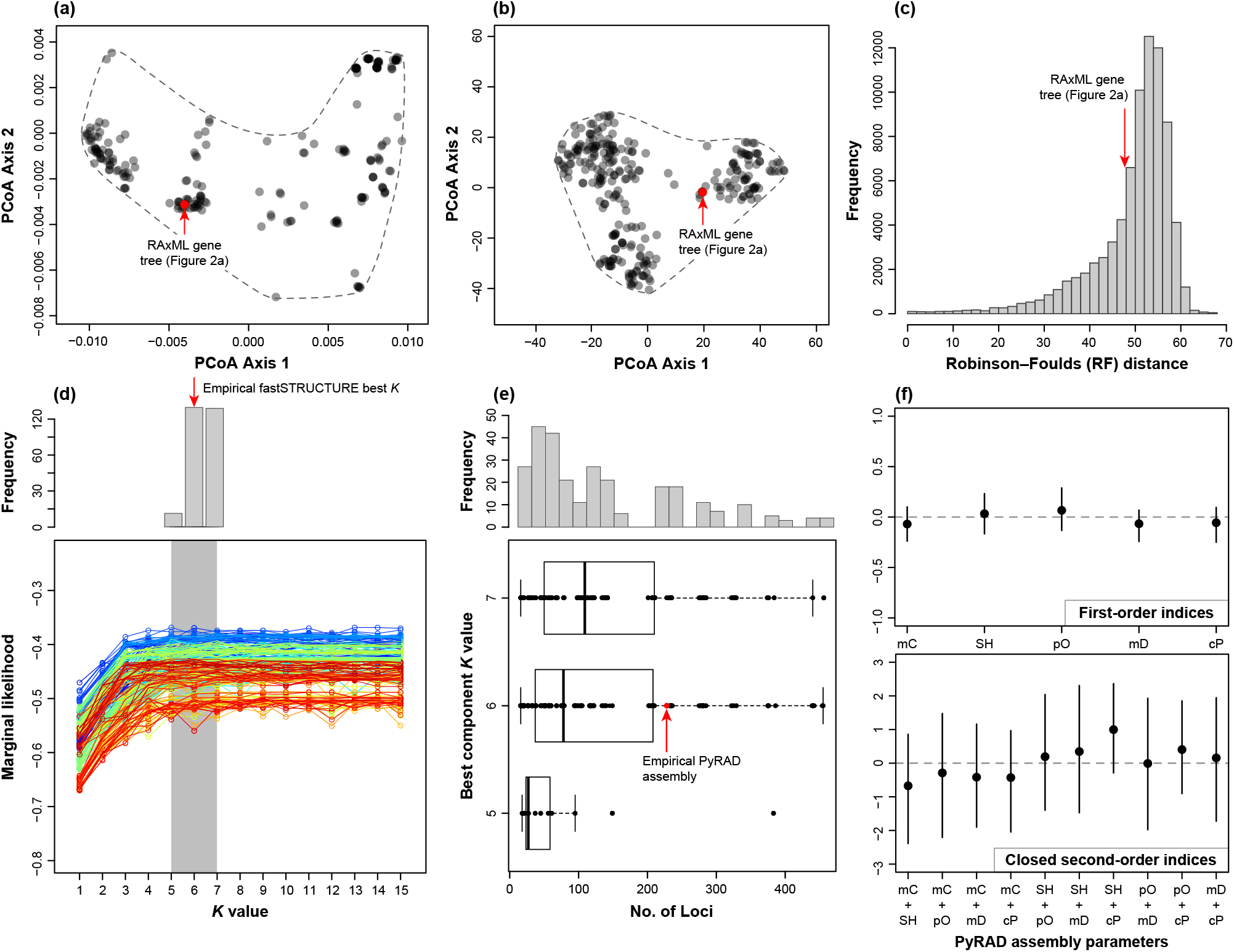
Sensitivity of our downstream gene tree and population admixture group analyses, and number of ddRAD tag sequences assembled, to varying assembly parameters in PyRAD (Eaton, 2014). Results are shown for genetic analyses of a test set of 280 assemblies of our *Hypostomus* ddRAD-seq dataset generated by varying parameter settings in PyRAD, including the empirical assembly. Panels (a)–(e) show results of *qualitative* sensitivity analyses, while panel (f) shows results of *quantitative* Sobol’ variance-based sensitivity analyses. Positions of the RAxML gene tree from our main empirical analysis (Figure 2a) are shown (red circles, arrows) within the treespace formed by PCoA plots of *d*_G_ (a; geodesic distances) and *d*_RF_ (b; Robinson-Foulds [RF] distances) between the best RAxML gene tree from each assembly in the test set. The histogram in panel (c) shows the position of our main gene tree (mean *d*_RF_ = 47.65) within the distribution of pairwise *d*_RF_ values between all 280 gene trees. (d) Distribution of best component *K* values (analogous to interpretation of empirical results in Section 2.4 above) for each assembly (mean *K* = 6.45) over (e) a plot of marginal likelihoods for the corresponding fastSTRUCTURE models over *K* = 1–15 computed for each assembly. (f) Effects of varying assembly parameters on number of output ddRAD tag loci, and (g) positive correlation between number of loci and best component *K*. (h, top) Plots of Sobol’ first-order indices, or main effect contributions, of each of five parameters varied in PyRAD on assembly output variance. (f, bottom) Plots of Sobol’ second-order indices, or interaction effects, of pairs of PyRAD parameters on assembly output variance.

*Quantitatively* assessing the contributions to total variance at four assembly output variables using variance-based sensitivity analysis (Sobol’, 1990; Saltelli et al., 2008) revealed that fractional contributions of the individual variables (*X_i_*) alone, or in terms of their interactions, were minimal. None of the five varied parameters made a significant contribution to the total output variance individually, as judged by the overlap of first-order indices (*S_Xi_*) with zero (Figure 6f, top), or had joint effects on the output that were different from their individual effects, as judged by the overlap of second-order indices (*S_Xi_*_,*Xj*_) with zero (Figure 6f, bottom). Despite this, pO and mD were closest to having significant individual main effects, and the interaction effect of SH+cP parameters was nearly significant.

## 4. Discussion

The Brazilian Shield highlands present an excellent system for studying the effects of river capture events on the phylogenetic structure and historical biogeography of stream fish assemblages (Ribeiro, 2006; Lima and Ribeiro, 2011; Aquino and Colli, 2017). The main goal of this study was to use ddRAD-seq phylogeography to test for predicted spatial-genetic effects of river capture in the Paraná–Tocantins–São Francisco headwater river system in central Brazil (Table 1; Figure 1), derived from two main hypotheses advanced in a previous phylogenetic community ecology study of freshwater fishes in this system (Aquino and Colli, 2017). We focused on a distinct lineage or species complex (Bagley et al., 2019) of suckermouth armored catfish, *Hypostomus* sp. 2 (Aquino et al., 2009), that is endemic to and distributed across this region, and was included in Aquino and Colli’s (2017) study. Using phylogeographic analyses of genome-wide data, we tested hypotheses on the direction and frequency of river-capture events informed by geological and ecological data. Overall, while distinguishing between hypotheses proved difficult, and evidence was partially mixed, the accumulated data and results largely support predictions of the Paraná Capture Hypothesis, while providing less support for the non-mutually exclusive Frequent Interdrainage Dispersal Hypothesis (Aquino and Colli, 2017). Below, we explore varying aspects of support for our a priori hypotheses and insights into the phylogeographic history of *Hypostomus* sp. 2. We also discuss the utility of our phylogeographic approach based on ddRAD-seq data to distinguish river capture from recent wet-connection mechanisms that could have facilitated interdrainage dispersal. Last, we explore implications of our genetic sensitivity analyses evaluating the robustness of our assembly outcomes and downstream genetic results to varying ddRAD-seq assembly parameters in PyRAD, framing the discussion in the context of two areas where greater research attention is needed.

### 4.1. Testing hypothesized effects of river capture in the central Brazilian Shield

The majority of our results including 9/11 (82%) analyses and statistical tests (Figure S8) agreed with predictions of the Paraná Capture Hypothesis (Table 1). Consistent with prediction 1 of this hypothesis, *Hypostomus* sp. 2 populations exhibited a significant phylogeographic association (section 3.5), and Upper Tocantins or Upper São Francisco basin alleles were phylogenetically nested within or sister to alleles sampled from the Upper Paraná basin four to five times in each reconstructed gene tree (Figures 2a, 4, S2, S4 and S7). This interpretation is further supported by a higher diversity of genetic clusters/lineages of *Hypostomus* sp. 2 in Upper Paraná River basin headwaters, as compared with the headwaters and streams of neighboring basins (four versus three each, respectively; Figure 1; Table S1). Also, in our species trees, cluster 2 from the Tocantins basin was consistently supported as sister to cluster 3 (Figure 2b,c), which was primarily from three Upper Paraná River sites (Figure 1a). This relationship agrees with vicariant isolation due to river capture between populations across the Upper Paraná–Upper Tocantins drainage divide and was strongly supported after accounting for incomplete lineage sorting in SVDquartets (Chifman and Kubatko, 2014; Long and Kubatko, 2018).

Our efforts at phylogeographic inference and hypotheses testing were complicated by two patterns—lack of lineage monophyly across loci and lineages distributed in multiple basins, the latter of which led to gene tree–species tree incongruence. In particular, while primarily from the Upper Paraná basin, cluster 3 genotypes were also sampled from one Upper Tocantins site. The corresponding multilocus genotype was from a single individual, HypJCB66, that had derived, polyphyletic positions in our gene tree and network results, and individual HypJCB156 from the Paraná basin exhibited a similar polyphyletic pattern (Figures 2a, S5). While these individuals could represent additional dispersals to or from the Upper Paraná basin, three of our findings suggest that their phylogenetic positions, and the gene tree–species tree incongruences they represent, are more likely an artefact of incomplete sorting of allelic polymorphism through large ancestral populations over a geologically recent timeframe of divergence (e.g. reviewed by Avise, 2000; Funk and Omland, 2003). These include the (1) substantial population sizes, (2) effectively zero inter-lineage migration rates, and (3) Pleistocene divergence times inferred for cluster 3 and others (Tables 4, S4–S6). Assuming neutral loci sampled from a Wright–Fisher population with no migration, coalescent theory predicts that 9*N*_e_ to 12*N*_e_ generations are required for two lineages diverging in allopatry to become monophyletic at 95% of nuclear loci (Hudson and Coyne, 2002). Based on our G-PhoCS results, the time to nearly complete lineage sorting in *Hypostomus* sp. 2 for our nuclear SNP data would therefore be since *c.* ∼2.87–3.82 Ma, vastly predating our ingroup root divergence time of ∼1.25 Ma (∼3.9*N*_e_ generations). Under similar calculations for the cluster 2–cluster 3 split, we estimate that the time to nearly complete lineage sorting would be since *c.* ∼0.75–1.0 Ma, again vastly predating our estimated divergence time of ∼0.15 Ma (145,517 ybp) for these two lineages. Taken together, all of the above relationships strongly suggest that our results have been affected by incomplete lineage sorting. From the initial ingroup divergence to the cluster 2–3 split, we thus expect lineages within *Hypostomus* sp. 2 to retain part, or perhaps much, of their ancestral allelic variation, even if they are distinct lineages, incipient species, or reproductively isolated species. Also, we interpret their alleles as being caught in the transition from polyphyly to monophyly in contemporary populations (Avise 2000; additional details in Appendix S1C). This view matches well with our *gsi* results (e.g. Figure 3) and related interpretations, and it also permits a clear reconciliation of results conflicting with, versus supporting, our predictions for the Paraná Capture Hypothesis.

Traditionally, species-level inferences of ancestral areas in historical biogeography were made using simple rule-based algorithms to determine sequences of changes in species geographical distributions over phylogenies, particularly using endemic taxa (e.g. Bremer, 1992; Ronquist, 1997; Hausdorf, 1998). By contrast, phylogeographic models in PhyloMapper and BEAST permit inferring statistically robust estimates of the ancestral areas and spatial dynamics of population lineages based on probabilistic models of dispersal across a continuous or discrete landscape, over ultrametric or time-calibrated phylogenies (e.g. Lemmon and Lemmon, 2008; Lemey et al., 2009). While phylogenetic results discussed above provide indirect evidence for an Upper Paraná ancestral area for the ingroup, our ML and Bayesian phylogeographic modeling results provide direct evidence and finer resolution into the ancestral area supported by the data. Consistent with Paraná Capture Hypothesis prediction 3 (Table 1), results from PhyloMapper’s spatial random walk model and center test supported the geographical location of the ingroup ancestor as occurring near the center of the area in headwater reaches between sites 7 (Ribeirão Pipiripau) and 10 (Córrego Saco dos Pilões) in the Rio São Bartolomeu catchment of the Upper Paraná basin (Figures 1 and 4b,c). Results of the overall directionality test also supported Paraná Capture prediction 6 (Table 1) of non-random, directional dispersal. Migration vectors also clearly indicated dispersal into both adjacent basins from the Upper Paraná ancestral area. As PhyloMapper results may not reflect sources of uncertainty associated with sequentially inferring the phylogeny and then the parameters of the spatial random walk model, we complemented this analysis with a BEAST discrete trait diffusion analysis in which the tree and diffusion model parameters were jointly estimated in a Bayesian framework. The BEAST analysis inferred a slightly different tree and deeper divergence times (Figure S7), probably because it used a gene tree framework, and gene divergences predate species divergences (e.g. Edwards and Beerli, 2000). Still, BEAST inferred the same five geographical patterns of dispersals into and out of the Paraná basin as those inferred in PhyloMapper (Figure 4a–e), suggesting results were robust to topological uncertainty and algorithmic differences between software programs. A weakness of these diffusion modeling results is that they are all based on gene trees, which we concluded above were heavily influenced by incomplete lineage sorting. Yet, forcing our gene trees to fit the species tree relationships would, at most, drop our inferences of shared river capture events across these two sets of analyses from five shared events down to two shared events.

Another prediction of the Paraná Capture Hypothesis that was supported by our results was prediction 4 that multilocus genotypes of fish populations would provide evidence of founder effects or population expansions into the Tocantins basin (Table 1). Demographic inferences from G-PhoCS and SNAPP results yielded *N*_e_ values that we interpret to be consistent with a relatively recent founder effect in Tocantins cluster 2 (Tables S5 and S6; Figures 4a and S6). Consistent with this same hypothesis, we find larger ancestral population sizes and consistently higher *N*_e_ values for the predominantly Upper Paraná cluster 3 across analyses; indeed, SNAPP population size estimates yielded a massive *N*_e_ estimate for cluster 3.

Two sets of Paraná Capture Hypothesis predictions were not consistently supported by our results, including prediction 5 that a priori direction tests in PhyloMapper would support significant migration into the Tocantins basin (Table 1). This result likely owes to the inferred patterns of migration being from the Paraná into both adjacent basins, rather than a single basin. Alternatively, the a priori direction tests may have been non-significant if they had low power to detect directional migration over small sampling areas or sample sizes. The other Paraná Capture Hypothesis prediction set that was not supported by our results was prediction 2, under which we expected that *gsi* would be significant for drainage configurations 1 and 2. While expected levels of allele sorting were not met, those for drainage configuration 3 under this hypothesis were met in the form of non-significant *gsi*. While we recognize that statistically rejecting a null hypothesis of random genealogical sorting would provide a firmer basis for inference than failing to reject the null, this outcome is nonetheless in line with our a priori expectations. Importantly, all of our *gsi* results also match well with our interpretation that incomplete allelic sorting in recently diverged lineages has played a major role influencing gene tree–species tree incongruence in *Hypostomus*, as discussed above.

Compared with support for most predictions of the Paraná Capture Hypothesis above, fewer of our results, including 2/6 (33%) analyses and statistical tests (Figure S8), supported predictions of the non-mutually exclusive Frequent Interdrainage Dispersal Hypothesis. Phylogenetic and coalescent-based demographic modeling results lent support to predictions 1 and 4 of this hypothesis, including the expectation of lineage paraphyly/polyphyly across areas and loci. Specifically, clustering and phylogenetic analyses inferred *Hypostomus* sp. 2 clusters 1 (clade 1), 6 (clade 2), 5 (clade 4), and 3 (clade 5) each to be distributed in multiple drainage basins (Figures 1a and 2; Table S1). Also, our a priori directionality tests in PhyloMapper were non-significant, consistent with what we would expect if *Hypostomus* sp. 2 lineages had experienced multidirectional dispersal and vicariance between basins, due to frequent and randomly-oriented river captures over the Pleistocene to recent. However, even these patterns of support remain problematic. As discussed above, para-/polyphyletic patterns of areas and loci in our gene trees have likely been heavily influenced by incomplete lineage sorting; hence, these patterns cannot be taken objectively as positive evidence in favor of the Frequent Interdrainage Dispersal Hypothesis. Moreover, the majority of analyses and tests provided evidence against predictions of the Frequent Interdrainage Dispersal Hypothesis. The most incriminatory among these included results supporting effectively zero pairwise estimates of migration per generation between clades in G-PhoCS (Figure 4a; Tables 4, S4–S6), which rejected prediction 5 under this hypothesis, and the significant PhyloMapper overall directionality test (section 3.5), which rejected Frequent Interdrainage Dispersal Hypothesis prediction 4.

### 4.2. Pleistocene phylogeography of Hypostomus catfish

Our sampling of *Hypostomus* sp. 2 spans the currently known distribution of this lineage or species complex, whose alpha taxonomy we are formally assessing elsewhere, with colleague Y. F. F. Soares. Given its limited geographical range, this species expectedly shows relatively limited or shallow evolutionary diversification, with the deepest molecular divergence between its lineages occurring between clade 1 from the upper Paraná and São Francisco basins and all other clades (Figures 1a and 2). This divergence was dated to 1.25 Ma in the early Pleistocene by G-PhoCS (Figure 4a), with Bayesian credible intervals within the epoch (maximum 1.43 Ma; Table S5), and up to around ∼2.5 Ma were inferred by SNAPP (Figure 2c; Table S6). The timescale of diversification within *Hypostomus* sp. 2 thus post-dates the formation of modern South American drainage basins since ∼10 Ma in the Miocene, based on reconstructed paleogeography and paleodrainage routes (e.g. Lundberg et al., 1998; Hoorn et al., 2010; Albert and Reis, 2011), as well as the origin of the Hypostominae estimated by Silva et al. (2016) to be at around the same time, at ∼12 Ma, based on Bayesian divergence time analyses. Historical biogeographical reconstructions by Silva et al. (2016) inferred six independent colonizations of the Paraná basin by different *Hypostomus* lineages since the Miocene; therefore, our study highlights post-colonization patterns of diversification and river capture at finer spatial and temporal scales following the establishment of an ancestral *Hypostomus* population in the region. The estimated initial timing of ingroup population divergence matches a period of global cooling and expansion–contraction of savannas, with cooler-than-modern temperatures during the growth of Northern Hemisphere ice sheets (e.g. Zachos et al., 2001). By contrast, clades 2 through 6 are inferred to have diverged since *c.* ∼220,000 ybp in the mid–late Pleistocene, based on divergence dates from G-PhoCS with a relatively high degree of precision (narrow credible intervals; Figure 4a; Tables S4 and S5). This mid-late Pleistocene cluster of divergences corresponds to a period of exacerbated Pleistocene climatic fluctuations during intensified glacial cycles, with credible intervals spanning *c.* ∼265,500–26,700 ybp (Table S5) during Marine Isotope Stages (MIS) 3–8 (Lisiecki and Raymo, 2005). The peak-likelihood *t*_MRCA_ estimate for clusters 2–6 dates closest to MIS 7, one of the last three full interglacial periods (McManus et al., 1999), which could have favored larger water volumes resulting in greater erosion. However, the geomorphological setting appears not to have restrained river capture to MIS 7. In particular, the single arguably most important of these divergences for our purposes—the ∼145,500 ybp Paraná–Tocantins divergence between clusters 2 and 3 in our species tree (Figure 4b), falls in the MIS 6 glacial just following the interglacial of MIS 7. Therefore, we hypothesize that geological factors combined with warmer, wetter climates during MIS 7 created an erosional setting favoring headwater river capture of parts of the Upper Paraná basin into the Upper Tocantins basin via headward stream erosion, promoting the last divergence between clusters 2 and 3.

### 4.3. Distinguishing river capture and wet-connection mechanisms of interdrainage dispersal

We have implicitly assumed that dispersal and isolation of stream fish lineages in the Paraná– Tocantins–São Francisco drainage system has been caused by capture(s) of headwater tributaries by adjacent basins, mediated by geomorphological changes, as indicated by independent geological and ecological data (Saadi, 1993; Aquino and Colli, 2017). Here, river capture is the null hypothesis based on external information, and ‘wet connections’ mechanisms of interdrainage dispersal, such as those demonstrated for some New Zealand freshwater fishes (e.g. Craw et al., 2007; Burridge et al., 2006, 2008), represent alternative hypotheses for spatial-genetic patterns we have elucidated for *Hypostomus* sp. 2. However, we see no cause for concern over the utility of our approach to distinguish between river capture versus wet-connection mechanisms of dispersal and vicariance. This is because the only geographic evidence for wet connections between the study drainages supports a recent, ephemeral connection between the upper Paraná and Tocantins (e.g. Maranhão River) rivers in an approximately ∼25 km^2^ area of Vereda Grande spring in the Águas Emendadas Ecological Station. The timing of the origin of Vereda Grande is constrained to less than 26,000 ybp by stratigraphic and palynological records (Barberi, 2008, references therein), thus bidirectional water flows during periods of inundation of the wetland only could have facilitated interdrainage dispersal in this small area during the late Pleistocene (Ribeiro et al., 2008) to our knowledge. We did not sample *Hypostomus* from Vereda Grande; however, mean divergence times of all *Hypostomus* sp. 2 clades in this study, and their 95% credible intervals, substantially predate this period. Therefore, we reject the hypothesis that late–end Pleistocene wet connections, instead of river captures, have shaped the resulting genetic patterns. In light of these findings, and the precise estimates of *N*_e_ and *T* parameters obtained herein, we conclude that our NGS-based phylogeographic approach is suitable for distinguishing between river capture and wet-connection mechanisms of interdrainage dispersal, especially when inferred dates of river capture (1) substantially predate evidence for wet connections and (2) coincide with interglacial periods when headward erosion should be more commonplace.

### 4.4. Effects of varying RAD-seq assembly parameters on downstream genetic analyses

The ddRAD-seq (Peterson et al., 2012) approach employed herein and related ‘RADseq’ approaches have revolutionized the fields of population genomics and phylogeography due to their numerous advantages, including low cost, high throughput, and applicability to non-model organisms (e.g. reviewed by Andrews et al., 2016). While the ability to assemble the data de novo is also an advantage, the frequent lack of genomic resources (e.g. reference genomes) for focal taxa makes it difficult to validate the orthology of the resulting SNP loci. Additionally, a variety of approaches are available for assembling sequence data from RAD-seq experiments and calling SNPs, including Stacks (Catchen et al., 2011), PyRAD (Eaton, 2014), and ipyrad (Eaton and Overcast, 2020), among others, and these approaches differ in their bioinformatic details. As a result, there are no widely agreed upon best practices for which software or settings to use when assembling ddRAD-seq data, and the onus is on researchers to (1) test the robustness of downstream genetic results to varying the assembly parameters, and (2) attempt to optimize assembly parameter settings and other bioinformatics steps (reviewed by Rodríguez-Ezpeleta et al., 2017; O’Leary et al., 2018; McCartney-Melstad et al., 2019). We consider these to be important areas for improvement, as, in practice, biologists still commonly select an assembly, or SNP calls, from a single assembler/pipeline run with a fixed set of parameter values (e.g. clustering percentage), or a small number of runs. Results that are reported in a study may thus stem entirely from a single assembly and remain unvalidated. This is problematic because assembly parameters can have substantial effects on the resulting assembly and can also substantially affect downstream genetic analyses, including inference of population structure, demography, or phylogenetic relationships (e.g. Linck and Battey, 2019).

Regarding the two areas of improvement mentioned above, we rigorously tested the sensitivity of our assemblies and our downstream genetic results to varying assembly parameters in PyRAD using a comprehensive set of qualitative and quantitative analyses. A common challenge in RAD-seq studies is determining a reasonable setting for merging putatively homologous reads to identify orthologous loci, which in PyRAD is controlled by the clustering percentage threshold parameter (cP, herein). We found our assembly and genetic analysis results to be relatively insensitive to varying this parameter across a range of values from 75% to 88%, for example with virtually no effect on output ddRAD-seq loci (Figure 5e) and non-significant individual or interaction effects of this parameter on several genetic analysis outcomes in variance-based sensitivity analyses (Figure 6f). Our results also showed that our empirical assembly was intermediate in values of output ddRAD-seq loci and SNPs, and that our genetic results did not constitute outliers, relative to the other 279 assemblies in our test set (Figures 5 and 6). These findings indicate that our empirical results were both broadly representative and unlikely to be heavily biased by our choice of assembly parameters during bioinformatic processing of the data (additional details in Appendix S1). Moreover, that very few of our loci were flagged as putative paralogs (section 3.1; Appendix S1) suggests that assembly settings that we used were effective in guarding against paralogy to instead obtain orthologous loci.

Recent studies have found that varying cP in PyRAD, and related parameters in Stacks (Catchen et al., 2011), over a wider range can be informative for optimizing this parameter setting and guarding against incorrect handling of paralogous loci during assembly (McCartney-Melstad et al., 2019). However, in view of the limited evidence for paralogy in our empirical assembly, further work to optimize this parameter setting for our dataset would therefore seem unnecessary. Indeed, we view paralogy as more of a concern for assemblies in our test set based on less stringent settings (e.g. lower mC, pO, and mD values and higher SH values) than our empirical assembly. These less stringent assemblies are more likely to have resulted in undersplitting or oversplitting of paralogous gene regions, or yielded less accurate assemblies. While it appears to have worked well for our purposes, the clustering threshold level of 88% similarity used in our empirical assembly may not be appropriate for other RADseq or ddRAD-seq datasets, even in freshwater fish taxa. Given that the levels of genetic variation and paralogy are inherently species-specific, different clustering levels may be needed to effectively handle paralogs and identify and assemble orthologous loci in other systems (Rodríguez-Ezpeleta et al., 2017; O’Leary et al., 2018; McCartney-Melstad et al., 2019). We recommend that other authors employ sensitivity analyses similar to those used herein, or pipelines aimed at optimizing assembly parameters (McCartney-Melstad et al., 2019), in order to gauge the levels of sensitivity of downstream genetic results to varying RAD-seq assembly parameters and select optimal assemblies and results when analyzing ddRAD-seq datasets.

### 4.5. Merging phylogeography and phylogenetic community ecology

Recent studies have called for or empirically demonstrated how integrating phylogeography with other disciplines in ecology and evolution (e.g. community ecology, dispersal ecology, ecological niche modeling) can aid distinguishing the roles of historical and ecological processes in shaping diversification, community assembly and species turnover, and dispersal (Craft et al., 2010; Dexter et al., 2012; D’Amen et al., 2013; Marske et al., 2013; Hoban et al., 2019). In the context of this new trend towards a unified ecological and evolutionary framework, our study presents a first step towards integrating phylogeography and community ecology to understand the composition, evolution, and genetic diversity of stream fish communities in the central Brazilian Shield. We resampled the same headwater sites investigated in a community phylogenetic study (Aquino and Colli, 2017), to test hypotheses about the effects of river capture events suggested by the community data using Next-Generation phylogeography. By supporting their hypotheses, our results suggest that inferences of phylogenetic community ecology reflect historical processes, which phylogeography can further clarify. We recommend a comparative phylogeographical study of freshwater taxa in this study system to test the generality of patterns and processes of diversification that we inferred for *Hypostomus* sp. 2. However, given the confounding effects of incomplete lineage sorting revealed by our study, we predict that codistributed freshwater taxa may not exhibit phylogeographic congruence (*cf.* Avise 2000) with *Hypostomus* sp. 2. Genetic signatures of shared effects of drainage connectivity and geometry on demographic history (e.g. mediated by geological changes) may have been obscured by complex patterns of superimposed river capture events. Also, taxa with smaller ancestral population sizes or more ancient divergences may have experienced more complete lineage sorting (Hudson and Coyne, 2002; Cummings et al., 2008). Thus, it would be valuable to investigate the demographic histories of codistributed freshwater species, in a comparative context, using similar ddRAD-seq data and methods accounting for mutational and coalescent stochasticity.

## CRediT Authorship Contribution Statement

**Justin C. Bagley:** Conceptualization, Data curation, Formal analysis, Investigation, Methodology, Software, Supervision, Validation, Visualization, Writing - original draft. **Pedro De Podestà Uchôa de Aquino:** Investigation, Methodology, Project administration, Resources, Writing - review & editing. **Tomas Hrbek:** Formal analysis, Investigation, Methodology, Project administration, Resources, Supervision, Writing - review & editing. **Sandra Hernandez:** Formal analysis, Investigation, Methodology, Writing - review & editing. **Francisco Langeani:** Conceptualization, Funding acquisition, Investigation, Project administration, Supervision, Writing - review & editing. **Guarino R. Colli:** Conceptualization, Funding acquisition, Project administration, Resources, Supervision, Writing - review & editing. All authors have read and approved this version of the manuscript for submission.

## Acknowledgements

We are grateful to SISBIO/ICMBio for granting collecting permits supporting this work, and to Thiago B. d’Araujo Couto, Ingrid Pinheiro Paschoaletto, and Yan F. Figueira-Soares for assistance with field collections. We also thank the Virginia Commonwealth University Center for High Performance Computing (CHiPC) for providing computational resources used during the analyses. This research was funded by a Ciência Sem Fronteiras (Science Without Borders) postdoctoral fellowship from the Brazilian Conselho Nacional de Desenvolvimento Científico e Tecnológico (CNPq; Processo 314724/2014-1) to JCB and FL. GRC thanks Coordenação de Apoio à Formação de Pessoal de Nível Superior (CAPES), Conselho Nacional do Desenvolvimento Científico e Tecnológico (CNPq), Fundação de Apoio à Pesquisa do Distrito Federal (FAPDF), and the Partnerships for Enhanced Engagement in Research (PEER) program for financial support.

## Data Accessibility

Raw sequence data are deposited in the NCBI Sequence Read Archive database (BioProject XXXXXX). Sequence alignments, scripts, and Supporting Information files are available from the Zenodo repository (http://doi.org/10.5281/zenodo.16XXXX).

## Supporting Information

All supplementary materials, including Tables S1–S6, Figures S1–S8, and supporting information in Appendix S1 (sections A–C) are provided in a single external Supporting Information file (PDF).

