## Supporting Information for "Using ddRAD-seq phylogeography to test for genetic effects of headwater river capture in suckermouth armored catfish (Loricariidae: *Hypostomus*) from the central Brazilian Shield"

#### Table of Contents:


| | |
|---|---|
| **Table S1** | Pages 2 to 3 |
| **Table S2** | Pages 4 to 5 |
| **Table S3** | Pages 6 to 7 |
| **Table S4** | Pages 8 to 9 |
| **Table S5** | Pages 10 to 11 |
| **Table S6** | Page 12 |
| **Figure S1** | Page 13 |
| **Figure S2** | Page 14 |
| **Figure S3** | Page 15 |
| **Figure S4** | Page 16 |
| **Figure S5** | Page 17 |
| **Figure S6** | Page 18 |
| **Figure S7** | Page 19 |
| **Figure S8** | Page 20 |
| **Appendix S1 (A–C)** | Pages 21–29 |
| **References** | Pages 29 to 30 |

**Table S1.** Sample list, collection locality details, and summary of individual assignment to population clusters and gene tree clades.

| Taxon | ID | Field No. | Site No. | Site Code | Locality | Drainage1 | Drainage2 | Drainage3 | Basin Code | Latitude | Longitude | Elevation (ft.) | fastSTRUCTURE (Fig. 1b) | DAPC cluster (Fig. 1c) | RAxML clade (Fig. 2a) |
|---|---|---|---|---|---|---|---|---|---|---|---|---|---|---|---|
| *Hypostomus* sp. 2 | HypJCB100 | B15-09 | 6 | RdM | Córrego Retiro do Meio | Córrego Retiro do Meio | – | Río Alto São Francisco | USF | −15.630520 | −47.396450 | 2942 | 2 | 5 | 4 |
| *Hypostomus* sp. 2 | HypJCB11 | B15-01 | 1 | TA | Córrego Taquara | Río Taquara | Río Taquara | Río Alto Paraná | UP | −15.915030 | −47.911240 | 3397 | 4 | 4 | 6 |
| *Hypostomus* sp. 2 | HypJCB123 | B15-10 | 7 | RP | Ribeirão Pipiripau | Ribeirão Pipiripau | – | Río Alto Paraná | UP | −15.585700 | −47.507390 | 3441 | 2 | 5 | 4 |
| *Hypostomus* sp. 2 | HypJCB124 | B15-10 | 7 | RP | Ribeirão Pipiripau | Ribeirão Pipiripau | – | Río Alto Paraná | UP | −15.585700 | −47.507390 | 3441 | 1 | 1 | 1 |
| *Hypostomus* sp. 2 | HypJCB13 | B15-01 | 1 | TA | Córrego Taquara | Río Taquara | Río Taquara | Río Alto Paraná | UP | −15.915030 | −47.911240 | 3397 | 4 | 4 | 6 |
| *Hypostomus* sp. 2 | HypJCB134 | B15-12 | 9 | RE2 | Ribeirão Extrema site 2 | Ribeirão Extrema | Río São Bartolomeu | Río Alto São Francisco | USF | −15.787060 | −47.450170 | 2932 | 1 | 1 | 1 |
| *Hypostomus* sp. 2 | HypJCB148 | B15-14 | 10 | SdP | Córrego Saco dos Pilões | Córrego Saco dos Pilões | – | Río Alto Paraná | UP | −15.745780 | −47.665850 | 2947 | 2 | 5 | 4 |
| *Hypostomus* sp. 2 | HypJCB149 | B15-14 | 10 | SdP | Córrego Saco dos Pilões | Córrego Saco dos Pilões | – | Río Alto Paraná | UP | −15.745780 | −47.665850 | 2947 | 6 | 3 | 5 |
| *Hypostomus* sp. 2 | HypJCB15 | B15-01 | 1 | TA | Córrego Taquara | Río Taquara | Río Taquara | Río Alto Paraná | UP | −15.915030 | −47.911240 | 3397 | 4 | 4 | 6 |
| *Hypostomus* sp. 2 | HypJCB150 | B15-14 | 10 | SdP | Córrego Saco dos Pilões | Córrego Saco dos Pilões | – | Río Alto Paraná | UP | −15.745780 | −47.665850 | 2947 | 6 | 3 | 5 |
| *Hypostomus* sp. 2 | HypJCB152 | B15-14 | 10 | SdP | Córrego Saco dos Pilões | Córrego Saco dos Pilões | – | Río Alto Paraná | UP | −15.745780 | −47.665850 | 2947 | 6 | 3 | 5 |
| *Hypostomus* sp. 2 | HypJCB156 | B15-14 | 10 | SdP | Córrego Saco dos Pilões | Córrego Saco dos Pilões | – | Río Alto Paraná | UP | −15.745780 | −47.665850 | 2947 | 6 | 3 | 3 |
| *Hypostomus* sp. 2 | HypJCB161 | B15-15 | 11 | RdB | Ribeirão do Buraco | Ribeirão do Buraco | – | Río Alto Tocantins | UT | −15.601750 | −47.911170 | 2946 | 6 | 2 | 3 |
| *Hypostomus* sp. 2 | HypJCB163 | B15-15 | 11 | RdB | Ribeirão do Buraco | Ribeirão do Buraco | – | Río Alto Tocantins | UT | −15.601750 | −47.911170 | 2946 | 6 | 2 | 3 |
| *Hypostomus* sp. 2 | HypJCB167 | B15-15 | 11 | RdB | Ribeirão do Buraco | Ribeirão do Buraco | – | Río Alto Tocantins | UT | −15.601750 | −47.911170 | 2946 | 6 | 2 | 3 |
| *Hypostomus* sp. 2 | HypJCB168 | B15-15 | 11 | RdB | Ribeirão do Buraco | Ribeirão do Buraco | – | Río Alto Tocantins | UT | −15.601750 | −47.911170 | 2946 | 6 | 2 | 3 |
| *Hypostomus* sp. 2 | HypJCB17 | B15-01 | 1 | TA | Córrego Taquara | Río Taquara | Río Taquara | Río Alto Paraná | UP | −15.915030 | −47.911240 | 3397 | 4 | 4 | 6 |
| *Hypostomus* sp. 2 | HypJCB18 | B15-01 | 1 | TA | Córrego Taquara | Río Taquara | Río Taquara | Río Alto Paraná | UP | −15.915030 | −47.911240 | 3397 | 4 | 4 | 6 |
| *Hypostomus* sp. 2 | HypJCB20 | B15-01 | 1 | TA | Córrego Taquara | Río Taquara | Río Taquara | Río Alto Paraná | UP | −15.915030 | −47.911240 | 3397 | 4 | 4 | 6 |
| *Hypostomus* sp. 2 | HypJCB201 | B15-16 | 12 | RdC | Ribeirão da Contagem | Ribeirão do Buraco | – | Río Alto Tocantins | UT | −15.595920 | −47.888580 | 2821 | 6 | 2 | 3 |
| *Hypostomus* sp. 2 | HypJCB22 | B15-01 | 1 | TA | Córrego Taquara | Río Taquara | Río Taquara | Río Alto Paraná | UP | −15.915030 | −47.911240 | 3397 | 3 | 6 | 2 |
| *Hypostomus* sp. 2 | HypJCB226 | B15-27 | 8 | RE | Ribeirão Extrema | Ribeirão Extrema | Río São Bartolomeu | Río Alto São Francisco | USF | −15.849860 | −47.387030 | 2828 | 2 | 5 | 4 |
| *Hypostomus* sp. 2 | HypJCB253 | B15-26 | 9 | RE2 | Ribeirão Extrema site 2 | Ribeirão Extrema | Río São Bartolomeu | Río Alto São Francisco | USF | −15.787060 | −47.450170 | 2932 | 3 | 6 | 2 |
| *Hypostomus* sp. 2 | HypJCB254 | B15-26 | 9 | RE2 | Ribeirão Extrema site 2 | Ribeirão Extrema | Río São Bartolomeu | Río Alto São Francisco | USF | −15.787060 | −47.450170 | 2932 | 2 | 5 | 4 |
| *Hypostomus* sp. 2 | HypJCB265 | B15-21 | 14 | MC | Córrego Milho Cocido | Córrego Milho Cocido | Córrego Milho Cocido | Río Alto Paraná | UP | −15.667130 | −48.018910 | 3539 | 4 | 4 | 6 |
| *Hypostomus* sp. 2 | HypJCB42 | B15-04 | 2 | RdT | Ribeirão Do Torto | Ribeirão Do Torto | – | Río Alto Paraná | UP | −15.703430 | −47.908410 | 3387 | 3 | 6 | 2 |
| *Hypostomus* sp. 2 | HypJCB44 | B15-04 | 2 | RdT | Ribeirão Do Torto | Ribeirão Do Torto | – | Río Alto Paraná | UP | −15.703430 | −47.908410 | 3387 | 4 | 4 | 6 |

| | | | | | | | | | | | | | | | |
|---|---|---|---|---|---|---|---|---|---|---|---|---|---|---|---|
| *Hypostomus* sp. 2 | HypJCB46 | B15-04 | 2 | RdT | Ribeirão Do Torto | Ribeirão Do Torto | – | Río Alto Paraná | UP | −15.703430 | −47.908410 | 3387 | 4 | 4 | 6 |
| *Hypostomus* sp. 2 | HypJCB47 | B15-04 | 2 | RdT | Ribeirão Do Torto | Ribeirão Do Torto | – | Río Alto Paraná | UP | −15.703430 | −47.908410 | 3387 | 4 | 4 | 6 |
| *Hypostomus* sp. 2 | HypJCB49 | B15-04 | 2 | RdT | Ribeirão Do Torto | Ribeirão Do Torto | – | Río Alto Paraná | UP | −15.703430 | −47.908410 | 3387 | 4 | 4 | 6 |
| *Hypostomus* sp. 2 | HypJCB50 | B15-04 | 2 | RdT | Ribeirão Do Torto | Ribeirão Do Torto | – | Río Alto Paraná | UP | −15.703430 | −47.908410 | 3387 | 4 | 4 | 6 |
| *Hypostomus* sp. 2 | HypJCB51 | B15-04 | 2 | RdT | Ribeirão Do Torto | Ribeirão Do Torto | – | Río Alto Paraná | UP | −15.703430 | −47.908410 | 3387 | 4 | 4 | 6 |
| *Hypostomus* sp. 2 | HypJCB62 | B15-05 | 3 | RB | Ribeirão Bananal | Ribeirão Bananal | – | Río Alto Paraná | UP | −15.732000 | −47.912250 | 3372 | 3 | 6 | 2 |
| *Hypostomus* sp. 2 | HypJCB63 | B15-05 | 3 | RB | Ribeirão Bananal | Ribeirão Bananal | – | Río Alto Paraná | UP | −15.732000 | −47.912250 | 3372 | 4 | 4 | 6 |
| *Hypostomus* sp. 2 | HypJCB64 | B15-06 | 4 | U | Unnamed trib. | Unnamed trib. | – | Río Alto Tocantins | UT | −15.577880 | −47.942260 | 2931 | 3 | 6 | 2 |
| *Hypostomus* sp. 2 | HypJCB65 | B15-07 | 5 | RC | Ribeirão Cafuringa | Ribeirão Cafuringa | – | Río Alto Tocantins | UT | −15.505810 | −47.978430 | 2569 | 6 | 2 | 3 |
| *Hypostomus* sp. 2 | HypJCB66 | B15-07 | 5 | RC | Ribeirão Cafuringa | Ribeirão Cafuringa | – | Río Alto Tocantins | UT | −15.505810 | −47.978430 | 2569 | 6 | 3 | 4 |
| *Hypostomus* sp. 2 | HypJCB97 | B15-09 | 6 | RdM | Córrego Retiro do Meio | Córrego Retiro do Meio | – | Río Alto São Francisco | USF | −15.630520 | −47.396450 | 2942 | 1 | 1 | 1 |
| *Hypostomus* sp. 2 | HypJCB98 | B15-09 | 6 | RdM | Córrego Retiro do Meio | Córrego Retiro do Meio | – | Río Alto São Francisco | USF | −15.630520 | −47.396450 | 2942 | 2 | 5 | 4 |
| *Hypostomus* sp. 2 | HypJCB99 | B15-09 | 6 | RdM | Córrego Retiro do Meio | Córrego Retiro do Meio | – | Río Alto São Francisco | USF | −15.630520 | −47.396450 | 2942 | 2 | 5 | 4 |
| *Hypostomus* sp. 1 (outgroup) | HypJCB309 | B15-08 | 13 | QL | Córrego Queima Lençol | Córrego Queima Lençol | – | Río Alto Tocantins | UT | −15.546100 | −47.861470 | 2582 | 5 | 7 | Outgroup |
| *Hypostomus* sp. 1 (outgroup) | HypJCB310 | B15-08 | 13 | QL | Córrego Queima Lençol | Córrego Queima Lençol | – | Río Alto Tocantins | UT | −15.546100 | −47.861470 | 2582 | 5 | 7 | Outgroup |

Locality and drainage names are given by their actual names in Brazilian Portuguese, where possible. Potentially admixed individuals (based on fastSTRUCTURE results in Figure 1b) have their "fastSTRUCTURE" and "DAPC cluster" column entries given in red font. Abbreviated drainage basin names ("Basin Code") are based on English names for presentation in the main text. Abbreviations: sp., species (candidate species); trib., tributary; UP, Upper Paraná; USF, Upper São Francisco; UT, Upper Tocantins.

**Table S2.** Sequences of 42 DNA barcodes unique to each specimen, used for multiplexing samples in our genomic library.

| No. | Sample | Barcode sequence (5′- to -3′) |
|---|---|---|
| 1 | HypJCB100 | CCTGAGATAC |
| 2 | HypJCB11 | CTAAGGTAAC |
| 3 | HypJCB123 | TTACAACCTC |
| 4 | HypJCB124 | AACCATCCGC |
| 5 | HypJCB13 | TAAGGAGAAC |
| 6 | HypJCB134 | TCGACCACTC |
| 7 | HypJCB148 | TCTTACACAC |
| 8 | HypJCB149 | CGCATCGTTC |
| 9 | HypJCB15 | AAGAGGATTC |
| 10 | HypJCB150 | AGGAATCGTC |
| 11 | HypJCB152 | AACAATCGGC |
| 12 | HypJCB156 | TCCACTTCGC |
| 13 | HypJCB161 | AGCACGAATC |
| 14 | HypJCB163 | TCTGCCTGTC |
| 15 | HypJCB167 | TTCAATTGGC |
| 16 | HypJCB168 | CCTACTGGTC |
| 17 | HypJCB17 | CAGAAGGAAC |
| 18 | HypJCB18 | CTGCAAGTTC |
| 19 | HypJCB20 | TTCGTGATTC |
| 20 | HypJCB201 | CGATCGGTTC |
| 21 | HypJCB22 | TTCCGATAAC |
| 22 | HypJCB226 | ATCCGGAATC |
| 23 | HypJCB253 | CGAGGTTATC |
| 24 | HypJCB254 | TCCAAGCTGC |
| 25 | HypJCB265 | TTGGCATCTC |
| 26 | HypJCB42 | TGAGCGGAAC |
| 27 | HypJCB44 | CTGACCGAAC |
| 28 | HypJCB46 | TCCTCGAATC |

| | | |
|---|---|---|
| 29 | HypJCB47 | TAGGTGGTTC |
| 30 | HypJCB49 | TCTAACGGAC |
| 31 | HypJCB50 | TTGGAGTGTC |
| 32 | HypJCB51 | TCTAGAGGTC |
| 33 | HypJCB62 | TCTGGATGAC |
| 34 | HypJCB63 | TCTATTCGTC |
| 35 | HypJCB64 | AGGCAATTGC |
| 36 | HypJCB65 | CAGATCCATC |
| 37 | HypJCB66 | TCGCAATTAC |
| 38 | HypJCB97 | TTCGAGACGC |
| 39 | HypJCB98 | TGCCACGAAC |
| 40 | HypJCB99 | AACCTCATTC |
| 41 | HypJCB309 | TCAGGAATAC |
| 42 | HypJCB310 | CCTGGTTGTC |

**Table S3.** Heterozygosity (*H*) and sequencing error rates (*E*) estimated for each individual sample in this study in PyRAD v3.0.66.

| **Taxon** | **Sample** | ***H*** | ***E*** |
|---|---|---|---|
| *Hypostomus* sp. 2 | HypJCB100 | 0.00060 | $8.61 \times 10^{-5}$ |
| | HypJCB11 | 0.00031 | 0.00013 |
| | HypJCB123 | 0.00026 | $7.19 \times 10^{-5}$ |
| | HypJCB124 | 0.00024 | $4.55 \times 10^{-5}$ |
| | HypJCB13 | 0.00033 | 0.00011 |
| | HypJCB134 | 0.00021 | $2.86 \times 10^{-5}$ |
| | HypJCB148 | 0.00029 | $9.46 \times 10^{-5}$ |
| | HypJCB149 | 0.00016 | $3.67 \times 10^{-5}$ |
| | HypJCB15 | 0.00033 | $4.10 \times 10^{-5}$ |
| | HypJCB150 | 0.00088 | $9.49 \times 10^{-5}$ |
| | HypJCB152 | 0.00014 | $5.18 \times 10^{-5}$ |
| | HypJCB156 | 0.00124 | 0.00015 |
| | HypJCB161 | 0.00059 | $6.07 \times 10^{-5}$ |
| | HypJCB163 | 0.00026 | $6.11 \times 10^{-5}$ |
| | HypJCB167 | 0.00026 | $3.90 \times 10^{-5}$ |
| | HypJCB168 | 0.00046 | $7.81 \times 10^{-5}$ |
| | HypJCB17 | 0.00051 | $7.50 \times 10^{-5}$ |
| | HypJCB18 | 0.00020 | $7.72 \times 10^{-5}$ |
| | HypJCB20 | 0.00037 | $8.63 \times 10^{-5}$ |
| | HypJCB201 | 0.00025 | $7.95 \times 10^{-5}$ |
| | HypJCB22 | 0.00005 | $6.80 \times 10^{-5}$ |
| | HypJCB226 | 0.00024 | $4.47 \times 10^{-5}$ |
| | HypJCB253 | 0.00050 | 0.00019 |
| | HypJCB254 | 0.00035 | $3.38 \times 10^{-5}$ |
| | HypJCB265 | 0.00023 | $4.19 \times 10^{-5}$ |
| | HypJCB42 | 0.00031 | $9.83 \times 10^{-5}$ |
| | HypJCB44 | 0.00053 | $5.86 \times 10^{-5}$ |
| | HypJCB46 | 0.00038 | $9.79 \times 10^{-5}$ |

| | | | |
|---|---|---|---|
| | HypJCB47 | 0.00024 | 0.00011 |
| | HypJCB49 | 0.00055 | $9.51 \times 10^{-5}$ |
| | HypJCB50 | 0.00038 | 0.00011 |
| | HypJCB51 | 0.00040 | 0.0001 |
| | HypJCB62 | 0.00017 | $7.03 \times 10^{-5}$ |
| | HypJCB63 | 0.00085 | 0.00012 |
| | HypJCB64 | 0.00015 | 0.00011 |
| | HypJCB65 | 0.00038 | $7.55 \times 10^{-5}$ |
| | HypJCB66 | 0.00039 | $8.45 \times 10^{-5}$ |
| | HypJCB97 | 0.00033 | $7.22 \times 10^{-5}$ |
| | HypJCB98 | 0.00040 | $9.96 \times 10^{-5}$ |
| | HypJCB99 | 0.00020 | 0.00013 |
| *Hypostomus* sp. 1 (outgroup) | HypJCB309 | 0.00015 | 0.00017 |
| | HypJCB310 | 0.00041 | $9.32 \times 10^{-5}$ |
| Mean: | | 0.00037 | $8.46 \times 10^{-5}$ |
| Minimum: | | 0.00005 | $2.86 \times 10^{-5}$ |
| Maximum: | | 0.00124 | 0.00019 |

These parameter estimates were calculated by PyRAD from the clusters of reads at 88% sequence similarity, and with more than or equal to the minimum depth of five reads (i.e., those also summarized in columns four and five of Table 1 in the main text).

**Table S4.** Summary of G-PhoCS v1.2.3 demographic modeling results, including raw estimates of mean parameter values and their lower and upper 95% HPDs (highest posterior densities).

| | **Mean** | **Lower 95% HPD** | **Upper 95% HPD** |
|---|---|---|---|
| Data-ld-ln | −52.7396 | −59.5187 | −45.9020 |
| Full-ld-ln | −134503.9818 | −134587.9234 | −134420.7119 |
| $\theta_1$ | $5.06 \times 10^{-4}$ | $3.99 \times 10^{-4}$ | $6.10 \times 10^{-4}$ |
| $\theta_2$ | $2.81 \times 10^{-4}$ | $2.10 \times 10^{-4}$ | $3.50 \times 10^{-4}$ |
| $\theta_3$ | $6.89 \times 10^{-4}$ | $5.10 \times 10^{-4}$ | $8.50 \times 10^{-4}$ |
| $\theta_4$ | $1.37 \times 10^{-3}$ | $1.14 \times 10^{-3}$ | $1.59 \times 10^{-3}$ |
| $\theta_5$ | $3.37 \times 10^{-4}$ | $2.70 \times 10^{-4}$ | $4.00 \times 10^{-4}$ |
| $\theta_6$ | $5.11 \times 10^{-4}$ | $4.30 \times 10^{-4}$ | $5.80 \times 10^{-4}$ |
| $\theta_{56}$ | $3.31 \times 10^{-3}$ | $4.60 \times 10^{-4}$ | $6.28 \times 10^{-3}$ |
| $\theta_{23}$ | $9.82 \times 10^{-3}$ | $1.60 \times 10^{-3}$ | $1.68 \times 10^{-2}$ |
| $\theta_{2356}$ | $3.33 \times 10^{-3}$ | $3.10 \times 10^{-4}$ | $7.14 \times 10^{-3}$ |
| $\theta_{23456}$ | $1.95 \times 10^{-2}$ | $1.62 \times 10^{-2}$ | $2.28 \times 10^{-2}$ |
| $\theta_{root}$ | $1.15 \times 10^{-2}$ | $9.99 \times 10^{-3}$ | $1.30 \times 10^{-2}$ |
| $\tau_{56}$ | $5.02 \times 10^{-4}$ | $4.20 \times 10^{-4}$ | $5.80 \times 10^{-4}$ |
| $\tau_{23}$ | $4.22 \times 10^{-4}$ | $3.10 \times 10^{-4}$ | $5.60 \times 10^{-4}$ |
| $\tau_{2356}$ | $6.33 \times 10^{-4}$ | $5.00 \times 10^{-4}$ | $7.70 \times 10^{-4}$ |
| $\tau_{23456}$ | $6.38 \times 10^{-4}$ | $5.00 \times 10^{-4}$ | $7.70 \times 10^{-4}$ |
| $\tau_{root}$ | $3.63 \times 10^{-3}$ | $3.18 \times 10^{-3}$ | $4.14 \times 10^{-3}$ |
| $m_{1\rightarrow 2}$ | $5.13 \times 10^{-4}$ | $4.30 \times 10^{-4}$ | $6.00 \times 10^{-4}$ |
| $m_{1\rightarrow 3}$ | $4.01 \times 10^{-4}$ | $3.20 \times 10^{-4}$ | $4.70 \times 10^{-4}$ |
| $m_{1\rightarrow 4}$ | $1.41 \times 10^{-4}$ | $1.20 \times 10^{-4}$ | $1.70 \times 10^{-4}$ |
| $m_{1\rightarrow 5}$ | $4.03 \times 10^{-4}$ | $3.50 \times 10^{-4}$ | $4.50 \times 10^{-4}$ |
| $m_{1\rightarrow 6}$ | $4.02 \times 10^{-4}$ | $3.40 \times 10^{-4}$ | $4.70 \times 10^{-4}$ |
| $m_{2\rightarrow 1}$ | $5.60 \times 10^{-4}$ | $4.70 \times 10^{-4}$ | $6.50 \times 10^{-4}$ |
| $m_{2\rightarrow 3}$ | $5.58 \times 10^{-4}$ | $4.90 \times 10^{-4}$ | $6.20 \times 10^{-4}$ |

| | | | |
|---|---|---|---|
| $m_{2\rightarrow4}$ | $2.62 \times 10^{-4}$ | $2.20 \times 10^{-4}$ | $3.10 \times 10^{-4}$ |
| $m_{2\rightarrow5}$ | $6.15 \times 10^{-4}$ | $5.20 \times 10^{-4}$ | $7.50 \times 10^{-4}$ |
| $m_{2\rightarrow6}$ | $2.43 \times 10^{-4}$ | $2.10 \times 10^{-4}$ | $2.80 \times 10^{-4}$ |
| $m_{3\rightarrow1}$ | $4.13 \times 10^{-4}$ | $3.50 \times 10^{-4}$ | $4.70 \times 10^{-4}$ |
| $m_{3\rightarrow2}$ | $5.62 \times 10^{-4}$ | $4.90 \times 10^{-4}$ | $6.20 \times 10^{-4}$ |
| $m_{3\rightarrow4}$ | $1.83 \times 10^{-4}$ | $1.50 \times 10^{-4}$ | $2.20 \times 10^{-4}$ |
| $m_{3\rightarrow5}$ | $1.80 \times 10^{-4}$ | $1.50 \times 10^{-4}$ | $2.20 \times 10^{-4}$ |
| $m_{3\rightarrow6}$ | $4.98 \times 10^{-4}$ | $4.20 \times 10^{-4}$ | $5.80 \times 10^{-4}$ |
| $m_{4\rightarrow1}$ | $1.40 \times 10^{-4}$ | $1.10 \times 10^{-4}$ | $1.60 \times 10^{-4}$ |
| $m_{4\rightarrow2}$ | $2.37 \times 10^{-4}$ | $2.00 \times 10^{-4}$ | $2.80 \times 10^{-4}$ |
| $m_{4\rightarrow3}$ | $2.09 \times 10^{-4}$ | $1.70 \times 10^{-4}$ | $2.60 \times 10^{-4}$ |
| $m_{4\rightarrow5}$ | $2.56 \times 10^{-4}$ | $2.20 \times 10^{-4}$ | $3.00 \times 10^{-4}$ |
| $m_{4\rightarrow6}$ | $1.60 \times 10^{-4}$ | $1.40 \times 10^{-4}$ | $1.80 \times 10^{-4}$ |
| $m_{5\rightarrow1}$ | $4.18 \times 10^{-4}$ | $3.70 \times 10^{-4}$ | $4.70 \times 10^{-4}$ |
| $m_{5\rightarrow2}$ | $6.83 \times 10^{-4}$ | $5.70 \times 10^{-4}$ | $8.30 \times 10^{-4}$ |
| $m_{5\rightarrow3}$ | $1.68 \times 10^{-4}$ | $1.40 \times 10^{-4}$ | $2.10 \times 10^{-4}$ |
| $m_{5\rightarrow4}$ | $2.69 \times 10^{-4}$ | $2.30 \times 10^{-4}$ | $3.10 \times 10^{-4}$ |
| $m_{5\rightarrow6}$ | $4.56 \times 10^{-4}$ | $3.70 \times 10^{-4}$ | $5.30 \times 10^{-4}$ |
| $m_{6\rightarrow1}$ | $4.33 \times 10^{-4}$ | $3.60 \times 10^{-4}$ | $5.00 \times 10^{-4}$ |
| $m_{6\rightarrow2}$ | $2.45 \times 10^{-4}$ | $2.10 \times 10^{-4}$ | $2.80 \times 10^{-4}$ |
| $m_{6\rightarrow3}$ | $4.50 \times 10^{-4}$ | $3.80 \times 10^{-4}$ | $5.20 \times 10^{-4}$ |
| $m_{6\rightarrow4}$ | $1.46 \times 10^{-4}$ | $1.30 \times 10^{-4}$ | $1.70 \times 10^{-4}$ |
| $m_{6\rightarrow5}$ | $5.00 \times 10^{-4}$ | $4.10 \times 10^{-4}$ | $5.80 \times 10^{-4}$ |

Log-likelihoods and $\theta$ and $\tau$ results are presented as averages over 15 independent runs specifying different pairwise migration bands. Parameter subscript numbers correspond to clades in the species tree; that is, number labels 1–6 indicate *Hypostomus* genetic clusters 1–6 discussed in the main text. The summarized log-likelihoods included the data likelihood (‘Data-ld-ln’) and posterior probability (‘Full-ld-ln’). Each of the 15 pairs of migration rate estimates ($m_{sx}$, $m_{xs}$) is reported from an independent run using a different migration band. The dates of putative migration events (bands) were not estimated. Units are discussed in section 2 (Material and methods) of the main text.

**Table S5.** Summary of converted G-PhoCS parameter estimates.

| | Mean | Lower 95% HPD | Upper 95% HPD |
|---|---|---|---|
| $N_{e1}$ | 43,620.69 | 34,396.55 | 52,586.21 |
| $N_{e2}$ | 24,224.14 | 18,103.45 | 30,172.41 |
| $N_{e3}$ | 59,396.55 | 43,965.52 | 73,275.86 |
| $N_{e4}$ | 118,103.45 | 98,275.86 | 137,068.97 |
| $N_{e5}$ | 29,051.72 | 23,275.86 | 34,482.76 |
| $N_{e6}$ | 44,051.72 | 37,068.97 | 50,000.00 |
| $N_{e56}$ | 285,344.83 | 39,655.17 | 541,379.31 |
| $N_{e23}$ | 846,551.72 | 137,931.03 | 1,448,275.86 |
| $N_{e2356}$ | 287,068.97 | 26,724.14 | 615,517.24 |
| $N_{e23456}$ | 1,681,034.48 | 1,396,551.72 | 1,965,517.24 |
| $N_{e\ root}$ | 991,379.31 | 861,206.90 | 1,120,689.66 |
| $T_{56}$ | 173,103.45 | 36,206.90 | 200,000.00 |
| $T_{23}$ | 145,517.24 | 26,724.14 | 193,103.45 |
| $T_{2356}$ | 218,275.86 | 43,103.45 | 265,517.24 |
| $T_{23456}$ | 220,000.00 | 43,103.45 | 265,517.24 |
| $T_{root}$ | 1,251,724.14 | 274,137.93 | 1,427,586.21 |
| $M_{1\rightarrow2}$ | $3.60 \times 10^{-8}$ | $2.26 \times 10^{-8}$ | $5.25 \times 10^{-8}$ |
| $M_{1\rightarrow3}$ | $6.91 \times 10^{-8}$ | $4.08 \times 10^{-8}$ | $9.99 \times 10^{-8}$ |
| $M_{1\rightarrow4}$ | $4.83 \times 10^{-8}$ | $3.42 \times 10^{-8}$ | $6.76 \times 10^{-8}$ |
| $M_{1\rightarrow5}$ | $3.40 \times 10^{-8}$ | $2.36 \times 10^{-8}$ | $4.50 \times 10^{-8}$ |
| $M_{1\rightarrow6}$ | $5.14 \times 10^{-8}$ | $3.66 \times 10^{-8}$ | $6.82 \times 10^{-8}$ |
| $M_{2\rightarrow1}$ | $7.08 \times 10^{-8}$ | $4.69 \times 10^{-8}$ | $9.91 \times 10^{-8}$ |
| $M_{2\rightarrow3}$ | $9.61 \times 10^{-8}$ | $6.25 \times 10^{-8}$ | $1.32 \times 10^{-7}$ |
| $M_{2\rightarrow4}$ | $8.97 \times 10^{-8}$ | $6.27 \times 10^{-8}$ | $1.23 \times 10^{-7}$ |
| $M_{2\rightarrow5}$ | $5.18 \times 10^{-8}$ | $3.51 \times 10^{-8}$ | $7.50 \times 10^{-8}$ |

| | | | |
|---|---|---|---|
| $M_{2\to6}$ | $3.10 \times 10^{-8}$ | $2.26 \times 10^{-8}$ | $4.06 \times 10^{-8}$ |
| $M_{3\to1}$ | $5.22 \times 10^{-8}$ | $3.49 \times 10^{-8}$ | $7.17 \times 10^{-8}$ |
| $M_{3\to2}$ | $3.95 \times 10^{-8}$ | $2.57 \times 10^{-8}$ | $5.43 \times 10^{-8}$ |
| $M_{3\to4}$ | $6.27 \times 10^{-8}$ | $4.28 \times 10^{-8}$ | $8.75 \times 10^{-8}$ |
| $M_{3\to5}$ | $1.52 \times 10^{-8}$ | $1.01 \times 10^{-8}$ | $2.20 \times 10^{-8}$ |
| $M_{3\to6}$ | $6.36 \times 10^{-8}$ | $4.52 \times 10^{-8}$ | $8.41 \times 10^{-8}$ |
| $M_{4\to1}$ | $1.77 \times 10^{-8}$ | $1.10 \times 10^{-8}$ | $2.44 \times 10^{-8}$ |
| $M_{4\to2}$ | $1.66 \times 10^{-8}$ | $1.05 \times 10^{-8}$ | $2.45 \times 10^{-8}$ |
| $M_{4\to3}$ | $3.60 \times 10^{-8}$ | $2.17 \times 10^{-8}$ | $5.53 \times 10^{-8}$ |
| $M_{4\to5}$ | $2.16 \times 10^{-8}$ | $1.49 \times 10^{-8}$ | $3.00 \times 10^{-8}$ |
| $M_{4\to6}$ | $2.04 \times 10^{-8}$ | $1.51 \times 10^{-8}$ | $2.61 \times 10^{-8}$ |
| $M_{5\to1}$ | 5.29 E-08 | $3.69 \times 10^{-8}$ | $7.17 \times 10^{-8}$ |
| $M_{5\to2}$ | 4.80 E-08 | $2.99 \times 10^{-8}$ | $7.26 \times 10^{-8}$ |
| $M_{5\to3}$ | 2.89 E-08 | $1.79 \times 10^{-8}$ | $4.46 \times 10^{-8}$ |
| $M_{5\to4}$ | 9.21 E-08 | $6.56 \times 10^{-8}$ | $1.23 \times 10^{-7}$ |
| $M_{5\to6}$ | 5.83 E-08 | $3.98 \times 10^{-8}$ | $7.69 \times 10^{-8}$ |
| $M_{6\to1}$ | 5.48 E-08 | $3.59 \times 10^{-8}$ | $7.63 \times 10^{-8}$ |
| $M_{6\to2}$ | 1.72 E-08 | $1.10 \times 10^{-8}$ | $2.45 \times 10^{-8}$ |
| $M_{6\to3}$ | 7.75 E-08 | $4.85 \times 10^{-8}$ | $1.11 \times 10^{-7}$ |
| $M_{6\to4}$ | 5.00 E-08 | $3.71 \times 10^{-8}$ | $6.76 \times 10^{-8}$ |
| $M_{6\to5}$ | 4.21 E-08 | $2.77 \times 10^{-8}$ | $5.80 \times 10^{-8}$ |

Shown are values of demographic parameters for each lineage ($x$; *Hypostomus* genetic clusters 1–6 discussed in the main text), including mean $N_{ex}$ (converted from $\theta_x$ in Table S4), mean $T$ (converted from $\tau$ in Table S4), and their lower and upper 95% HPDs, from G-PhoCS. Results from converting the migration rate ($m_{sx}$) into the migration rate per generation ($M_{sx} = m_{sx} \times \theta_x/4$) are also shown, and $M_{sx}$ represents the is the proportion of individuals in population $x$ that arrived by migration from population $s$ per generation (interpretation: $\approx 2Nm_{sx}$); these correspond to effectively zero total migration rates. Results are presented as averages over 15 independent runs specifying different pairwise migration bands. Parameter subscript numbers correspond to clades in the species tree. Units: $N_e$ , number of breeding individuals; $M_{sx}$, proportion of migrant individuals; $T$, absolute time in years. The mean $N_{ex}$ values across all clusters sum to *c.* ~318,448 breeding individuals.

**Table S6.** Summary of population size estimates from Bayesian analysis of our SNP dataset to infer a species tree under the multispecies coalescent model in SNAPP (Bryant et al., 2012).

| | **Raw $\theta_x$ values** | | | **$N_e$ estimates** | | |
|---|---|---|---|---|---|---|
| | **Mean** | **Lower 95% HPD** | **Upper 95% HPD** | **Mean** | **Lower 95% HPD** | **Upper 95% HPD** |
| Cluster 1 ($\theta_5$) | $4.63 \times 10^{-4}$ | $2.00 \times 10^{-4}$ | $1.16 \times 10^{-3}$ | 39,971.78 | 17,259.40 | 99,756.29 |
| Cluster 2 ($\theta_4$) | $2.15 \times 10^{-2}$ | $9.84 \times 10^{-3}$ | $3.56 \times 10^{-2}$ | 1,851,644.83 | 848,060.26 | 3,065,941.38 |
| Cluster 3 ($\theta_1$) | $1.47 \times 10^{-1}$ | $8.25 \times 10^{-2}$ | $2.18 \times 10^{-1}$ | 12,665,982.76 | 7,110,139.66 | 18,828,853.45 |
| Cluster 4 ($\theta_3$) | $1.06 \times 10^{-1}$ | $3.07 \times 10^{-3}$ | $1.97 \times 10^{-2}$ | 917,425.86 | 264,388.45 | 1,698,193.97 |
| Cluster 5 ($\theta_0$) | $1.31 \times 10^{-1}$ | $4.66 \times 10^{-3}$ | $2.25 \times 10^{-2}$ | 1,133,265.52 | 401,907.84 | 1,938,948.28 |
| Cluster 6 ($\theta_2$) | $7.40 \times 10^{-2}$ | $1.74 \times 10^{-3}$ | $1.50 \times 10^{-2}$ | 640,161.90 | 149,980.34 | 1,293,204.31 |
| Internal branch 1 ($\theta_6$) | $1.71 \times 10^{-3}$ | $2.00 \times 10^{-4}$ | $4.64 \times 10^{-3}$ | 147,446.55 | 17,254.57 | 399,896.12 |
| Internal branch 2 ($\theta_7$) | $2.65 \times 10^{-3}$ | $2.00 \times 10^{-4}$ | $4.13 \times 10^{-3}$ | 228,416.38 | 17,263.19 | 355,875.95 |
| Internal branch 3 ($\theta_8$) | $1.72 \times 10^{-3}$ | $2.00 \times 10^{-4}$ | $5.21 \times 10^{-3}$ | 148,262.33 | 17,270.09 | 449,198.71 |
| Internal branch 4 ($\theta_9$) | $2.43 \times 10^{-2}$ | $2.00 \times 10^{-4}$ | $4.84 \times 10^{-3}$ | 2,095,884.48 | 17,247.84 | 417,136.55 |
| Internal branch 5 ($\theta_{10}$) | $2.54 \times 10^{-3}$ | $2.00 \times 10^{-4}$ | $4.41 \times 10^{-3}$ | 218,866.72 | 17,255.00 | 379,963.62 |

Shown at left (columns 2–4) are values of the population mutation rate parameter ($\theta_x$) for each tip lineage ($x$) and internal branch lineage in our SNAPP species tree (Figure 2c), given as means and their lower and upper 95% HPDs from SNAPP. Columns 5–7 present mean $N_{ex}$ (converted from $\theta_x$s at left) and their lower and upper 95% HPDs. Parameter subscript numbers correspond to genetic cluster lineages in the species tree. Population size estimates for *Hypostomus* cluster 3 were much larger, in some cases by orders of magnitude, than those for other clusters. Units: $\theta_x$, coalescent units; $N_e$, number of breeding individuals.

**Figure S1.** Plot of Bayesian information criterion (BIC) values over the range of numbers of *K* clusters analyzed during DAPC, supporting *K* = 7 as the levelling-off point. This value included six ingroup clusters and one outgroup cluster.

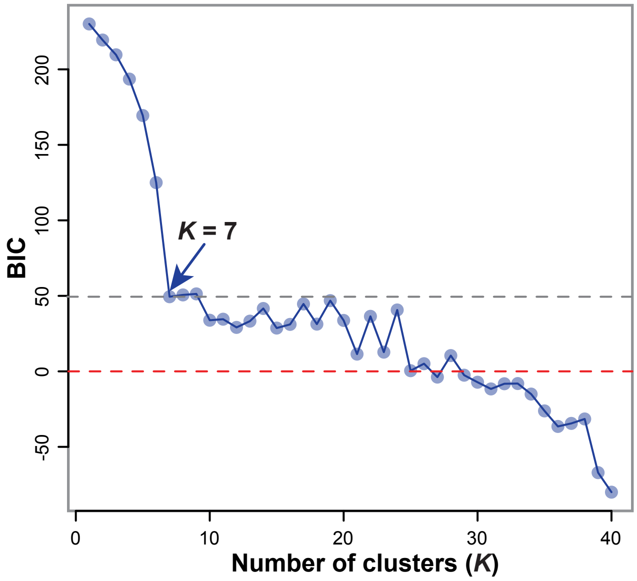

**Figure S2.** Bayesian 50% majority-rule consensus tree from partitioned analysis of the ddRAD-seq dataset in MrBayes v3.2. Nodal support values are Bayesian posterior probabilities, represented by closed or open circles (see legend).

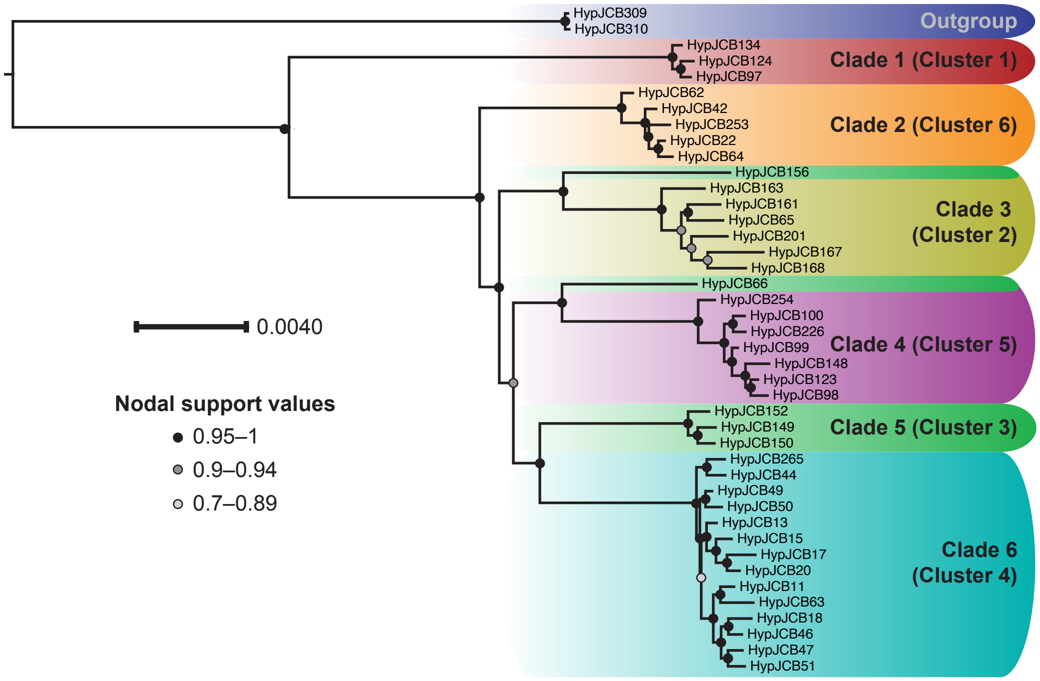

**Figure S3.** DensiTree plot of 5000 posterior (post-burn-in) trees from species tree analysis in SNAPP (Bryant et al., 2012). Shown in thick black lines is the final (~best) MCC tree from the SNAPP analysis.

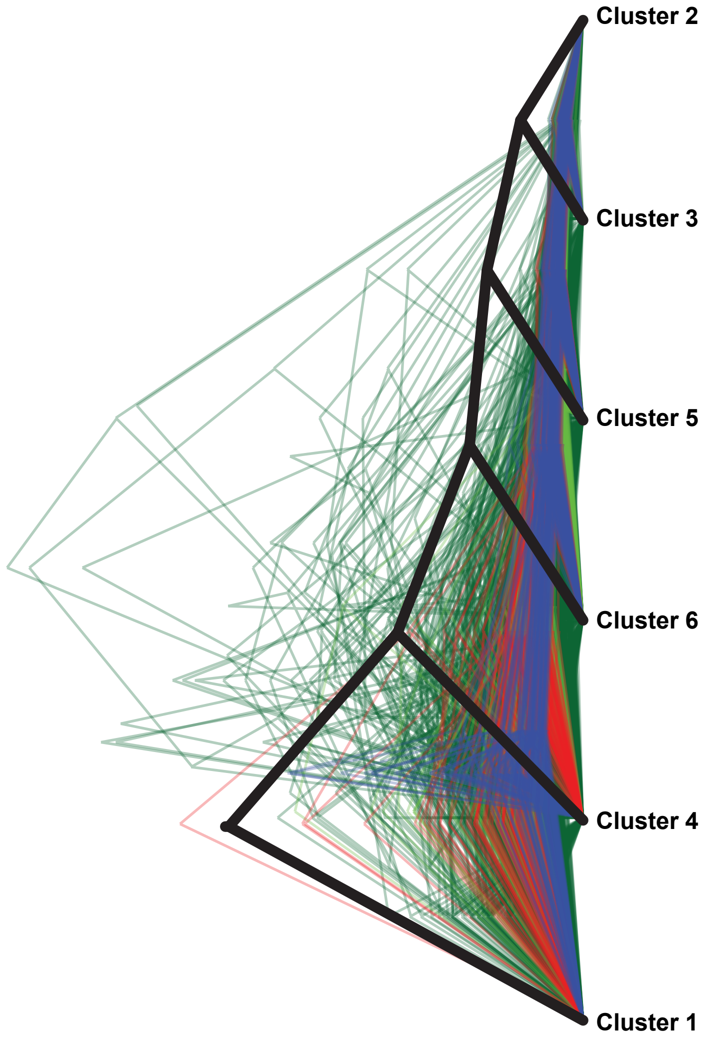

**Figure S4.** SVDquartets gene tree estimated from quartets under the multispecies coalescent model. Nodal support values are bootstrap proportions from 500 bootstrap pseudoreplicates in SVDquartets.

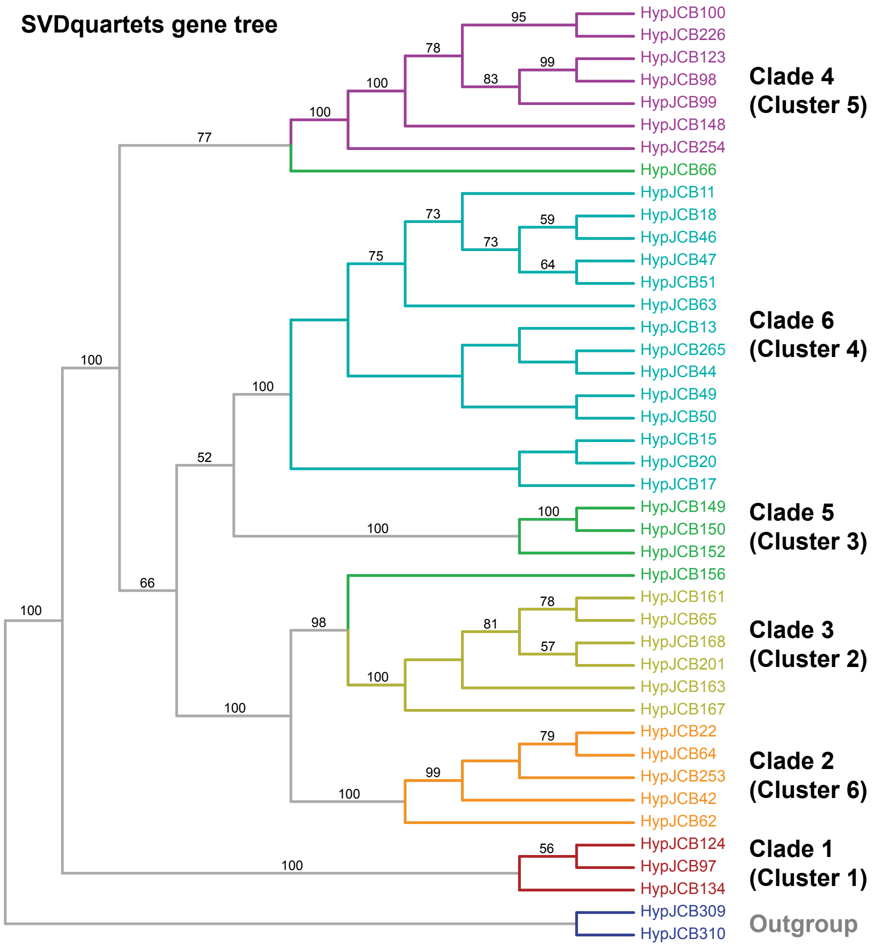

**Figure S5.** Multilocus phylogenetic network computed from *p*-distances between all sequences in SplitsTree v4.13.1 (Huson and Bryant, 2006) using the Neighbor-Net approach. Circles at the tips of the network represent individual samples (multilocus genotypes), and clade colors correspond to genetic clusters in Figure 1c.

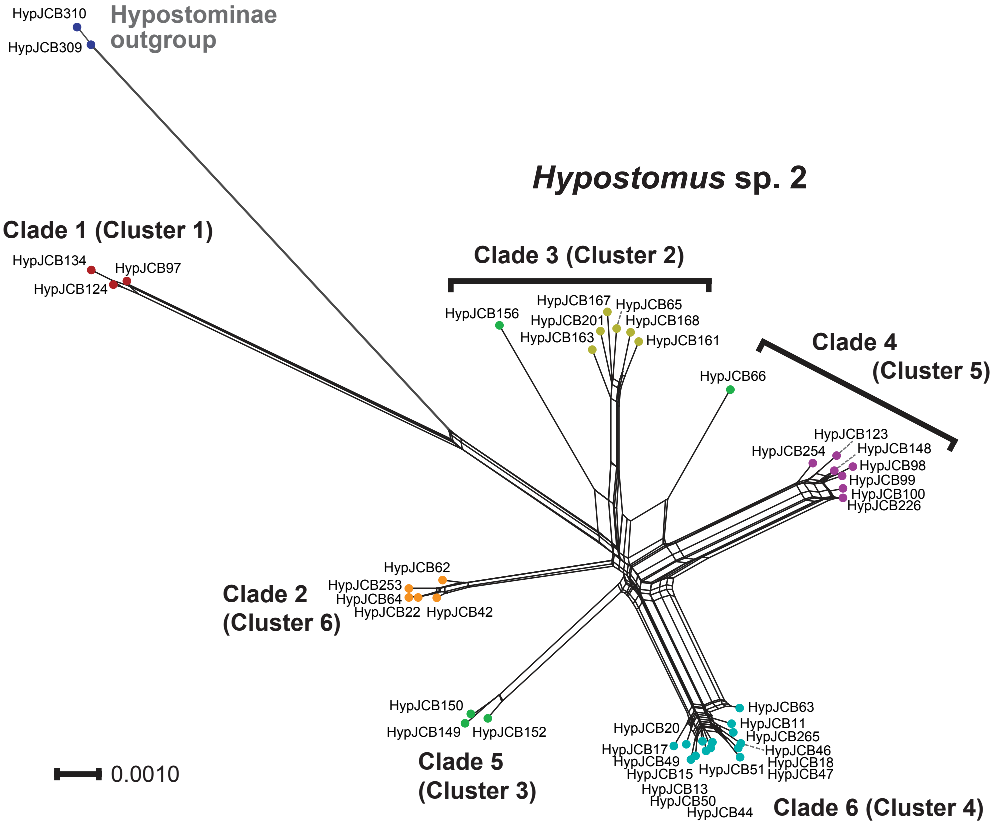

**Figure S6.** Species and population size estimates inferred during analysis of *Hypostomus* sequence data in SNAPP (Bryant et al., 2012). Internal branch widths (black lines) are scaled to effective population sizes ($N_e$) of the genetic clusters/lineages at the tips of the tree; raw ($\theta_x$) and converted values of these parameters are given in Table S6. Nodal support values are raw Bayesian posterior probabilities (BPP); values ≥ 0.95 were considered significant in this study.

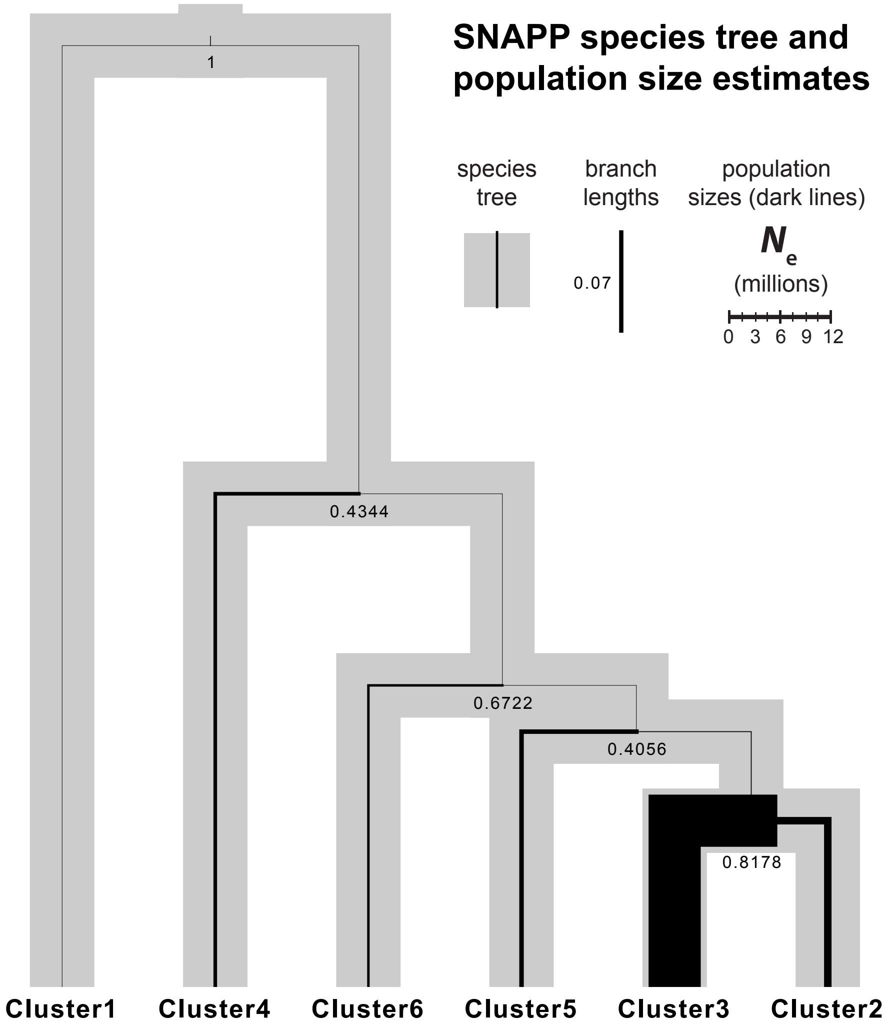

**Figure S7.** Gene tree and divergence times from Bayesian discrete trait diffusion analysis (Lemey et al., 2009) in BEAST. Tip labels contain the sample IDs, followed by latitude and longitude coordinates of the corresponding sampling location (given in decimal degrees), the two- or three-letter drainage code (see legend), and a final drainage code including a site number. Here, the site number is not the same as that in Table S1, but is included to differentiate samples from different sites within the same drainage basin. As in Figure 2, clade colors correspond to genetic clusters in Figure 1c (see legend, below left).

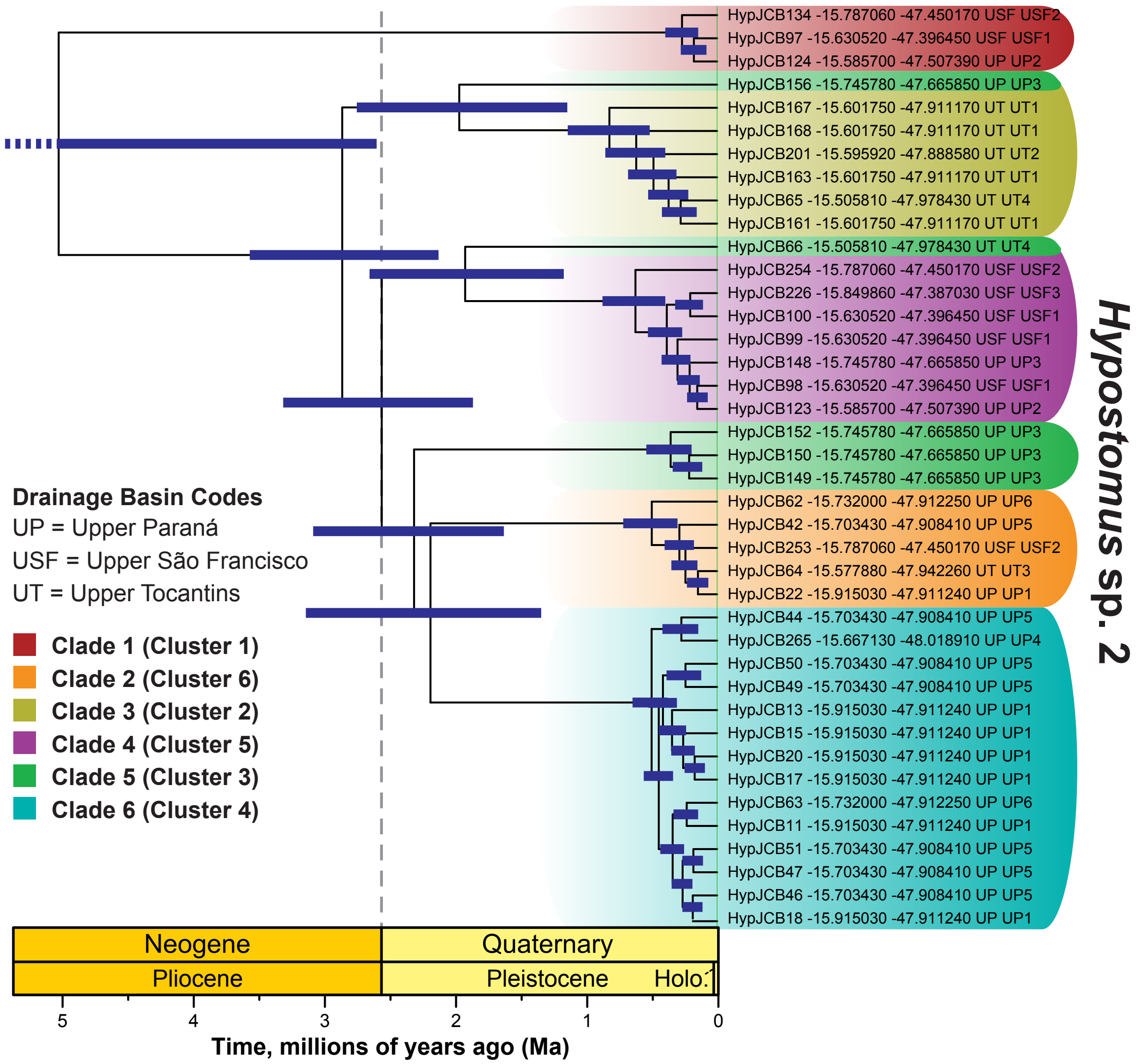

**Figure S8.** Graphical summary of Table 1 predictions of our two main hypotheses, the ‘Paraná Capture Hypothesis’ and the non-mutually exclusive ‘Frequent Interdrainage Dispersal Hypothesis’ (derived from Aquino and Colli, 2017), that were supported (green check marks) versus rejected (red crosses) by our results. See main text and Appendix S1C for additional details and interpretations.

**Legend**

Prediction(s) rejected

Prediction(s) supported

***Note:*** *Assessments based on evidence from “Phylogenetic gene tree and species tree reconstructions” (under “Methods”) are included three times in the table. However, evidence for or against a hypothesis is only counted once for each analysis/method. Thus, three green check marks for ”gene tree and species tree” evidence only count as one. Taking such patterns into account, a total of 9/11 (82%) of analyses/ methods supported the Paraná Capture Hypothesis, while only 2/6 (33%) supported the Frequent Interdrainage Dispersal Hypothesis.*

| Predictions | ‘Paraná Capture Hypothesis’ | ‘Frequent Interdrainage Dispersal Hypothesis’ | Methods |
|---|---|---|---|
| (1) Phylogenetic divergence, overall phylogeographic structure | Phylogeographic structure present; genetic divergences at drainage divide(s); Upper Tocantins (UT) or Upper São Francisco (USF) captured-population alleles sister to, or nested within, clades containing Upper Paraná (UP) source alleles | Low/non-existent phylogeographic structure; para-/polyphyletic lineages result from gene flow following secondary contact | • ‘Phylogeographic association test’ (spatial randomization)<br>• Phylogenetic gene tree and species tree reconstructions<br>• Multilocus phylogenetic network |
| (2) Genealogical sorting | Patterns of *gsi* or $gsi_T$ for samples grouped by drainages as follows:<br>• configuration 1: UP+UT vs. USF (significant)<br>• configuration 2: UP+USF vs. UT (significant)<br>• configuration 3: UP vs. USF vs. UT (non-significant overall, or UP *gsi* non-significant) | N/A | • Genealogical sorting index (single-locus *gsi*, multilocus ‘ensemble’ $gsi_T$) tests of degree of gene/allele sorting of between drainages or drainage groups, based on individual gene trees |
| (3) Ancestral populations located in | Ancestral geographical location in UP basin | N/A | • Phylogenetic gene tree and species tree reconstructions<br>• Spatial random walk model of migration (continuous phylogeography); ‘center test’ (spatial randomization)<br>• Bayesian discrete trait diffusion analysis (by drainage basin) |
| (4) Derived (e.g. tip-positioned) populations located in | UT or USF river basins | Multiple basins | • Phylogenetic gene tree and species tree reconstructions |
| (5) Population history includes | Founder effects (smaller $N_e$) or recent population expansion in UT basin | Migration resulting in gene flow; evidence of admixture between lineages and basins | • Coalescent-based demographic modeling |
| (6) Spatial dispersal includes | Non-random dispersal | Random dispersal | • ‘Overall directionality test’ (spatial randomization) |
| | Directional dispersal, from UP basin into UT basin | Multidirectional dispersal between basins (no single prevailing direction) | • ‘A priori direction test’ (spatial randomization) |

### Appendix S1.

#### A. Supplementary Materials and Methods

***DNA extraction, library preparation, and sequencing*—**We extracted whole genomic DNA from tissue samples using Invitrogen PureLink Genomic Purification kits. In the final extraction step, DNA was eluted from spin columns using two consecutive steps, each consisting of a 2 min incubation in 75–100 µL of elution buffer followed by a 1 min spin at 14,000×*g*. We confirmed the presence of high molecular weight genomic DNA from each specimen on 1% agarose gels and then quantified sample DNA concentration and purity using a NanoDrop 2000c Spectrophotometer (Thermo Scientific). Samples with < 10 ng/µL of DNA were dried down for 30 min on a vacuum centrifuge and then resuspended in ~50 µL of water that had been previously deionized and purified using a water filtration station.

We prepared a double-digest restriction site-associated DNA sequencing (ddRAD-seq) library for our samples using the method of Peterson et al. (2012), adapted for the Ion Torrent platform (https://github.com/legalLab/protocols-scripts). First, we conducted a simultaneous digestion/ligation polymerase chain reaction (PCR) starting from 200 ng of DNA from each sample. This step digested the DNA using restriction enzymes *SdaI* (8-cutter enzyme, `TGCA`-3′ overhang) and *Csp6I* (cuts `GTAC` with 5′-`TA` overhang), and ligated adapters A*Y* (where A is 5′-`CCATCTCATCCCTGCGTGTC`-3′, and *Y* is one of 42 barcode primers unique to each specimen, listed in Table S2) and P1 (5′-`CCTCTCTATGGGCAGTCGGTGAT`-3′). After confirming high adapter incorporation on 1% agarose gels, we conducted an additional, nested digestion PCR using four 15-cycle replicates per sample, starting from 1 µL of product from the initial digestion/ligation. Afterwards, four replicates for each sample were combined, and we arrived at fluorometric estimates of DNA concentrations of the samples using a Qubit dsDNA HS Assay Kit (Life Technologies). Barcoded libraries from each of the individual specimens were pooled into a single tube in an equimolar fashion. We concentrated the library by splitting it equally among three 1.5 mL tubes and drying it down for 1 h in a vacuum centrifuge system; resuspending the contents of each tube in 100 µL of filtered, deionized water; and then re-pooling the three aliquots into a single tube. We purified the library using Agencourt AMPure XP beads and the manufacturer's protocol (Beckman Coulter), finishing with an elution in 60 µL of purified water.

Previous ddRAD-seq studies have shown that automated technology is superior to manual DNA size selection (e.g. Peterson et al., 2012). We conducted size selection on the pooled library by selecting 415 bp fragments (range = 374–456 bp; including adapter sequences) using a Pippin Prep procedure with 2% DF Marker L agarose gel cassettes (Sage Science). Following size selection, we conducted a second AMPure bead purification of the pooled library, with final elution in 23 µL of purified water. After emulsion PCR, the final library was sequenced to generate millions of single-end reads on an Ion Torrent machine, as described in the main text. The raw reads have been deposited in the National Center for Biotechnology Information (NCBI) Short Read Archive (SRA; http://www.ncbi.nlm.nih.gov/sra; SRA data PRJNAXXXXXX).

***De novo assembly and SNP processing*—**We summarized potential levels of paralogous loci in our data, which were flagged and excluded by PyRAD during paralog detection and filtering in PyRAD steps 5 and 7 (Eaton, 2014), in two ways. First, the number of flagged paralogs was calculated for each individual sample as f1loci − f2loci based on statistics in the 'S5consens.txt' file output by our empirical PyRAD run. The proportion of flagged paralogs was calculated from the same summary statistics as [f1loci − f2loci] / f1loci, and these statistics were also converted to percentage values. Descriptive statistics (minimum, mean, and maximum) for putative paralog and ortholog loci were calculated across individuals. The '*.excluded_loci' file output by PyRAD also provides opportunities for counting the number of putatively paralogous SNPs (from potentially paralogous gene regions) that were excluded from the dataset during a given assembly (contained within the excluded ddRAD tag sequences). For our empirical assembly, we assessed this value by counting the number of non-parsimony informative and parsimony-informative SNP locations in the '*.excluded_loci' file output by PyRAD, and calculating the proportion of this value into the total number of SNPs.

***Population structure and admixture***—As noted in the text, we inferred population structure and genomic admixture using two different methods, including fastSTRUCTURE and DAPC as implemented in adegenet. Here, we provide more information on these methods and their complementarity, as well as additional justification for preferring our DAPC clustering results.

First, fastSTRUCTURE (Raj et al., 2014) has the advantages of being flexible and allowing individuals to have genomic ancestry from multiple populations, and to infer this pattern of admixture when present. Additionally, fastSTRUCTURE is orders of magnitude faster than the similar assignment software program STRUCTURE (Pritchard et al., 2000).

Second, while DAPC does not infer mixed ancestry proportions (which would otherwise be a drawback), the *k*-means approach used in DAPC has the advantages of being incredibly fast, completed in a matter of seconds, and of being highly accurate for genomic data of known 'dosage' such as genotyping-by-sequencing data (Stift et al., 2019). Indeed, a recent study comparing the accuracy of the software programs STRUCTURE, fastSTRUCTURE, and ADMIXTURE (Alexander et al., 2009), as well as the *k*-means algorithm used in DAPC, showed that *k*-means clustering is just as accurate as STRUCTURE (which was the most accurate method overall) for codominant data over a range of levels of population differentiation (Stift et al., 2019). As for how DAPC works, given a small number of linear combinations of the original allelic variation in the SNP data, constructed as principal components (PCs), the DAPC approach estimates group membership probabilities for individuals using discriminant analysis of the PCs (Jombart et al., 2010). The main drawback we see with DAPC, recently also iterated by Stift et al. (2019), is the possibility of spurious clustering by ploidy, but this is highly unlikely to have affected our results, as all cytogenetic information on loricariid catfishes to date indicates that they are diploid (e.g. de Oliveira et al., 2019). Such issues are most likely to arise in the analysis of seed plants (e.g. angiosperms) or other taxa that are known to exhibit widespread and high degrees of variation in ploidy due to polyploidization events in their recent or ancient past.

***Phylogenomic gene tree and species tree analyses***—Here, we briefly provide additional details on our phylogenetic methods for which there was insufficient space in the main text. Regarding our gene tree analyses, we note that RAxML runs used a stopping rule to determine when convergence had been reached and output a single 'best' tree. This stopping rule is determined by the program.

The evolutionary histories of populations, loci, or genome regions may not always follow a strictly bifurcating path such as that represented in phylogenetic trees, due to hybridization, horizontal gene transfer, or recombination (e.g. Bryant and Moulton, 2004). By contrast, phylogenetic networks can infer reticulate histories of the genomes of organisms, hence overcoming inherent limitations of traditional phylogenies. Phylogenetic networks also can help identify sequencing errors or other issues with samples. As noted in the main text, we inferred a phylogenetic network from the full SNP dataset using the Neighbor-Net method implemented in SplitsTree v4.13.1 (Huson and Bryant, 2006), based on uncorrected nucleotide distances (*p*-distances) among samples. Boxes in the resulting network are interpreted as uncertainty in topological relationships due to confounding factors or factors that cause homoplasy (identity by state not due to shared ancestry; character states that evolve independently), including gene flow, incomplete lineage sorting, or recombination (Huson and Bryant, 2006). Often in SplitsTree networks, when one lineage is comprised of one or a few individuals and is intermediate between two well-established lineages that are major clades in regular phylogenetic gene tree or species tree analyses, researchers assume that those individuals may be hybrids or individuals of admixed ancestry. SplitsTree networks can also be useful in visualizing character conflict, and may indicate taxa to test for removal as candidates for rogue taxa or long-branch attraction, although we did not employ such methods in our study.

***Phylogeographic analyses*****—**Here, we briefly provide additional background information on our phylogeographic methods. In particular, we note that PhyloMapper estimates the geographical location of the ancestor of a clade from geographical coordinates of collection localities for each tip in its phylogeny, based on a spatially explicit random walk model of migration estimated using a ML algorithm (Lemmon and Lemmon, 2008).

***Effects of varying ddRAD-seq assembly parameters on downstream genetic analyses*****—** Regarding our sensitivity analyses, our abbreviations of PyRAD (Eaton, 2014) run parameters vary slightly from the descriptions and abbreviations used in the PyRAD v3+ documentation and Eaton (2014) paper describing the software program. To clarify our choice of abbreviations for these parameters, our abbreviations mC, SH, pO, mD, and cP are the same as 'MinCov', 'MaxSH', 'phred Qscore offset', 'Mindepth', and 'Wclust', respectively, in the defalut PyRAD 'params.txt' file (parameter settings file). This default params.txt file may be generated using PyRAD v3.0.66 by entering `pyrad -n` at the command line interface, as noted in the README file in the original PyRAD GitHub repository (https://github.com/dereneaton/pyrad). Additionally, our parameter abbreviation SH refers to the same parameter as 'maxSharedH' mentioned in Eaton (2014).

One important parameter in our PyRAD analyses, cP, is analogous to the "distance allowed between stacks" setting (`-M` flag) in Stacks (Catchen et al., 2011, 2013). The default PyRAD setting of 85% (0.85) means that sequences will be clustered based on 85% sequence similarity, which is equivalent to 15 base differences between 100-bp reads (Eaton, 2014). The analogous settings in Stacks would be the following parameters: `-m6`, `-M15`. By contrast, setting `-M4` in Stacks ould be analogous to cp = 96% in PyRAD (Catchen et al., 2011, 2013; McCartney-Melstad et al., 2019).

#### B. Supplementary Results

***DNA extraction, library preparation, and sequencing*****—**To supplement our presentation of results in the main text, we first note that checks confirmed that, after filtering out low-quality ISPs (ion sphere particles), ~55.8–62.9% of library ISPs carried template DNA during Ion Torrent sequencing (two runs, using two different 318 chips). The moderate ISP enrichment values we obtained were likely due to polyclonal ISPs. This is a relatively common issue for Ion

Torrent library preps and may have resulted from poor emulsion during emulsion PCR, or more likely an excess of library over beads when plating. However, values were typical for the machine and laboratory used, and similar results were obtained in several recently generated datasets for taxonomically diverse groups of Brazilian fishes in the lab (T. Hrbek, unpublished data). Additionally, despite any ISP issues, we still obtained excellent amounts of data with high phylogenetic information content for our analyses (see main text for details).

***De novo assembly and SNP processing*****—**Regarding the de novo assembly of our ddRAD-seq data in PyRAD, we note that, among the consensus loci mentioned in the text as passing paralogy filtering, the mean heterozygosity per individual ($3.69 \times 10^{-4}$) was approximately 4-fold greater than the mean error rate ($8.46 \times 10^{-5}$) (Table S3). These values are used to calculate the binomial probability of homozygosity versus heterozygosity at each site, and this information is then used to determine whether there is sufficient statistical support for making a base call at the site (Eaton, 2014).

During the final empirical assembly steps in PyRAD, putative paralog sequences are excluded when they contain repetitive DNA or have high copy number (e.g. read depth) sequence regions (Eaton, 2014). In PyRAD, as well as the Stacks pipeline (Catchen et al., 2011), the number of putative paralogs that is excluded during ddRAD-seq data assembly has been shown to be negatively correlated with clustering percentage threshold values (McCartney-Melstad et al., 2019). High levels of paralogy, e.g. leading to ~20% or more loci being excluded, indicate that the clustering threshold setting is a high-risk setting or is not optimal for a given dataset (McCartney-Melstad et al., 2019). During PyRAD assembly of our data leading to our final empirical dataset, we found that on average only ~11 ddRAD tag sequences (mean: 10.8 ddRAD tags, within individuals) were flagged as paralogs and excluded when conducting paralog detection and filtering. Looking at the number of SNP locations flagged as paralogs and excluded from the final dataset when applying paralogy filters across individuals, we found that of 4197 SNPs present before these filters were applied, only 294, or ~7%, of the SNPs in our dataset were excluded by paralog filters. And these were SNPs in the excluded ddRAD tags.

***Population structure and admixture*****—**The inflection point in Bayesian information criterion (BIC) values calculated across a range of $K$ values during DAPC in adegenet corresponded to an initial leveling off of support at $K = 7$ clusters (Figure S1). This value included six ingroup clusters and one outgroup cluster.

***Phylogenomic gene tree and species tree analyses*****—**Other than inferring the genetic structure or history of a sample, networks can be used to check for data-sampling errors or outliers revealed through mislabeling or reticulate patterns; our genome-wide network did not reveal any such data-processing errors. The Neighbor-Net network inferred from the ddRAD-seq dataset was treelike, with the same six ingroup clades (corresponding well overall to DAPC genetic clusters) and one outgroup clade supported by ML and Bayesian analyses above (Figure S5). However, unlike our traditional phylogenies, the network exhibited evidence of incongruences reflecting potential admixture, incomplete lineage sorting, or other sources of homoplasy. For example, putatively admixed individuals identified by fastSTRUCTURE fell out in network cluster 5 (clade 4), which had an intermediate position between the two populations with the highest admixture proportions in Figure 1b, i.e., cluster 2 (clade 3) and cluster 4 (clade 6), and boxes in the network indicated that the greatest number of conflicting splits occurred in the region between clusters 2, 4, and 5 in the network. The network also exhibited conflicts for two polyphyletic samples of genetic cluster 3 (clade 5; HypJCB66 and HypJCB156), which were connected to cluster 5 and cluster 2 (Figure S5), reflecting the same relationships as in our gene trees (Figures 2a, S2, and S4).

***Demographic history and interdrainage migration***—Final chain length and priors yielded good post-burn-in convergence and mixing during G-PhoCS runs, as judged by visualizing parameter traces and ESS scores in Tracer. G-PhoCS runs typically obtained sufficient ESS scores and stationary parameter distributions after the first 50,000 to 600,000 generations. The ESS scores for all parameters were greater than 100–200 in all runs.

Running G-PhoCS with and without estimating migration yielded similar results for $\theta$s and $\tau$s (JCB, unpublished results), and $m_2$ runs inferred some nonzero migration; thus, here and in the main text, we only report parameter estimates from $m_2$ models including low levels of migration (Table S4 and S5). Migration was technically not zero since the 95% HPDs did not include zero, but the migration signals nevertheless were very weak. Overall, inter-lineage migration rates, $m_{sx}$, were small (generally, $< 2.0 \times 10^{-4}$ as raw values; Table S4; even if these were biased downward by an order of magnitude, then the corrected values would still be considered small/weak). After conversions, these values yielded migration proportions ($M_{sx} = m_{sx} \times \theta_x/4$, or migration rate per generation) that were even smaller (Table S5), indicating effectively zero migration in all cases of pairwise migration bands between cluster lineages. Converted $N_e$ values (Table S5) indicated that effective population sizes were inferred by G-PhoCS to be greatest for clusters 3 and 4, but highest for cluster 4, at ~118,000 breeding individuals.

***Ancestral geographic location analyses***— Our BEAST discrete trait diffusion analyses exhibited excellent chain mixing and convergence, with ESS > 200 for all parameters and yielded a single best tree with relationships similar to that of our RAxML gene tree.

***Effects of varying ddRAD-seq assembly parameters on downstream genetic analyses***— Regarding our quantitative sensitivity analyses, we initially wrote a more complex code pipeline in which we used the 'relaimpo' R package (Grömping, 2006) to quantitatively estimate the relative importance of the PyRAD assembly parameters mC, SH, pO, mD, and cP as predictors of outcome variables $d_{RF}$, best $K$, and number of ddRAD tag loci in standard multiple regression models. In the code, discrete variables, and continuous variables identified as non-normally distributed using Shapiro–Wilk tests ($\alpha = 0.05$), were log-transformed prior to relative importance analyses (which can be robust to such violations). However, we realized, based on our findings, that none of the varied parameters had overarching effects on our downstream genetic inferences when considered singly (relative to total variance) or in pairs, even though they did have large effects on practical outcomes such as total output ddRAD tag loci (see Results section, main text). As a result, it did not make sense for us to implement relative importance analyses (or other post-hoc analyses); the corresponding results are not presented.

#### C. Supplementary Discussion

***Testing hypothesized effects of river capture in the central Brazilian Shield***—In the past, various biogeography studies used non-phylogenetic methods, qualitative approaches, or mappings of areas onto phylogenies or area cladograms, or other less rigorous approaches, to infer patterns of dispersal and vicariance of species. Many of these more traditional studies were based on phylogenies reconstructed from small numbers of characters, or were based solely on gene tree topologies for single genes (e.g. mitochondrial or chloroplast DNA markers)—practices generally considered at odds with current paradigms in the field. More recent advances, including improvements in amplicon and genomic sequencing methods, especially those powered by NGS-related developments of the last 15 years, have drastically increased the amount of sequence data (homologous characters) available to infer historical biogeographical processes from natural populations. These breakthroughs have also precipitated a more quantitative basis for hypothesis testing in phylogeography, both in terms of methodological developments for analyzing multilocus or genomic datasets (e.g. Gutenkunst et al., 2009; Gronau et al., 2011; Bouckaert et al., 2014; among many others) and in terms of practical applications to empirical systems (e.g. Solomon et al., 2008; Bagley et al., 2013; Menon et al., 2018; among many others). Our hypothesis testing framework is similar to a priori hypothesis-testing frameworks employed in previous phylogeography studies (e.g. Solomon et al., 2008; Bagley et al., 2013) following in a strong Popperian tradition of rejecting a priori hypotheses. In this style of approach, hypotheses are strictly generated before seeing the data, which requires a foundation of knowledge of the study system or hypotheses beforehand. Another advantage of our study is that we have also relied on molecular sequencing to generate many thousands more characters for analysis than traditional phylogeography approaches (e.g. single-gene, mitochondrial DNA, or traditional morphology-based phylogenetic methods).

In our study, we benefited from a foundation provided by previous studies of the geology and community ecology of freshwater fishes in our study area (e.g. Aquino and Colli, 2017; see section 1 of the main text). On this foundation, we were able to build several predictions of our two main hypotheses, which represent scenarios in which the frequency, directionality, and genetic signatures of past river capture events vary. Discriminating between these hypotheses turned out to be challenging, not necessarily because the hypotheses make predictions that are similar to one another, which for example has posed (partly surmountable) issues for distinguishing between Pleistocene refugia versus marine incursions as drivers of phylogeographic structuring in Amazonian taxa (e.g. Solomon et al., 2008). Rather, the primary issue stemmed from complications that we determined, based on our results, to have most likely arisen due to incomplete sorting of ancestral polymorphism (e.g. reviewed by Funk and Omland, 2003; see discussion below, and in main text sections 3 and 4 for additional details).

The most general interpretation of our results, despite that they were mixed, is that approximately 9/11 (82%) of prediction-by-test comparisons provided positive support for the Paraná Capture Hypothesis, whereas only 2/6 (33%) such comparisons supported the Frequent Interdrainage Dispersal Hypothesis (Figure S8). Therefore, the weight of evidence from the data and results accumulated herein is largely in favor of the Paraná Capture hypothesis, rather than the Frequent Interdrainage Dispersal Hypothesis. Informally (stated as such because we have not actually calculated the probabilities), this means that, whereas both hypotheses may be said to have had equal prior probability, Paraná Capture seems to have greater probability after factoring in our data and results. However, these hypotheses are non-mutually exclusive, meaning that different components of the evidence may support one or both of the hypotheses, and that the 'true' (unknown) evolutionary history of the species/lineage in question may include elements unique or common to either hypothesis.

Regarding our inference that gene tree–species tree incongruence caused by cluster 3 relationships (which were polyphyletic in the gene tree) was likely due to incomplete sorting rather than migration and gene flow, we raise one minor point of note to expand upon our discussion in the main text (section 4.1). Namely, while in theory lineage sorting could have been achieved faster under a scenario involving population expansions (e.g. spatial-demographic expansions generating population structure during and following river capture), our demographic modeling results are more consistent with a history of population size reduction during river captures, reducing a very large ancestral population, and perhaps causing founder effects. This is particularly obvious in our G-PhoCS results (Figure 4a). The SNAPP results show the opposite pattern of large population size or population expansion in cluster 3 (Figure S6). This latter pattern could reasonably be interpreted as consistent with the Paraná Capture Hypothesis; however, if there has been expansion of the cluster 3 population, then we do not see any evidence of complete lineage sorting as a result (or any real effect of this on genealogical sorting levels). It should also be noted that our G-PhoCS and SNAPP analyses yielded estimates of population sizes of each lineage (e.g. demographic changes between lineages), rather than estimates of population size transformation (e.g. demographic expansion within lineages), making any further more nuanced interpretation of demographic history difficult.

Last, we briefly discuss some caveats and limitations of our phylogeographic analyses. One potential weakness of our PhyloMapper results is that they may not reflect sources of uncertainty associated with sequentially inferring the phylogeny and then the parameters of the spatial random walk model, which would require a simultaneous analysis. As discussed in the main text, we overcame this weakness by complementing our PhyloMapper results with a BEAST discrete trait diffusion analysis, in which the tree and parameters of the model were simultaneously estimated in a Bayesian framework; those results are presented in Figures 4 and S7. Nevertheless, we are positive about our PhyloMapper results. The rate-smoothing step of the original PhyloMapper analysis was conducted in 10 replicates, and the parameter optimization step was conducted in 100 replicate analyses with randomized starting locations, to avoid entrapment in local optima and correctly infer the global optimum for the ancestral geographic location (Lemmon and Lemmon, 2008). Indeed, results from 100 independent runs showed that PhyloMapper obtained precise ancestral location estimates (Figure 4b). Moreover, PhyloMapper was still useful to us because it is the only software available (to our knowledge) that performs several spatial analyses (randomization tests) providing direct tests of predictions in Table 1, such as the center test, directionality tests, etc., discussed in the main text.

***Pleistocene phylogeography of* Hypostomus *catfish*—**Regarding our inferences on the demographic history of *Hypostomus* sp. 2, the converted $N_e$ values from G-PhoCS (Table S5) indicated effective population size values that seemed reasonable, or possibly slightly to moderately elevated, based on our experience sampling these fish communities in the field. Of course, the true population sizes of these lineages are unknown. We also did not attempt to validate these numbers in the field (doing so was beyond the scope of our study), and no data are available currently for doing so. Thus, it is difficult to evaluate whether and to what extent these $N_e$ values might have been overestimated. And we would consider additional field-based ecological studies of the demography of catfishes in the genus *Hypostomus* to be a worthwhile goal for future work in the study, for example to estimate abundances or effective sizes through depletion sampling and mark-recapture methods.

The pattern of obtaining the highest $N_e$ estimate for *Hypostomus* cluster 4 in our G-PhoCS results made sense, as this was the cluster from which we had the greatest sampling, and $N_e$ estimation can be sensitive to sample size. The large number of incongruences between this cluster and related clusters in our multilocus phylogenetic network (Figure S5) also seemed consistent with very large ancestral sizes in this lineage, and also earlier in the ancestral population leading to clusters 2–6, as inferred in our G-PhoCS results.

From our SNAPP analysis results, we inferred the largest population size as being that for cluster 3 (Table S6). It is possible that the congruence of relatively higher cluster 3 population size estimates, overall, between SNAPP and G-PhoCS reflects an actual pattern of historically large population sizes in this lineage. However, this was surprising, because cluster 4 had the largest $N_e$ in our G-PhoCS results, and because population size estimates from SNAPP (for this and other clusters) were orders of magnitude higher than those from G-PhoCS. In view of this and our experience working with these fishes in the field, we view the SNAPP converted $N_e$ values ranging in the millions for these lineages, and particularly for cluster 3, as being gross overestimates of actual sizes, and as being more likely reflective of very large ancestral sizes. Our prediction that the actual $N_e$ for cluster 3 is much smaller seems particularly likely to be true in light of the very small geographical ranges of our focal candidate species/lineage and its inferred genetic clusters, including cluster 3 (Figure 1a,c).

Overall, our results show that G-PhoCS and SNAPP can perform differently in estimating $N_e$ and other parameters, and it is unclear which results should be preferred. The G-PhoCS analyses were the first analyses we ran, and our interpretation has been to rely most heavily on them because G-PhoCS is a coalescent-with-migration approach that infers not only population sizes and divergence times, but also migration levels, with good ESS scores. However, recent empirical and simulation-based work has shown that SNAPP is apt to overestimate the divergence time (and possibly also $N_e$) for two lineages when the ancestral population was structured by isolation-by-distance (IBD) or experienced a stepping-stone-like model of connectivity with low migration levels (Hancock and Blackmon, 2020). The $N_e$ estimates from SNAPP and other approaches are also likely to be overestimates when ancestral populations experienced IBD or bottlenecks (which is what in this case the authors found IBD to be mimicking; Hancock and Blackmon, 2020). It remains unclear whether either of these features were present in ancestral *Hypostomus* sp. 2 populations, and our G-PhoCS results certainly cast doubt on the idea that migration levels would be detectable with our data. Still, the low dispersal abilities of *Hypostomus* catfishes and their opportunities for lineage divergence and speciation through vicariance mediated by river captures, e.g. demonstrated by our study and others (e.g. Silva et al., 2016), suggests that the possibility of IBD in the *Hypostomus* sp. 2 ancestral lineage may have been real. Nevertheless, situations in which species have broad geographical distributions are most ideal for generating and hence detecting such patterns as IBD and their influence on divergence times. By contrast, the situation with *Hypostomus* sp. 2 is a very geographically restricted one, making this system less amenable to testing for such IBD effects by comparing results across multiple SNAPP runs or using simulations.

***Effects of varying ddRAD-seq assembly parameters on downstream genetic analyses*** —First, we note that, looking at our empirical assembly, results indicate putative gene-level paralogy of only ~1–2% in our final dataset, and that only ~7% of SNPs in our dataset were in putative paralogous genes. This strongly suggests that paralogy is not a major concern for our dataset; hence, paralogy is unlikely to have had a major influence on our downstream genetic or phylogenetic inferences. We conclude that either paralogy is not a major issue for restriction site-associated loci in our focal taxon, or that our experimental approaches (library preparation and sequencing), data processing, and assembly settings that we used were effective in guarding against paralogy and instead obtaining orthologous loci.

Aside from evaluating our empirical assembly, we only conducted a cursory assessment of paralogy among our other 279 PyRAD assemblies in the test set. Nevertheless, randomly checking 5 assemblies with very different parameter settings indicated similar levels of paralogy to our empirical assembly, around 1–2% of total ddRAD tags. According to other authors, the fraction of loci flagged by PyRAD and Stacks as putatively paralogous can vary widely across RADseq datasets, but ultimately provides insight into the number of loci that were improperly lumped during RADseq data assembly (McCartney-Melstad et al., 2019). The filters in PyRAD and similar software are imperfect and may fail to catch all true paralogs (McCartney-Melstad et al., 2019); however, our results imply that improperly lumping or splitting loci, due to issues related with paralogy, is unlikely to have been substantial in our study.
